# Genomic and metabolic plasticity drive alternative scenarios for adapting *Pseudomonas putida* to non-native substrate D-xylose

**DOI:** 10.1101/2023.05.19.541448

**Authors:** Pavel Dvořák, Barbora Burýšková, Barbora Popelářová, Birgitta Ebert, Tibor Botka, Dalimil Bujdoš, Alberto Sánchez-Pascuala, Hannah Schöttler, Heiko Hayen, Víctor de Lorenzo, Lars M. Blank, Martin Benešík

**Author notes:** Shared first authors (Pavel Dvořák, Barbora Burýšková, Barbora Popelářová, Birgitta Ebert). Current address: Food Science Building, College Rd, University College, Cork, Ireland (Dalimil Bujdoš). Corresponding author: Dr. Pavel Dvořák, Department of Experimental Biology (Section of Microbiology, Microbial Bioengineering Laboratory), Faculty of Science, Masaryk University, Kamenice 735/5, Brno 62500, Czech Republic, https://mik.sci.muni.cz/mbl.

## Abstract

D-Xylose, a major constituent of plant biomass and second most abundant sugar on Earth, holds a considerable potential as a substrate for sustainable bio-production. Pseudomonas putida KT2440 is an attractive bacterial host for valorizing biogenic feedstocks but lacks a xylose utilization pathway. While several attempts to engineer P. putida for growth on xylose have been reported, a comprehensive understanding of xylose metabolism in this bacterium is lacking, hindering its further improvement and rational tailoring for specific biotechnological purposes. In this study, we elucidated the xylose metabolism in the genome-reduced P. putida strain, EM42, endowed with xylose isomerase pathway (xylAB) and transporter (xylE) from Escherichia coli and used the obtained knowledge in combination with adaptive laboratory evolution to accelerate the bacterium’s growth on the pentose sugar. Carbon flux analyses, targeted gene knock-outs, and in vitro enzyme assays portrayed xylose assimilation in P. putida and confirmed a partially cyclic upper xylose metabolism. Deletion of the local transcriptional regulator gene hexR de-repressed genes of several key catabolic enzymes and reduced the lag phase on xylose. Guided by metabolic modeling, we augmented P. putida with additional heterologous pentose phosphate pathway genes and subjected rationally prepared strains to adaptive laboratory evolution (ALE) on xylose. The descendants showed accelerated growth and reduced growth lag. Genomic and proteomic analysis of engineered and evolved mutants revealed the importance of a large genomic re-arrangement, transaldolase overexpression, and balancing gene expression in the synthetic xylABE operon. Importantly, omics analyses found that similar growth characteristics of two superior mutants were achieved through distinct evolutionary paths. This work provides a unique insight into how cell metabolism adjusts to a non-native substrate; it highlights the remarkable genomic and metabolic plasticity of P. putida and demonstrates the power of combining knowledge-driven engineering with ALE in generating desirable microbial phenotypes.

**Highlights:**

- Elucidated xylose catabolism via exogenous isomerase pathway in *P. putida* EM42.
- Deletion of transcriptional regulator HexR improved growth on xylose.
- Knowledge-guided interventions and adaptive evolution accelerated growth.
- Omics analyses of selected mutants highlighted the genomic and metabolic plasticity of *P. putida*.
- Two mutants with superior characteristics emerged from distinct evolutionary paths.

## 1. Introduction

Renewable lignocellulose can be converted into valuable biochemicals in biotechnological processes that leverage natural or recombinant microorganisms as biocatalysts. D-Xylose is a predominant monomeric hemicellulose component, which forms 10 – 50 % of the mass of lignocellulosic residues (Taha et al., 2016). Efficient utilization of xylose and other lignocellulose-derived sugars (e.g., D-glucose, L-arabinose) and aromatic chemicals (e.g., p-coumarate, ferulate) is a key prerequisite for the economic viability of these processes (Narisetty et al., 2021; Valdivia et al., 2016). However, many biotechnologically relevant microorganisms lack xylose metabolism (*Zymomonas mobilis*, *Corynebacterium glutamicum*, *Saccharomyces cerevisiae*), while others can grow on this pentose sugar but prefer glucose or other available organic substrates as their primary carbon and energy source (e.g., *Escherichia coli*, *Bacillus subtilis*) (Görke and Stülke, 2008). In these cases, the desirable phenotype can be established using the tools of metabolic engineering, often followed by adaptive laboratory evolution (ALE) (Sandberg et al., 2019).

Gram-negative bacteria from the genus *Pseudomonas* are sought after in biotechnology, particularly for their robustness, fast growth in inexpensive media, and versatile metabolism (Nikel et al., 2014). While some pseudomonads such as *P. taiwanensis* VLB120 (Köhler et al., 2015), *P. fluorescens* SBW25 (Liu et al., 2015), or *P. putida* W619 (Davis et al., 2013) were reported to grow on xylose, others including solvent-tolerant *P. putida* S12 or *P. putida* KT2440 do not possess the initial catabolic steps for pentose sugars (Belda et al., 2016; Meijnen et al., 2008). *P. putida* KT2440 is one of the most studied pseudomonads, especially in the biodegradation and bioremediation field (Dvořák et al., 2017). Given its capacity to degrade high-molecular weight lignin and to stream the resulting aromatic chemicals into biomass and attractive bioproducts such as polyhydroxyalkanoates or *cis*,*cis*-muconic acid, strain KT2440 is now seen also as a promising cell factory for lignocellulose valorization (Kohlstedt et al., 2018; Ling et al., 2022; Salvachúa et al., 2020). Various derivatives of this bacterium with a reduced genome have been prepared of which strain EM42 (∼4.3% of the genome deleted) was repeatedly shown to possess properties advantageous for harsh biotechnological processes (Dvořák and de Lorenzo, 2018; Jayakody et al., 2018; Kohlstedt et al., 2022; Martínez-García et al., 2014b).

Several research groups intended to overcome *P. putida*’s inability to utilize xylose by implanting known bacterial xylose catabolic routes – the isomerase pathway, Weimberg pathway, and Dahms pathway (Bator et al., 2020; Dvořák and de Lorenzo, 2018; Elmore et al., 2020; Le Meur et al., 2012; Lim et al., 2021; Wang et al., 2019) – following to some extent the pioneering work conducted with *P. putida* S12 (Meijnen et al., 2012, 2009, 2008). The isomerase route from *E. coli* was introduced (Dvořák and de Lorenzo, 2018; Elmore et al., 2020; Le Meur et al., 2012; Wang et al., 2019) and enabled through the action of xylose isomerase XylA and xylulokinase XylB the conversion of xylose to xylulose 5-phosphate (**Figs 1 and 2**), which enters the pentose phosphate pathway (PPP) in so-called EDEMP cycle typical for *P. putida* (Nikel et al., 2015). In our former study (Dvořák and de Lorenzo, 2018), we found that *P. putida* EM42 requires the XylE xylose/H^+^ symporter from *E. coli* and the deletion of the *gcd* gene encoding periplasmic membrane-bound glucose dehydrogenase (PP_1444) to fully utilize xylose (**Fig. 1**). This is consistent with the findings of Meijnen et al. (Meijnen et al., 2012, 2008) in *P. putida* S12 and Elmore and co-workers (Elmore et al., 2020) in *P. putida* KT2440. We also demonstrated that the recombinant *P. putida* EM42 can co-utilize xylose with glucose or cellobiose and co-valorize these sugars (Dvořák et al., 2020; Dvořák and de Lorenzo, 2018). Elmore et al. (2020) expanded this idea further and prepared a *P. putida* KT2440 strain capable of simultaneously processing five major components of corn stover hydrolysates: D-glucose, D-xylose, L-arabinose, *p*-coumarate, and acetate (Elmore et al., 2020). Metabolic analyses demonstrated stoichiometric superiority of the isomerase pathway over alternative xylose catabolic routes for the production of a number of valuable biochemicals, for which the isomerase pathway showed the highest product yields (Bator et al., 2020). *P. putida*, empowered with an efficient xylose metabolism based on the exogenous isomerase route, thus represents a very attractive biocatalyst to produce biosurfactants, dicarboxylic acids, sorbitol, acetoin, some amino acids (e.g., threonine), and potentially many other compounds that can be synthesized from xylose alone or from complex lignocellulosic hydrolysates (Bator et al., 2020; Elmore et al., 2020; Ling et al., 2022). However, while the xylose oxidative metabolism in KT2440 was partially mapped by the analysis of proteinogenic amino acids (Bator et al., 2020) or transcriptomics (Lim et al., 2021), the xylose assimilation in the bacterium’s biochemical network extended with the isomerase route remains unresolved. This complicates further attempts to tailor the utilization and valorization of this sugar in *P. putida* on a rational basis.

**Figure 1.**
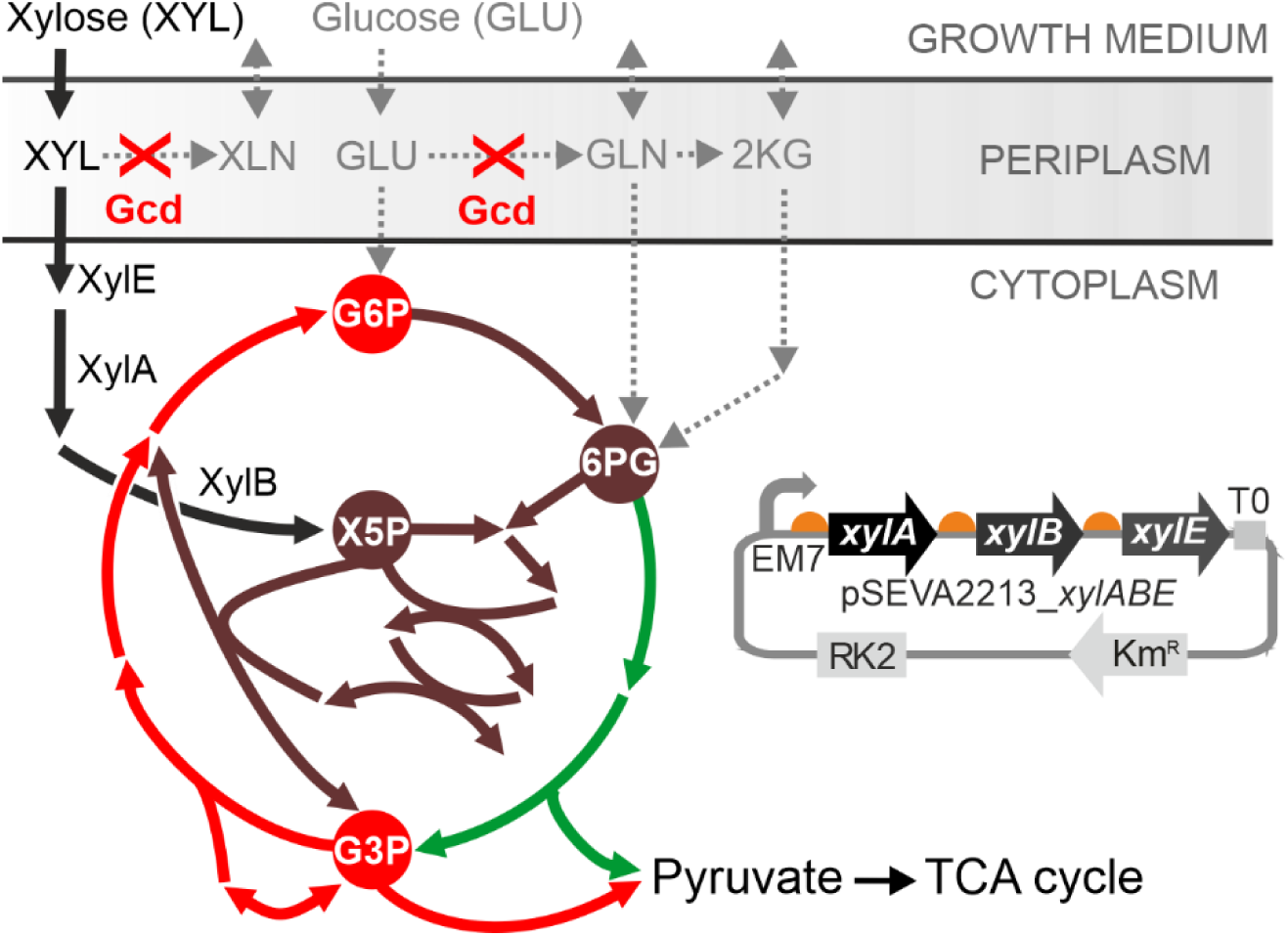
Schematic illustration of upper sugar metabolism of *Pseudomonas putida* EM42 PD310 used as a template strain in this study. The PD310 strain (Dvořák and de Lorenzo, 2018) capable of growth on D-xylose and its co-utilization with D-glucose bears low-copy-number plasmid pSEVA2213 with constitutive P_EM7_ promoter and a synthetic operon that encodes exogenous xylose isomerase pathway (XylA xylose isomerase and XylB xylulokinase) and xylose/H^+^ symporter XylE from *Escherichia coli* BL21(DE3). The *xylA* and *xylE* genes are preceded by synthetic ribosome binding sites (RBS, orange hemispheres), while *xylB* was left with its native RBS. Note that the elements in the plasmid scheme are not drawn to scale. The PD310 strain was also deprived of the *gcd* gene, which encodes periplasmic glucose dehydrogenase (PP_1444) to prevent the transformation of xylose to the dead-end product xylonate (XLN). The exogenous pathway converts xylose to xylulose 5-phosphate (X5P), which enters the EDEMP cycle formed by the reactions of the pentose phosphate pathway (brown arrows), the Embden-Meyerhof-Parnas pathway (red arrows), and the Entner-Doudoroff pathway (green arrows). Abbreviations: GLN, gluconate; G6P, glucose 6-phosphate; G3P, glyceraldehyde 3-phosphate; Km, kanamycin; 2KG, 2-ketogluconate; 6PG, 6-phosphogluconate; RK2, a broad-host-range origin of replication; T0, transcriptional terminator.

**Figure 2.**
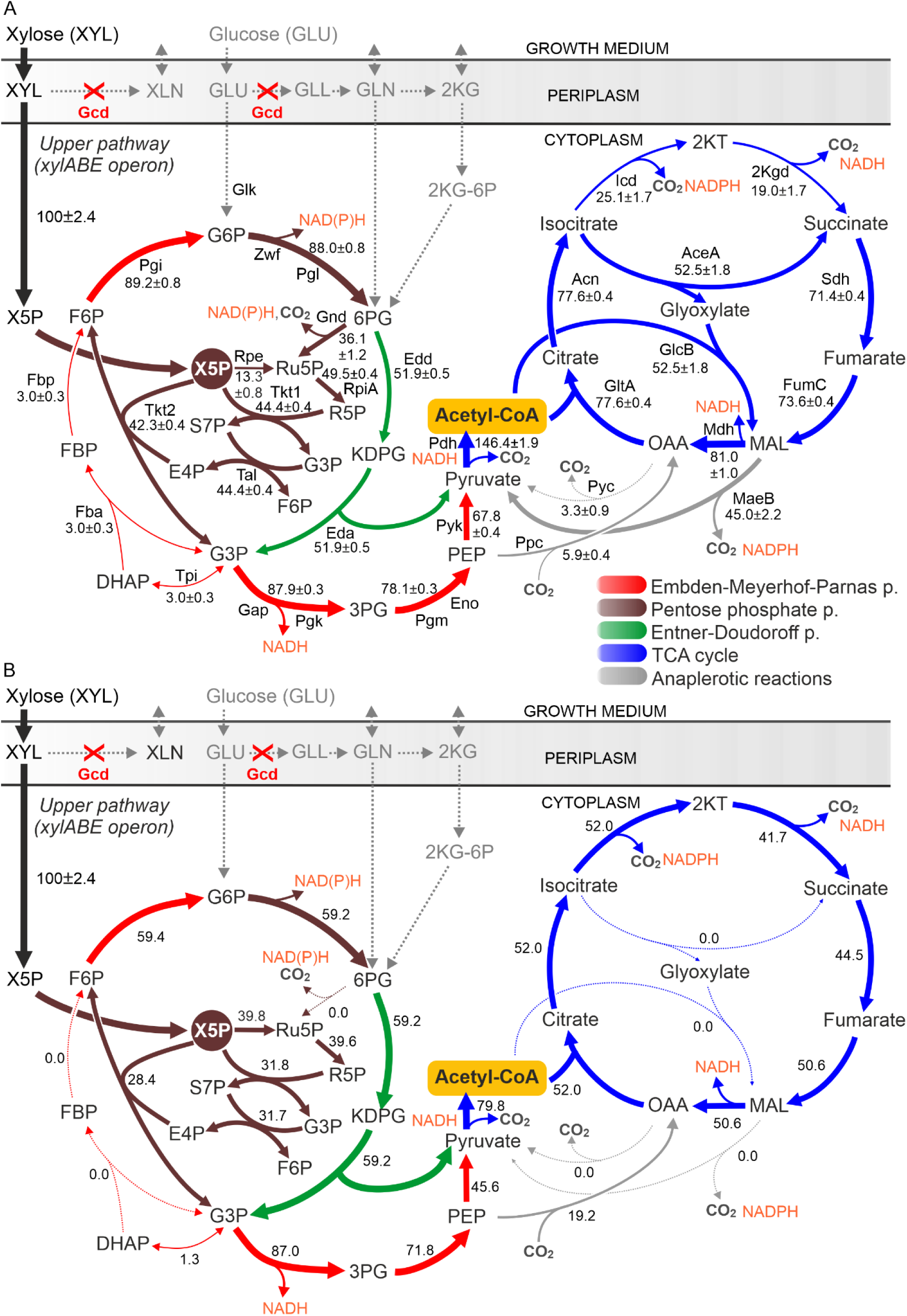
In vivo (**A**) and in silico (**B**) distribution of carbon fluxes in engineered strain *P. putida* EM42 PD310 grown on xylose as determined by ^13^C metabolic flux analysis and flux balance analysis, respectively. Fluxes are given as a molar percentage of the mean specific xylose uptake rate *q*_S_ = 1.45 mmol g_CDW_^−1^ h^−1^, which was set to 100 %. Arrow thickness roughly corresponds with the given flux value. Experimental flux values (**A**) represent the mean ± STD from two biological replicates. Abbreviations (enzymes): AceA, isocitrate lyase; Acn, aconitate hydratase; Eda, 2-keto-3-deoxy-6-phosphogluconate aldolase; Edd, 6-phosphogluconate dehydratase; Eno, phosphopyruvate hydratase; Fba, fructose-1,6– biphosphate aldolase; Fbp, fructose-1,6-biphosphatase; FumC, fumarate hydratase; Gap, glyceraldehyde-3-phosphate dehydrogenase; Gcd, glucose dehydrogenase; Gnd, 6-phosphogluconate dehydrogenase; GlcB, malate synthase; GltA, citrate synthase; Icd, isocitrate dehydrogenase; 2Kgd, 2– ketoglutarate dehydrogenase; MaeB, malic enzyme; Mdh, malate dehydrogenase; Pdh, pyruvate dehydrogenase; Pgi, glucose-6-phosphate isomerase; Pgk, phosphoglycerate kinase; Pgl, 6– phosphogluconolactonase; Pgm, phosphoglycerate mutase; Ppc, phosphoenolpyruvate carboxylase; Pyc, pyruvate carboxylase; Pyk, pyruvate kinase; Rpe, ribulose-5-phosphate 3-epimerase; RpiA, ribose– 5-phosphate isomerase; Sdh, succinate dehydrogenase; Tal, transaldolase; Tkt, transketolase; Tpi, triosephosphate isomerase; Zwf glucose-6-phosphate dehydrogenase. Abbreviations (metabolites): DHPA, dihydroxyacetone phosphate; E4P, erythrose 4-phosphate; FBP, fructose 1,6-biphosphate; F6P, fructose 6-phosphate; GLL, glucono-δ-lactone; GLN, gluconate; G3P glyceraldehyde 3-phosphate; G6P, glucose 6-phosphate; KDPG, 2-keto-3-deoxy-6-phosphogluconate; 2KG, 2-ketogluconate; 2KG-6P, 2– ketogluconate 6-phosphate; 2KT, α-ketoglutarate; MAL, malate; NADH, reduced nicotinamide adenine dinucleotide; NADPH, reduced nicotinamide adenine dinucleotide phosphate; OAA, oxaloacetate; PEP, phosphoenolpyruvate; 3PG, 3-phosphoglycerate; 6PG, 6-phosphogluconate; R5P, ribose 5-phosphate; Ru5P, ribulose 5-phosphate; S7P, seduheptulose 7-phosphate; XLN, xylonate; X5P, xylulose 5– phosphate. Individual enzyme reactions are described, and abbreviations explained in **Supplementary Files S1 and S2** and in the manuscript text.

In the present study, we used a knowledge-driven engineering approach to (i) explore how the metabolism of *P. putida*, natively evolved towards processing primarily organic acids, amino acids, or aromatic compounds (Belda et al., 2016; Rojo, 2010), handles carbon fluxes from non-native sugar substrate, and to (ii) boost xylose utilization in our previously reported *P. putida* EM42 strain with an exogenous xylose isomerase pathway (**Fig. 1**, **Table 1**). Carbon flux and enzyme activity analyses helped us draw the initial picture of xylose metabolism in *P. putida*, remove a bottleneck on the level of regulation of the EDEMP cycle enzymes, and identify other potentially problematic nodes in PPP. Following ALE of strains with and without rational interventions in the PPP gave rise to new mutants with a doubled growth rate and up to 5-fold reduced lag phase. Genomic and proteomic analyses of mutant strains demonstrated that *P. putida* can accelerate its growth on xylose even without additional exogenous genes but can adopt them when pushed to do so through rational engineering cuts. The bacterium takes advantage of its genomic and metabolic plasticity, which helps the organism in searching for alternative solutions that lead to an improved phenotype. The information presented here is pivotal to further harnessing xylose metabolism in *P. putida* and contributes to a better understanding of the adaptation of bacterial metabolism to a non-native substrate.

**Table 1.** Bacterial strains used in this study.

| Strain or plasmid | Characteristics | Source or reference |
| --- | --- | --- |
| ***Escherichia coli*** |  |  |
| Dh5α | Cloning host: F-λ-*endA1 glnX44(AS) thiE1 recA1 relA1 spoT1 gyrA96(Nal<sup>R</sup>) rfbC1 deoR nupG Φ80(lacZΔM15) Δ(argF-lac)U169 hsdR17(r<sub>K</sub>-m<sub>K</sub><sup>+</sup>)* | (Grant et al., 1990) |
| CC118 | Cloning host: Δ(*ara-leu*) *araD* Δ*lac* X174 *galE galK phoA thiE1 rpoB(Rif<sup>R</sup>) argE(Am) recA1* | (Manoil and Beckwith, 1985) |
| CC118λpir | Cloning host: *araD139* Δ(*ara-leu*)7697 Δ*lac*X74 *galE galK phoA20 thi-1 rpsE rpoB(Rif<sup>R</sup>) argE(Am) recA1*, λpir lysogen | (Herrero et al., 1990) |
| HB101 | Helper strain for tri-parental mating: F- λ- *hsdS20*(r<sub>B</sub><sup>-</sup> m<sub>B</sub><sup>-</sup>) *recA13 leuB6(Am) araC14* Δ(*gpt-proA*)62 *lacY1 galK2(OC) xyl-5 mtl-1 thiE1 rpsL20*(Sm<sup>R</sup>) *glnX44(AS)* | (Boyer and Roulland-Dussoix, 1969) |
| ***Pseudomonas putida*** |  |  |
| EM42 | Genome-reduced derivative of strain KT2440: Δprophages1,2,3,4 ΔTn7 Δ*endA1* Δ*endA2* Δ*hsdRMS* Δ*flagellum* ΔTn4652 | (Martínez-García et al., 2014b) |
| EM42 Δ*gcd* | EM42 with scarless deletion of *gcd* gene (PP_1444) encoding periplasmic glucose dehydrogenase | (Dvořák and de Lorenzo, 2018) |
| EM42 Δ*gcd* Δ*gnd* | EM42 with scarless deletion of *gcd* and *gnd* gene (PP_4043) encoding 6-phosphogluconate dehydrogenase | This study |
| EM42 Δ*gcd* Δ*pgi-I+II* | EM42 with scarless deletion of *gcd*, *pgiI* and *pgiII* genes (PP_1808 and PP_4701) encoding glucose 6-phosphate isomerase | This study |
| EM42 Δ*gcd* Δ*edd* | EM42 with scarless deletion of *gcd* and *edd* gene (PP_1010) encoding phosphogluconate dehydratase | This study |
| EM42 Δ*hexR* | EM42 with scarless deletion of *hexR* gene (PP_1021) encoding DNA-binding transcriptional regulator | This study |
| PD310 | EM42 Δ*gcd* with plasmid pSEVA2213_*xyIABE* bearing synthetic *xyIABE* operon encoding XylA xylose isomerase, XylB xylulokinase, XylE xylose-H<sup>+</sup> symporter from *E. coli* (*EcoRI/HindIII*), ~118 kbp multiplication (PP_2114 - PP_2219) in the chromosome, see plasmid characteristics in Supplementary Table S1 | (Dvořák and de Lorenzo, 2018) |
| EM42 Δ*gcd* pSEVA2213_*xyIABE* | EM42 Δ*gcd* transformed with pSEVA2213_*xyIABE* plasmid (Table S1 in Supplementary material) | This study |
| PD584 | EM42 Δ*gcd* Δ*hexR* with pSEVA2213_*xyIABE*, ~118 kbp multiplication (PP_2114 - PP_2219) in chromosome | This study |
| PD584 L3 | Derivative of PD584 obtained after adaptive laboratory evolution (ALE) on xylose, ~118 kbp multiplication (PP_2114 - PP_2219) in chromosome | This study |
| PD584 tt L3 | Derivative of PD584 with chromosomally integrated expression cassette ($P_{EM7}$-*talB*-*tktA*-$Sm^R$ bearing genes for transaldolase and transketolase from *E. coli*) obtained after ALE on xylose, ~118 kbp multiplication (PP_2114 - PP_2219) in chromosome | This study |
| PD584 ttrr L2 | Derivative of PD584 with chromosomally integrated expression cassette ($P_{EM7}$-*talB*-*tktA*-*rpe*-*rpiA*-$Sm^R$ bearing genes for transaldolase, transketolase, D-ribulose-5-phosphate 3-epimerase, and D-ribose-5-phosphate isomerase from *E. coli*) obtained after ALE on xylose, ~118 kbp multiplication (PP_2114 - PP_2219) in chromosome | This study |
| PD689 | EM42 $\Delta gcd \Delta hexR \Delta gnd$ with pSEVA2213_*xyIA**BE* | This study |
| PD689 tt L1 | Derivative of PD689 with chromosomally integrated expression cassette ($P_{EM7}$-*talB*-*tktA*-$Sm^R$ bearing genes for transaldolase and transketolase from *E. coli*) obtained after ALE on xylose | This study |

Abbreviations: $P_{EM7}$ , constitutive promoter EM7; Sm, streptomycin.

## 2. Materials and methods

### 2.1 Bacterial strains and conditions of routine cultures

Bacterial strains used in this study are listed in **Table 1**. *Escherichia coli* strains were used for cloning or triparental mating. They were routinely grown in lysogeny broth (LB; 10 g L^-1^ tryptone, 5 g L^-1^ yeast extract, 5 g L^-1^ NaCl) with agitation (300 rpm, Heidolph Unimax 1010 and Heidolph Incubator 1000; Heidolph Instruments) at 37°C. Chloramphenicol (Cm, 30 μg mL^-1^) was supplemented to the medium with *E. coli* helper strain HB101. Other antibiotics – kanamycin (Km, 50 µg mL^-1^), ampicillin (Amp, 150 or 450 µg mL^-1^ in *E. coli* and *P. putida* cultures, respectively), streptomycin (Sm, 50 or 60 µg mL^-1^ in *E. coli* and *P. putida* cultures, respectively), or gentamicin (Gm, 10 µg mL^-1^) – were added to liquid or solid media for plasmid maintenance and selection. *P. putida* strains were routinely pre-cultured overnight (16 h) in 15 mL falcon tubes with 2.5 mL of LB medium with agitation (300 rpm, Heidolph Unimax 1010 and Heidolph Incubator 1000; Heidolph Instruments) at 30°C. All precultures were inoculated directly from cryogenic glycerol stocks stored at –70 °C and prepared from single isolated clones. For the main cultures in 250 mL Erlenmeyer flasks or 48-well microtiter plates, overnight cultures were spun (2,000 g, room temperature RT, 7 min) and washed with M9 mineral salt medium (7 g L^-1^ Na_2_HPO_4_·7H_2_O, 3 g L^-1^ KH_2_PO_4_, 0.5 g L^-1^ NaCl, 1 g L^-1^ NH_4_Cl_2_, 2 mM MgSO_4_, 100 µM CaCl_2_, 20 µM FeSO_4_) supplemented with 2.5 mL L^-1^ trace element solution (Abril et al., 1989). Cells were then resuspended to a starting OD_600_ of 0.1 in 50 mL of M9 medium with Km in case of shake flask cultures or to an initial OD_600_ of 0.05 (as measured in a cuvette with an optical path of 1 cm) in 600 μL of M9 medium with Km in case of 48-well plate cultures. A carbon source (D-xylose, D-glucose, or D-fructose) was added in a concentration defined in text or respective figure caption. Flasks cultures were incubated at 30 °C with agitation (200 rpm) using IS-971R incubated shaker (Jeio Tech) and growth was monitored by measuring the OD_600_ of cultures using UV/VIS spectrophotometer Genesys 5 (Spectronic). Microtiter plates were placed in an Infinite M Plex plate reader (Tecan) and incubated at 30° C. The OD_600_ was measured every 15 minutes and orbital shaking with 2.5 μm amplitude was applied in between measurements, linear shaking with 2.5 μm amplitude was set for 10 s before each OD measurement. To avoid condensation of water vapor on the plate lid, the inner surface was treated with a detergent solution in ethanol (0.05% v/v Triton X-100 in 20% v/v EtOH). Excess liquid was decanted and the lid was dried and UV sterilized. All solid media used (LB and M9) contained 15 g L^-1^ agar. M9 solid media were supplemented with 2 g L^-1^ xylose and 50 µg mL^-1^ Km.

### 2.2 General cloning procedures, construction of mutant strains and plasmids

General cloning procedures are described in **Supplementary methods** section in Supplementary material. All plasmids used or prepared in this study are listed in **Table S1** in Supplementary material. Oligonucleotide primers used in this study (**Table S2** in Supplementary material) were purchased from Merck.

#### Preparation of deletion mutants of P. putida EM42

Deletion mutants EM42 Δ*gcd* Δ*gnd*, EM42 Δ*gcd* Δ*pgi*-I+II, EM42 Δ*gcd* Δ*edd*, EM42 Δ*hexR*, EM42 Δ*gcd* Δ*hexR*, and EM42 Δ*gcd* Δh*exR* Δ*gnd* were prepared using the homologous recombination-based protocol described previously (Dvořák and de Lorenzo, 2018; Martínez-García and de Lorenzo, 2012). Briefly, the regions of approximately 500 bp upstream and downstream of the *gnd* (PP_4043), *pgi*-I+II (PP_1808 and PP_4701), *edd* (PP_1010), and *hexR* (PP_1021) genes were PCR amplified with respective TS1F, TS1R (upstream) and TS2F, TS2R (downstream) primers (**Table S2**). TS1 and TS2 fragments were joined through overlap extension or SOEing-PCR (Horton et al., 1990), and the PCR product was digested with *Eco*RI and *Bam*HI and cloned into a non-replicative pEMG plasmid. The resulting pEMG constructs (**Table S1**) were individually propagated in *E. coli* CC118λpir cells and the whole TS1-TS2 region in each of the constructs was sequenced in several clones selected based on the results of colony PCR with respective TS1F and TS2R primers (product of about 1 kb expected). The sequence-verified plasmid was transformed into competent *P. putida* EM42 cells by electroporation. Transformants were selected on LB agar plates with Km and several co-integrates were pooled for further work. The pSW-I plasmid was transformed into co-integrates by electroporation. Transformants were plated on LB agar plates with Amp and expression of I-*Sce*I in selected clones inoculated into 5 mL of LB was induced with 5 mM 3-methylbenzoate (3MB) for 6 — 16 h depending on the deletion. Induced cells were plated on LB agar plates with and without Km and EM42 clones sensitive to Km were checked for the target deletion by colony PCR using respective TS1F/TS2R or check fw/check rv (anneal within the deleted gene) primer pair (**Table S2**). *P. putida* recombinants were cured of pSW-I plasmid after several passes in LB medium lacking Amp.

#### Construction of plasmids pBAMD1-4_EM7_talB-tktA and pBAMD1-4_EM7_talB-tktA-rpe-rpiA

The genes *talB* for transaldolase B (JW0007) and *tktA* for transketolase A (JW5478) were PCR amplified with their native ribosome binding sites (RBS) from *E. coli* BL21(DE3) genomic DNA using Q5 polymerase (NEB). In the second PCR step, an overhang homologous to the 5’ end of *tktA* was added to the 3’ end of *talB* to allow following SOEing-PCR and connecting the two genes in a synthetic operon. The used primers (talB_fw, talB_PCR1_rv, talB_PCR2_rv, tktA_fw, and tktA_rv) are listed in **Table S2**. The construct was digested with *Sac*I and inserted into the mini-Tn5 pBAMD1-4 vector cargo site (Martínez-García et al., 2014a) downstream of the previously added constitutive P_EM7_ promoter. Due to the use of a single restriction site (*Sac*I), the vector was dephosphorylated by the addition of 2 U of shrimp alkaline phosphatase (New England BioLabs) for the last 30 min of the restriction reaction. The genes *rpe* for D-ribulose-5-phosphate 3-epimerase (JW3349) and *rpiA* for ribose 5-phosphate isomerase A (JW5475) were codon optimized for *P. putida* KT2440 and complemented with synthetic RBS designed by RBS Calculator (Salis et al., 2009) to resemble the strengths of native *rpe* and *rpiA* RBS in the context of pBAMD1-4 expression cassette. The whole *rpe*-*rpiA* cassette, flanked with *Sph*I and *Not*I restriction sites, was synthesized by Eurofins Genomics. The cassette was subcloned from the delivery plasmid pEX into pBAMD1-4_EM7 or downstream of the *talB*-*tktA* genes in pBAMD1-4_EM7_*talB*-*tktA*. Chemocompetent *E. coli* CC118λπ cells or OneShot PIR1 *E. coli* cells (Thermo Fisher Scientific) were transformed with the plasmids. Cells were plated on agar plates with Sm (50 µg mL^-1^) and grown overnight at 37 °C. The presence of plasmid with an insert of the correct size was checked in individual clones by colony PCR and by restriction analysis of isolated plasmids. The sequences of all cloned genes were confirmed by Sanger sequencing and sequence alignments were carried out in Benchling (Alignment function MAFFT v7).

2.3 Genomic integrations of *talB*-*tktA* and *talB*-*tktA*-*rpe*-*rpiA* expression cassettes in *P. putida* EM42 Δ*gcd* Δ*hexR* and *P. putida* EM42 Δ*gcd* Δ*hexR* Δ*gnd* and subsequent adaptive laboratory evolution

The previously published procedure for genomic integration using the pBAMD vector system (Martínez-García et al., 2014a) was followed with some modifications. Both *P. putida* EM42 recipient strains (double deletion mutant Δ*gcd* Δ*hexR* and triple deletion mutant Δ*gcd* Δ*hexR* Δ*gnd* both with pSEVA2213_*xylABE* plasmid) were electroporated with the pBAMD1-4_EM7_*talB*-*tktA* or pBAMD1-4_EM7_*talB*-*tktA*-*rpe*-*rpiA* plasmid (100 ng). For controls, the recipient strains were electroporated with 1 µL of pure miliQ water instead of the plasmid. After the electroporation pulse, cells were immediately resuspended in 2.5 ml of SOC medium in a 15 mL falcon tube, and after a 5 h recovery period at 30 °C and 200 rpm, 10 µL were plated on LB agar with Sm to assess the integration efficiency. The rest of the suspension was inoculated into 20 mL of M9 medium supplemented with xylose (5 g L^-1^), Km, and Sm (only Km was used for the controls) in a 100 mL Erlenmeyer flask and incubated at 30 °C and 200 rpm (IS-971R, Jeio Tech) overnight. The next day, the cultures were inoculated into fresh medium to an OD_600_ of 0.2 and incubated for 4 – 5 days (time interval was longer in case of PD689 derivatives due to their negligible growth during the first four days). Then, the cells were transferred to fresh medium with xylose (5 g L^-1^) and a single antibiotic Km to a starting OD_600_ of 0.1 and passaged every 48 or every 24 h for 14 days in total (**Fig. 5**). The 48-h interval was used for the first two passages of PD584 transformants and the PD689 control because these cultures initially showed slower growth than the cultures of PD689 transformants. By adopting this strategy, we aimed to select for cells that grew faster and reached a higher biomass yield on xylose than the template strains (Meijnen et al., 2008). Twice during the experiment, glycerol stocks of the evolved cultures were prepared and individual clones were isolated. Firstly, when the OD_600_ of a given culture reached the value of ≥3.5 within 24 h of growth, and secondly at the end of the experiment (after ∼60–70 generations, **Fig. 5**). Cells were plated on M9 agar plates with 2 g L^-1^ xylose and Km and 2–3 fastest growing clones from each cultivation were picked for further characterization. Growth of the selected clones on xylose was first tested in a 48-well plate format and the nine fastest-growing candidates were verified in 24 h-long shake-flask cultures (**Supplementary Fig. S6**). The number of generations in the evolution experiment was calculated for each of the evolved strains using the equation (1): (Ling 2022 NatComm)

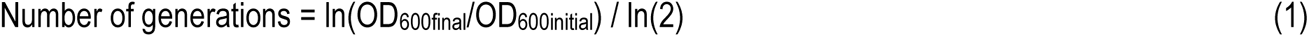

### 2.4 Calculations of dry cell weight and growth parameters

For biomass yield (Y_X/S_) calculations, biomass was determined as dry cell weight (DCW). Based on the previously prepared standard curve, one OD_600_ unit determined in *P. putida* EM42 culture in M9 medium is equivalent to 0.38 g L^-1^ of DCW (Bujdoš et al., 2023). Important culture parameters were determined as described by Long and Antoniewicz (Long and Antoniewicz, 2014). Specific growth rate (μ) in shake flask experiments was determined during exponential growth by plotting the natural logarithm of biomass concentration versus time and quantifying the slope from regression analysis. Biomass yield (Y_X/S_, where X represents biomass and S represents substrate) was calculated by plotting biomass concentration versus substrate concentration and quantifying the slope from regression analysis. Biomass-specific substrate uptake rate (*q*_s_) was determined during exponential growth using equation (2):

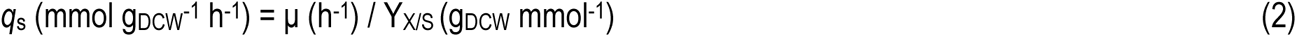

Maximal specific growth rate (μmax) and lag phase (in h) were calculated for cultures grown in 48-well microtiter plates using the default settings of The deODorizer program (Swain et al., 2016).

### 2.5 ^13^C labeling experiments and analysis of metabolic fluxes

An initial pre-culture in LB medium was inoculated from a cryogenic glycerol stock of strain PD310 (**Table 1**) and rotary shakers at 30°C, 200 rpm and 5 cm amplitude. The second preculture was performed in 100 mL shake flask using 10 mL M9 medium with 5 g L^-1^ xylose. The medium was inoculated with the first preculture to an OD_600_ of 0.05 and incubated overnight under otherwise identical conditions as the LB preculture. Three 250 mL Erlenmeyer flasks containing 25 mL M9 medium were inoculated with the second overnight culture to an OD_600_ of 0.025. The media contained 5 g L^-1^ 1,2-^13^C xylose, i.e., xylose labeled at positions C1 and C2 (Sigma-Aldrich, 99% purity). Cultures were grown under agitation (300 rpm, amplitude: 5 cm) at 30° C. Substrate uptake was monitored by HPLC-UV/RI analysis of the fermentation broth. Biomass concentrations were monitored using the cell growth quantifier (Scientific Bioprocessing) and determined manually by measuring the optical density at 600 nm with an Ultrospec 10 Cell Density Meter (GE Healthcare). The conversion factor of OD_600_ to cell dry weight (CDW) in g/L was gravimetrically determined to 0.39.

During the exponential phase, OD and xylose concentrations were determined in 1 h intervals to allow accurate determination of growth and substrate uptake rates. At mid-exponential growth (OD_600_ of 1.0–1.3), samples were taken to quantify the ^13^C isotope incorporation into proteinogenic amino acids and free intracellular metabolites. A fast filtration method (Bennette et al., 2011) was applied for intracellular metabolites to rapidly sample the biomass and quench the metabolism. Briefly, samples corresponding to 10 mg biomass were taken and the biomass was collected by vacuum filtration (Durapore, PVDF, 0.45 µm, 47 mm Sigma-Aldrich). After one washing step with 0.9 % saline, the filter was placed (upside down) into a small Petry dish (5 cm diameter) filled with methanol (pre-cooled at –80 °C), incubated at –80 °C for 1 h. Filters were scraped and rinsed to ensure all cell material was in the liquid solvent, the solvent and filter were transferred to microfuge tubes and vigorously vortexed at –20 °C. The filter was removed, the extract centrifuged at 17,000 g at 4 °C for 5 min, and the supernatant removed and stored at −80 °C. Filter and solvent were transferred into a tube and vortexed under cooled conditions; Extracts were dried in a lyophilizer and analysed using capillary IC-MS analysis (**Supplementary methods**). The metabolites from upper glycolysis were important for a better resolution of fluxes of this part of the metabolism, especially cyclic fluxes. Capillary IC-MS data were corrected for the natural abundance of heavy isotopes using IsoCorr (Millard et al., 2012).

### 2.6 Determination of ^13^C-labeling patterns in proteinogenic amino acids

A standard protocol was used as described in Schmitz *et al*. (Schmitz et al., 2017). Samples corresponding to 0.3 mg CDW were taken and centrifuged for 10 min at 4 °C at 17,000 x g. The pellets were washed and resuspended in 5 N HCl, transferred to GC vials that were carefully closed, and the samples were hydrolyzed at 105 °C for 6 h. The hydrolysates were dried at 95 °C and derivatized with tert-butyl dimethyl silane. GC-MS analysis of proteinogenic amino acids was performed as described in Schmitz *et al*. (Schmitz et al., 2017). Data were corrected for unlabeled biomass (introduced with the inoculum) and natural abundance of heavy isotopes using the software iMS2FLUX (Poskar et al., 2012). Metabolic flux analysis was performed using the Matlab-based tool INCA (Young, 2014). The model was constrained with mass isotopomer distribution data of intracellular metabolites and proteinogenic amino acids and the specific growth and xylose uptake rates. Confidence intervals were determined by parameter continuation using INCA’s in-build function.

### 2.7 Genome-scale metabolic model simulations

The most recent iJN1463 genome-wide metabolic reconstruction of *P. putida* KT2440 (Nogales et al., 2020) was downloaded from the BIGG database (http://bigg.ucsd.edu/). Two metabolites (xylose and xylulose) and four reactions – xylose transport to the periplasm and to cytosol (XYLtex, XYLt2pp respectively), xylose isomerase (XYLI1) and xylulokinase (XYLK) were added to the model. The glucose dehydrogenase reaction (GCD) was deleted from the model to prevent the oxidation of glucose to gluconate and 2-ketogluconate. To predict optimal flux distributions by flux balance analysis (FBA) (Orth et al., 2010) either COBRApy library (Ebrahim et al., 2013) or MATLAB COBRA toolbox (Heirendt et al., 2019) was used. Xylose uptake rate was fixed at experimentally determined 1.45 mmol g_CDW_^-1^ h^-1^. The model was then constrained with upper and lower bounds calculated from metabolic flux analysis (**Supplementary File S1**). Only reactions in the central carbon metabolism, TCA cycle, and CO_2_ production were constrained with MFA data. To get an upper and lower bound from MFA data, firstly, the standard error was calculated from two independent measurements and then the lower and upper bound on the fluxes were calculated as ± 1.96 x standard error (1.96 corresponds to the 97.5^th^ percentile of a standard normal distribution). iJN1463 contains two malate dehydrogenase reactions (MDH and MDH2). The combined flux through these two reactions was set to the upper and lower bound calculated from MFA. The two models – one constrained with MFA data (MFA model) and another one constrained only with xylose uptake (FBA model) – were then compared (both modified models are available at GitHub and will be made public with published manuscript). A for loop was set to test the scaling of flux distribution obtained from the MFA model against the optimal flux distribution obtained by the FBA model (we wanted to find out if the MFA flux distribution is scalable). In every step of the loop, the experimentally obtained upper and lower bounds of the MFA model were multiplied by coefficient 1 + *i* x 0.05 where *i* is the number of steps already taken in the loop. In every step of the loop, the initial xylose uptake rate (1.45 mmol g_CDW_^-1^ h^-1^) was multiplied by the same coefficient 1 + *i* x 0.05 and the FBA model was simulated with this xylose uptake rate. In total, 100 steps were simulated for each model. The glpk solver was used for all simulations. The biomass function was always the objective function for these simulations.

### 2.8 Analytical methods

The optical density in cell cultures was recorded at 600 nm using UV/VIS spectrophotometer Genesys 5 (Spectronic). Analytes from cultures were collected by withdrawing 0.5 ml of culture medium. The sample was then centrifuged (20,000 g, 10 min). The supernatant was filtered through 4 mm / 0.45 µm LUT Syringe Filters (Labstore) and stored at –20 °C. Prior to the HPLC analysis, 50 mM H_2_SO_4_ in degassed miliQ water was added to the samples in a 1:1 ratio to stop any hydrolytic activity and to dilute the samples. High-performance liquid chromatography (HPLC) was used to quantify xylose and glucose. HPLC analysis was carried out using Agilent 1100 Series system (Agilent Technologies) equipped with a refractive index detector and Hi-Plex H, 7.7 x 300 mm, 8 µm HPLC column (Agilent Technologies). Analyses were performed using the following conditions: mobile phase 5 mM H_2_SO_4_, mobile phase flow 0.5 mL min^-1^, injection volume 20 µL, column temperature 65 °C, RI detector temperature 55 °C. Xylose and glucose standards (Sigma-Aldrich) were used for the preparation of calibration curves. Xylose concentrations in labeling experiments were determined using a Beckman System Gold 126 Solvent Module equipped with a System Gold 166 UV-detector (Beckman Coulter) and a Smartline RI detector 2300 (Knauer). Analytes were separated on the organic resin column Metab AAC (Isera) eluted with 5 mM H_2_SO_4_ at an isocratic flow of 0.6 mL min^-1^ at 40 °C for 40 min.

Glucose and xylose concentrations in culture supernatants were alternatively determined also by Glucose (GO) Assay Kit (Sigma-Aldrich, USA) and Xylose Assay Kit (Megazyme, Ireland), following the manufactureŕs instructions. Product concentrations were measured spectrophotometrically using Infinite M Plex reader (Tecan). Xylonate was measured using the hydroxamate method (Lien, 1959) and the protocol described previously (Dvořák et al., 2020). Absorbance at 550 nm was measured by Infinite M Plex reader (Tecan). Xylonate concentrations were quantified with a standard curve prepared using the pure compound (Sigma-Aldrich).

### 2.9 Data and statistical analyses

The number of repeated experiments or biological replicates is specified in figure and table legends. The mean values and corresponding standard deviations (SD) are presented. When appropriate, data were treated with a two-tailed Student’s t-test in Microsoft Office Excel 2013 (Microsoft) and confidence intervals were calculated for the given parameters to test a statistically significant difference in means between two experimental datasets.

## 3 Results and discussion

### 3.1 Metabolic flux analyses elucidate xylose metabolism in *P. putida* PD310

The previously constructed strain *P. putida* PD310 with deleted glucose dehydrogenase gene (*gcd*) and recombinant xylose isomerase pathway (XylAB) and xylose/H^+^ symporter (XylE) from *E. coli* (**Fig. 1**) utilized D-xylose without accumulation of D-xylonate as a by-product (Dvořák and de Lorenzo, 2018). However, PD310 grew four-fold slower on xylose (0.11 h^-1^) than on glucose and exhibited a 5-times longer lag phase on the pentose sugar (∼10 h) (**Table 2, Supplementary Fig. S1**). This suboptimal growth of *P. putida* on xylose prompted investigations into the metabolic bottlenecks hindering efficient utilization of this non-native substrate.

**Table 2.** Growth parameters of *Pseudomonas putida* EM42 mutant strains in batch cultures with D-glucose or D-xylose as a sole carbon source.

| Strain and carbon source | $\mu_{\max}$ (h<sup>-1</sup>)<sup>1</sup> | $Y_{X/S}$ (g<sub>CDW</sub> g<sub>S</sub><sup>-1</sup>)<sup>2</sup> | $q_S$ (mmols g<sub>CDW</sub><sup>-1</sup> h<sup>-1</sup>)<sup>2</sup> | Lag phase (h)<sup>1</sup> |
| --- | --- | --- | --- | --- |
| CTRL GLU | 0.49 ± 0.01 | n.d. | n.d. | 2.08 ± 0.22 |
| PD310 GLU | 0.46 ± 0.01 | 0.38 ± 0.02 | 6.32 ± 0.26 | 1.40 ± 0.14 |
| EM42Δ*gcd*<br>*xylABE*<sup>+</sup> XYL | 0.09 ± 0.00 | n.d. | n.d. | 35.04 ± 0.60 |
| PD310 XYL | 0.11 ± 0.00 | 0.31 ± 0.03 | 1.97 ± 0.23 | 10.04 ± 0.53 |
| PD584 XYL | 0.13 ± 0.00 | 0.35 ± 0.01 | 2.22 ± 0.08 | 4.60 ± 0.44 |
| EM42Δ*gcd*Δ*gnd*<br>*xylABE*<sup>+</sup> XYL | 0.06 ± 0.00 | n.d. | n.d. | 32.27 ± 5.68 |
| PD689 XYL | 0.10 ± 0.00 | 0.31 ± 0.02 | 1.12 ± 0.06 | 18.25 ± 1.28 |
| PD584 tt L3 XYL | 0.23 ± 0.01 | 0.32 ± 0.01 | 3.98 ± 0.06 | 4.78 ± 0.35 |
| PD584 L3 XYL | 0.21 ± 0.01 | 0.41 ± 0.07 | 3.07 ± 0.46 | 3.40 ± 0.60 |
| PD689 tt L1 XYL | 0.21 ± 0.00 | 0.38 ± 0.02 | 3.54 ± 0.14 | 3.70 ± 0.35 |

Abbreviations: CTRL, *P. putida* EM42 Δ*gcd* pSEVA2213; GLU, D-glucose; XYL, D-xylose; n.d.: not determined
<sup>1</sup>The $\mu_{\max}$ and growth lag were calculated from growth data obtained from cultures in 48-well plates using the deODorizer program (Swain et al., 2016). Shown values are means from three (cultivations on glucose) or six (cultivations on xylose) biological replicates ± standard deviation.
<sup>2</sup>The biomass yield ( $Y_{X/S}$ ) and specific carbon uptake rate ( $q_S$ ) were calculated from data obtained from cultures in Erlenmeyer flasks. Shown values are means from three biological replicates ± standard deviation.

It has been demonstrated that adaptive laboratory evolution can boost the growth rate of a *P. putida* strain, which expresses the exogenous isomerase pathway and lacks Gcd, to approximately 0.3 h^-1^ on xylose (Elmore et al., 2020; Meijnen et al., 2008). Comparing growth data from different studies can be unreliable due to variations in experimental conditions. However, the findings of these authors suggest that *P. putida gcd*^-^ *xylABE*^+^ has the capacity to utilize xylose more efficiently than observed for PD310. In 2012, Meijnen et al. reported complex metabolic and regulatory re-arrangements underlying efficient xylose catabolism in an evolved recombinant strain of *P. putida* S12 (Meijnen et al., 2012). More recently, Elmore and co-workers (2020) emphasized the crucial role of the XylE transporter and its enhanced expression in promoting growth of evolved *P. putida* KT2440 (Elmore et al., 2020). In our work, we expected sufficient expression of the *xylE* gene as its expression from the low-copy-number plasmid pSEVA2213 is controlled by the strong constitutive P_EM7_ promoter and a strong synthetic ribosome binding site (RBS) with theoretical translation rate of 12,544 a.u. (Salis et al., 2009). While the expression of heterologous pathways and transporter may pose a hindrance to *P. putida*’s metabolism, the comparable growth rates of PD310 and the control strain without the synthetic *xylABE* operon (**Table 2**) indicated that this was not the case.

Therefore, we moved our attention to potential bottlenecks in the central carbon metabolism of *P. putida* and mapped the catabolism of xylose in PD310 using ^13^C-based metabolic flux analysis (MFA) (Wiechert, 2001). Distribution of carbon fluxes was previously determined for *P. putida* KT2440 or its derivatives cultured on glucose, benzoate, glycerol, gluconate, or succinate (Beckers et al., 2016; Ebert et al., 2011; Kohlstedt and Wittmann, 2019; Kukurugya et al., 2019; Nikel et al., 2015) but never for xylose-grown *P. putida* with the embedded isomerase pathway. The analysis performed with 1,2-^13^C D-xylose identified a partially cyclic upper xylose metabolism in engineered *P. putida* (**Fig. 2A, Supplementary File S1**). Theoretically, every three moles of xylulose 5-phosphate (X5P) produced by the isomerase pathway from xylose should yield two moles of fructose 6-phosphate (F6P) and one mole of glyceraldehyde 3-phosphate (G3P) (Volke et al., 2021). The MFA indeed showed that the majority (89 %) of the carbon that initially entered the non-oxidative branch of the pentose phosphate pathway (PPP) via X5P was converted into F6P and further to 6-phosphogluconate (6PG) through the reactions of glucose 6-phosphate isomerase (Pgi-I and Pgi-II), glucose 6-phosphate 1-dehydrogenase (ZwfA, ZwfB, ZwfC), and 6– phosphogluconolactonase (Pgl).

Flux through these reactions is much lower in wild-type *P. putida* KT2440 grown on glucose, which is preferentially utilized via the periplasmic oxidative route and directly funneled into the Entner-Doudoroff (ED) pathway (**Fig. 1**) (Kohlstedt and Wittmann, 2019; Kukurugya et al., 2019; Nikel et al., 2015). At the 6PG node, the flux from xylose branches (**Fig. 2A**). Over 50 % of the carbon enters the ED pathway while more than one-third is cycled back into the non-oxidative PPP via the 6-phosphogluconate dehydrogenase (Gnd) reaction to replenish the ribulose 5-phosphate (Ru5P) and ribose 5-phosphate (R5P) pools. R5P is needed together with X5P as substrate for transketolase Tkt. The Ru5P replenishment and partial carbon cycle via decarboxylation by Gnd seemed to compensate for the relatively weak flux through the ribulose-5-phosphate 3-epimerase (Rpe) reaction. Similar flux distributions were observed by Meijnen et al. (Meijnen et al., 2012), who also found that ALE resulted in increased channeling of 6PG into the non-oxidative PPP, which appeared to be crucial for improving xylose utilization in engineered *P. putida* S12. The bifurcation reduced ED pathway flux in xylose-grown PD310 compared to wild-type *P. putida* cultured on glucose (**Fig. 2A**) (Kohlstedt and Wittmann, 2019; Kukurugya et al., 2019; Nikel et al., 2015). ED pathway flux merges with the flux from PPP in the glyceraldehyde 3-phosphate node and continues through the lower glycolysis to pyruvate.

Another feature of xylose metabolism in PD310 revealed by MFA was a negligible flux through the EMP pathway reactions catalyzed by fructose 1,6-bisphosphatase (Fbp), fructose 1,6-bisphosphate aldolase (Fba), and triose phosphate isomerase (TpiA) (**Fig. 2A**). The flux was ∼3–8-fold lower than in KT2440 grown on glucose (Kohlstedt and Wittmann, 2019; Kukurugya et al., 2019; Nikel et al., 2015). During growth on glucose, the gluconeogenic operation of the upper EMP route in the EDEMP cycle allows for partial recycling of triose phosphates back into hexose phosphates and secures the resistance of *P. putida* to oxidative stress through supply of NADPH (generated by Zwf) (Nikel et al., 2021). Carbon cycling through the upper EMP pathway associated with increased NADPH availability was also reported for *P. putida* grown on glycerol (Beckers et al., 2016). In xylose-grown PD310, such operation of Fbp, Fba, and TpiA would be redundant because NADPH generation is maintained by the strong activities of Zwf and Gnd (**Fig. 2A**).

The activated glyoxylate shunt (isocitrate lyase AceA and malate synthase GlcB) in PD310 suggests a need to balance reducing cofactors (NADPH or NADH that inhibit the activity of isocitrate dehydrogenase) or decrease loss of CO_2_ (**Fig. 2A**) (Beckers et al., 2016; Ebert et al., 2011; Meijnen et al., 2012; Sudarsan et al., 2016). An active glyoxylate shunt was reported in *P. putida* KT2440 grown on glycerol (Beckers et al., 2016), benzoate (Sudarsan et al., 2016), or a mixture of glucose and succinate (La Rosa et al., 2015) but not on glucose alone (Kohlstedt and Wittmann, 2019; Kukurugya et al., 2019; Nikel et al., 2015). Meijnen et al. (2012) recognized an operating glyoxylate shunt in their S12 strain evolved on xylose and attributed this phenomenon to the level of reducing cofactors, which was increased compared with cells cultured on glucose (Meijnen et al., 2012). Given the high fluxes measured for the four central dehydrogenases – Zwf, Gnd, pyruvate dehydrogenase complex, and malate dehydrogenase (the middle two enzymes also have decarboxylating activity) – the same scenario is also possible for strain PD310 (**Fig. 2A**).

MFA confirmed that malic enzyme activity greatly contributed to the pyruvate pool (**Fig. 2A**) similar to previous analyses on glucose (Kohlstedt and Wittmann, 2019; Kukurugya et al., 2019; Nikel et al., 2015). However, xylose metabolism differs in its low pyruvate carboxylase activity and enhanced fluxes through the pyruvate dehydrogenase and malate dehydrogenase reactions.

As the next step, we applied flux balance analysis (Orth et al., 2010) using the most recent genome-scale model of *P. putida* KT2440 iJN1463 (Nogales et al., 2020) with modifications (see Materials and methods) to compute metabolic fluxes supporting optimal growth on xylose. We compared this optimal flux distribution with MFA data (**Fig. 2, Supplementary File S2**) and found several differences, including an almost evenly distributed flux from X5P into reactions of Rpe, Tkt, and Tal, and zero flux through Gnd, the glyoxylate shunt, and the malic enzyme reaction in FBA. Enhanced fluxes from X5P to Ru5P and higher TCA cycle activity in the FBA simulation probably fully replaced the carbon cycling via Gnd observed in vivo. FBA also computed a reduced (by ∼33 %) formation of hexoses F6P and G6P and, consequently, a lower flux through glucose 6-phosphate dehydrogenase Zwf. Importantly, at the same specific xylose uptake rate (*q*_S_ = 1.45 mmol g_CDW_^−1^ h^−1^) determined during the ^13^C labeling experiment, the model unconstrained with MFA data (FBA model) grew ∼41 % faster (µ = 0.118 h^-1^) than PD310 cells (µ = 0.083 h^-1^ ± 0.02, note that differences between the growth rate and xylose uptake rate determined during the labeling experiment and the values reported in **Table 2** are due to different experimental conditions). When the two metabolic models – one constrained with upper and lower bounds from MFA (MFA model) and FBA model were compared throughout the range of stepwise increased MFA bounds or xylose uptake rates, respectively, the FBA model always provided ˃40% higher growth rate (Materials and methods, **Supplementary Fig. S2**). In addition, the comparison of the sum of fluxes through oxidoreductase reactions that form reducing equivalents NADPH and NADH revealed a 1.6-fold (NADPH) and 1.5-fold (NADH) higher values in the MFA model (2.79 vs. 1.75 mmol g_CDW_^−1^ h^−1^ for NADPH producing reactions and 6.48 vs. 4.34 mmol g_CDW_^−1^ h^−1^ for NADH producing reactions). Also, the net rate of CO_2_ production determined in MFA (3.97 ± 0.12 mmol g_CDW_^−1^ h^−1^) was 1.6-fold higher than the net rate calculated in FBA (2.43 mmol g_CDW_^−1^ h^−1^, **Supplementary File S3**). As discussed above, the surplus of NAD(P)H and loss of carbon in the form of CO_2_ observed in PD310 strain grown on xylose may be caused, at least partially, by high fluxes through Zwf and Gnd and be responsible for glyoxylate shunt activation. Also, the lower biomass yield of PD310 on xylose compared with glucose can be explained by the high activity of decarboxylating enzymes such as Gnd (**Table 2**).

### 3.2 Targeted gene knockouts expose important nodes of xylose metabolism in *P. putida*

Next, we verified the importance of some key enzymes of the EDEMP cycle for the growth of PD310 on xylose. Genes encoding Pgi-I (PP_1808), Pgi-II (PP_4701), Gnd (PP_4043), and 6-phosphogluconate dehydratase (Edd, PP_1010) were knocked-out in PD310 and the growth of the three resulting mutants was tested on solid and in liquid medium (**Fig. 3**). Experiments on M9 agar plates showed that none of the deletions was detrimental for growth on citrate (gluconeogenic growth regime) (**Fig. 3B**) and only the *edd* deletion disabled growth on glucose. In contrast, growth on xylose was affected by all three deletions. The different effects of individual deletions were more pronounced in liquid medium (**Fig. 3C**). The Δ*gcd* Δ*pgi*-I Δ*pgi*-II and Δ*gcd* Δ*edd* mutants did not show any growth within three days, confirming the essentiality of Pgi and Edd for xylose catabolism in PD310. The importance of Pgi for growth on xylose was recently demonstrated in *P. putida* KT2440 harboring a recombinant xylose isomerase pathway (Ling et al., 2022). Interestingly, the Δ*gcd* Δ*gnd* mutant retained its growth capacity on xylose, though, with a reduced growth rate (0.06 ± 0.00 h^-1^) and substantially prolonged lag phase (32.3 ± 5.7 h) when compared to PD310 (**Fig. 3C**). This result showed that the carbon flux into the non-oxidative branch of PPP through Gnd is not an essential prerequisite for xylose utilization by *P. putida*, which was in line with the data from FBA.

**Figure 3.**
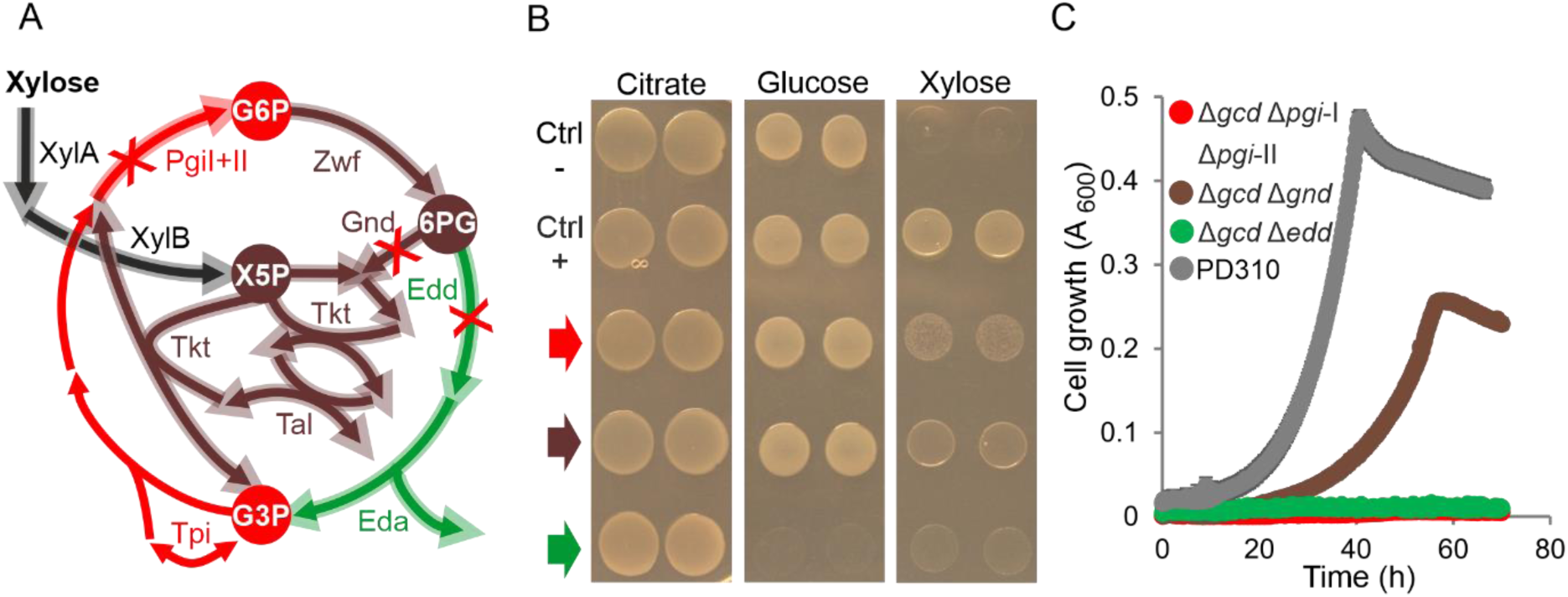
Growth of three *P. putida* deletion mutants on selected carbon sources including xylose. **(A)** Scheme of the EDEMP cycle with the incorporated xylose isomerase pathway (abbreviations and color coding are the same as in the previous figures) and highlighted reactions that were eliminated by knocking out the respective genes (red crosses). **(B)** Growth of Δ*gcd* Δ*pgi*-I Δ*pgi*-II (red arrow), Δ*gcd* Δ*gnd* (brown arrow), and Δ*gcd* Δ*edd* (green arrow) mutants on solid M9 agar medium with 2 g L^-1^ citrate, D-glucose, or D-xylose used as the sole carbon and energy source. Ctrl-stands for negative control (*P. putida* EM42 Δ*gcd* with empty pSEVA2213) and Ctrl+ stands for positive control (PD310). Cells pre-cultured in LB medium were washed with M9 medium and 10 µL of cell suspension of OD 2.0 was dropped on the agar and incubated for 24 h (agar with glucose or citrate) or 48 h (agar with xylose) at 30°C. **(C)** Growth of the three mutants and control PD310 in M9 minimal medium with 2 g L^-1^ D-xylose in 48-well microtiter plate. Data points are shown as mean from at least three biological replicates.

The flux analyses and experiments with deletion mutants pictured the distribution of carbon in the EDEMP cycle of xylose-grown PD310. Notably, the experimentally determined distribution of fluxes – especially in the PPP part of the EDEMP cycle – resembles the situation in glucose-grown *P. putida* exposed to oxidative stress (Nikel et al., 2021). Since the complex biochemical network of *P. putida* evolved primarily towards utilization of organic acids, aromatic compounds, or glucose (Rojo, 2010), it is plausible that the introduction of an exogenous xylose isomerase pathway resulted in a certain level of metabolic imbalance and, consequently, in the slow growth rate, which was far from values achievable on native substrates. Therefore, we aimed at knowledge-driven de-bottlenecking of xylose metabolism in the following steps.

### 3.3 Measurement of activities of EDEMP cycle enzymes reveals first metabolic bottleneck

To get an even deeper insight into the operation of the EDEMP cycle of PD310 grown on xylose, we measured activities of eight enzymes that contribute to the cycle together with activities of XylA and XylB (**Figs. 3A and 4**, **Supplementary methods**). The activity of the ED pathway (Edd and Eda) was measured in a combined assay (Stephenson et al., n.d.). The specific activities (U mg^-1^_total protein_) were determined in cell-free extracts (CFE) prepared from PD310 cultures in mid-exponential phase in LB medium, M9 medium with 2 g L^-1^ glucose, or M9 medium with 2 g L^-1^ xylose. The specific activity of both XylA and XylB was significantly higher (P<0.05) on xylose than on glucose substrate. This probably reflects a selective pressure due to the employment of these enzymes during the growth on the pentose. Activities of XylB, Gnd, Zwf, Pgi, and Edd-Eda in cells cultivated on either sugar were higher (P<0.05) than the activities in cells grown in LB medium (**Fig. 4A**). In the case of Zwf and Edd-Eda the difference vcan be attributed to the de-repression of the respective genes in cells consuming glucose or xylose. The *zwf*-*pgl*-*eda* and *edd*-*glk* operons together with the *gap*-1 gene are controlled by the HexR transcriptional repressor (Udaondo et al., 2018) (**Fig. 4B**). Binding 2-keto-3-deoxy-6-phosphogluconate (KDPG), an intermediate of the ED pathway (**Fig. 2**), which is formed during catabolism of glucose and xylose, causes HexR dissociation and transcriptional activation (Daddaoua et al., 2010). Interestingly, the de-repression seemed to be more efficient or faster on glucose because activities of Zwf and Edd-Eda measured in cells cultured on this sugar were ˃2-fold higher than on xylose (**Fig. 4A**). On the other hand, activities of transketolase, transaldolase, and Tpi were comparable over all three tested conditions (P˃0.05), which indicates that the levels of these enzymes in *P. putida* cells are rather constant irrespective of the substrate.

**Figure 4.**
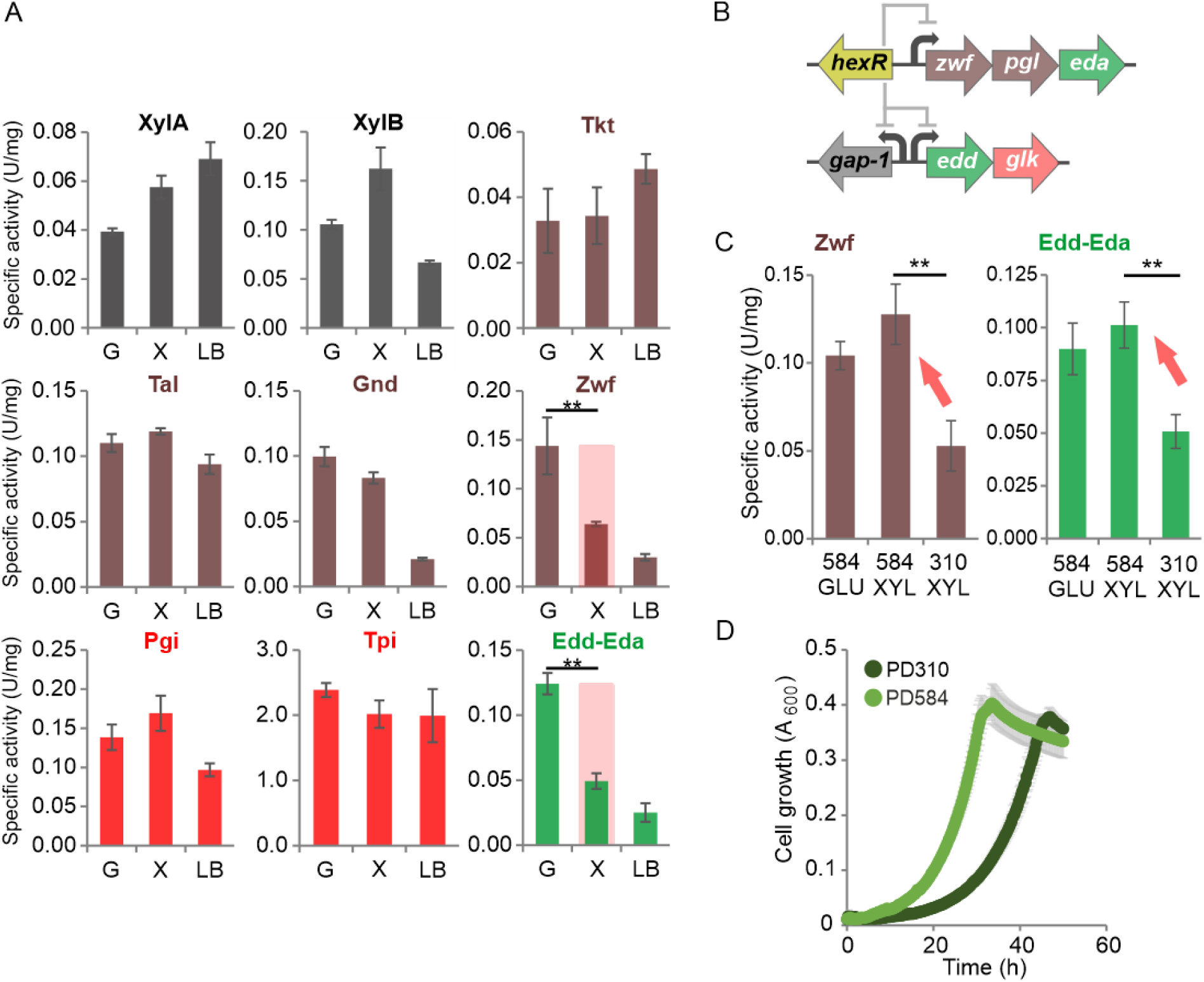
Activities of XylA, XylB and selected EDEMP cycle enzymes in *P. putida* PD310 cells and the effect of *hexR* gene deletion. **(A)** Specific activities of 10 selected enzymes (activity of Eda and Edd was measured in a combined assay) were determined in cell-free extracts (CFE) prepared from PD310 cells cultured till mid-exponential phase in rich LB medium (LB), in M9 medium with 2 g L^-1^ glucose (G), or in M9 medium with 2 g L^-1^ xylose (X) as detailed in Materials and methods section. **(B)** Genetic organization of relevant genes in operons regulated by HexR transcriptional regulator. The elements in this scheme are not drawn to scale. **(C)** Specific activity of Zwf or Edd-Eda measured in CFE from PD310 or PD584 cells grown in M9 medium with 2 g L^-1^ glucose (GLU), or in M9 medium with 2 g L^-1^ xylose (XYL). **(D)** Growth of PD310 and a new strain PD584 with additional deletion of *hexR* gene in M9 minimal medium with 2 g L^-1^ D-xylose in 48-well microtiter plate. Data are shown as mean ± SD from three biological replicates (A) or six biological replicates (C,D). Asterisks denote statistically significant difference between two means at P < 0.01 (**).

Low activities of Zwf and the ED pathway enzymes in cells grown on xylose could pose a bottleneck for pentose catabolism in this part of the EDEMP cycle. Del Castillo and colleagues (del Castillo et al., 2008) and Bentley et al. (Bentley et al., 2020) previously showed that the deletion of *hexR* gene can de-repress *zwf*-*pgl*-*eda* and *edd*-*glk* operons. In the latter study, the de-repression improved growth of *P. putida* KT2440 mutant on glucose and increased production rate of *cis*,*cis*-muconic acid. Down-regulation of *hexR* was identified in evolved *P. putida* S12 and linked to the boosted xylose-utilization phenotype (Meijnen et al., 2012). Therefore, we argued that the removal of the repressor could have a positive effect on xylose metabolism in PD310. The *hexR* gene (PP_1021) was deleted in *P. putida* EM42 and in *P. putida* EM42 Δ*gcd* strains. The purpose of the deletion on the background of wild-type *P. putida* EM42 strain was to verify the connection between HexR and periplasmic oxidation of xylose to xylonate, which was observed in *P. putida* S12 (**Supplementary results**, **Supplementary Fig. S3**) (Meijnen et al., 2012). The double deletion mutant *P. putida* EM42 Δ*gcd* Δ*hexR* pSEVA2213_*xylABE*, named PD584, showed improved growth compared with PD310 (**Fig. 4D**, **Table 2, Supplementary Fig. S3**). Notably, the lag phase on xylose was reduced by more than 2-fold, from 10.0 to 4.6 h. The reduced lag phase was also measured for PD584 grown on hexose substrates glucose and fructose (**Supplementary Fig. S4**). The significantly increased activities of Zwf and the ED pathway in PD584 on xylose (**Fig. 4C**) underpin that this improvement can be mainly attributed to the de-repression of EDEMP cycle enzymes. The high growth rate of PD584 on fructose (µmax = 0.30±0.00 h^-1^, **Supplementary Fig. S4**) which is, similarly to xylose, metabolized through Pgi, Zwf, Pgl, and the ED pathway (Volke et al., 2021) indicated that these reactions may not be limiting in xylose catabolism either. However, the growth rate of PD584 on xylose increased only modestly (by ∼18%) when compared to PD310 (**Table 2**) and we thus sought additional targets to eliminate further bottlenecks in the metabolism. To this end, we combined additional rational interventions in the EDEMP cycle guided by the MFA/FBA comparison with adaptive laboratory evolution.

### 3.4 Knowledge-driven engineering of PPP combined with the adaptive laboratory evolution boosts growth on xylose

The most evident differences between the distribution of fluxes in the EDEMP cycle in FBA and MFA were in the PPP reactions. During growth on native substrates such as glucose or fructose, the role of PPP in *P. putida* is rather complementary and mostly anabolic (Nikel et al., 2021). Periplasmic glucose oxidation is the preferred route for glucose uptake and therefore only a smaller fraction of carbon (∼20–50 %) flows through Zwf and Pgl and even much less (∼1–10 %) is directed to the non-oxidative branch of PPP to provide metabolic precursors for nucleotides and some amino acids (Kohlstedt and Wittmann, 2019; Kukurugya et al., 2019). The situation changes drastically upon the exposure of *P. putida* to oxidative stress when Zwf and Gnd start to generate reducing equivalents for the elimination of reactive oxygen species (ROS) (Nikel et al., 2021), or during the growth of recombinant *P. putida* on xylose where PPP is the metabolic entrypoint for this sugar. The reported ability of *P. putida* to adapt its metabolic fluxes in response to ROS and increase the activity of Zwf and Gnd multiple times without diminishing its growth rate on glucose (Nikel et al., 2021) indicates that the bacterium should have some capacity to adjust its PPP in favor of non-native pentose metabolism. However, during the growth on xylose, all carbon enters the PPP at the point of X5P, not G6P, as is the case during the *P. putida*’s physiological response to ROS. Hence, suboptimal activity or set-up of the non-oxidative branch of PPP might still be limiting xylose metabolism.

Elmore and co-workers (2020) endowed a recombinant KT2440 *xylABE*^+^ strain with additional *E. coli* transketolase (*tktA*) and transaldolase (*talB*) genes (Elmore et al., 2020). Such an intervention based on exogenous genes from a bacterium that grows well on pentoses significantly accelerated the growth of the engineered KT2440 on xylose (Elmore et al., 2020). A similar approach was previously found to be instrumental for establishing or enhancing non-native pentose metabolism in other bacteria as well, e.g., in *Zymomonas mobilis* (Zhang et al., 1995). Being inspired by these studies and our flux analyses, we decided to modulate the PPP in strain PD584 to further boost xylose utilization. To mimic the FBA scenario with zero flux through Gnd reaction, we first prepared a *P. putida* strain designated PD689 with deletions of *gcd*, *hexR* and *gnd* and harboring the pSEVA2213_*xylABE* plasmid (**Tables 1 and S1**). The absence of the *gnd* gene was verified by the lack of enzymatic activity of 6-phosphogluconate dehydrogenase in CFEs of PD689 (**Supplementary Fig. S5**). PD689 showed a 1.7-fold higher growth rate on xylose (0.10 ± 0.00 h^-1^ vs. 0.06 ± 0.00 h^-1^) and an almost 2-fold shorter lag phase (18.3 ± 1.3 h vs. 32.3 ± 5.7) than the parental strain EM42 Δ*gcd* Δ*gnd* pSEVA2213_*xylABE* confirming the beneficial effect of *hexR* deletion. In contrast, PD689 demonstrated a lower growth rate and 4-fold longer lag compared to PD584 reference (**Table 2**, **Supplementary Fig. S5**). This comparison again suggested that the null mutation of *gnd* may decelerate growth but is not fatal for xylose utilization by *P. putida* (**Fig. 3**).

We then integrated an expression cassette bearing either *talB*-*tktA* or *talB*-*tktA*-*rpe*-*rpiA* synthetic operon assembled from *E. coli* genes into the genome of PD584 and PD698. We presumed that the activity of Rpe and RpiA, whose genes were included in the second variant of the operon, could support the conversion of X5P to R5P and thus replenish metabolites depending on the Gnd reaction. The expression cassettes were inserted randomly into the host’s chromosome using mini-Tn5 delivery plasmid pBAMD1– 4 (Martínez-García et al., 2014a). Transformation of bacteria with this plasmid generally results in a single insertion per cell and allows to take advantage of chromosome position effect on the expression of the implanted genes (Bujdoš et al., 2023). Transformants with an optimal expression of the inserted genes can then be selected from the library based on improved phenotype – in our case, accelerated growth on xylose. To ensure transcription at various positions on chromosome, the expression cassettes were complemented with the constitutive P_EM7_ promoter. We presumed that the multiple rational interventions in our *P. putida* strains could cause temporary impairment of xylose utilization due to metabolic rewiring. Therefore, we combined the random integration of the synthetic operons into the chromosome of PD584 and PD689 with ALE (**Fig. 5, Materials and methods**), which has been often found useful for finetuning rationally engineered microbial cell factories and was previously adopted for complementing rational design of *P. putida* strains with an expanded substrate spectrum (Bator et al., 2020; Elmore et al., 2020; Espeso et al., 2020; Lim et al., 2021; Meijnen et al., 2008; Sandberg et al., 2019).

**Figure 5.**
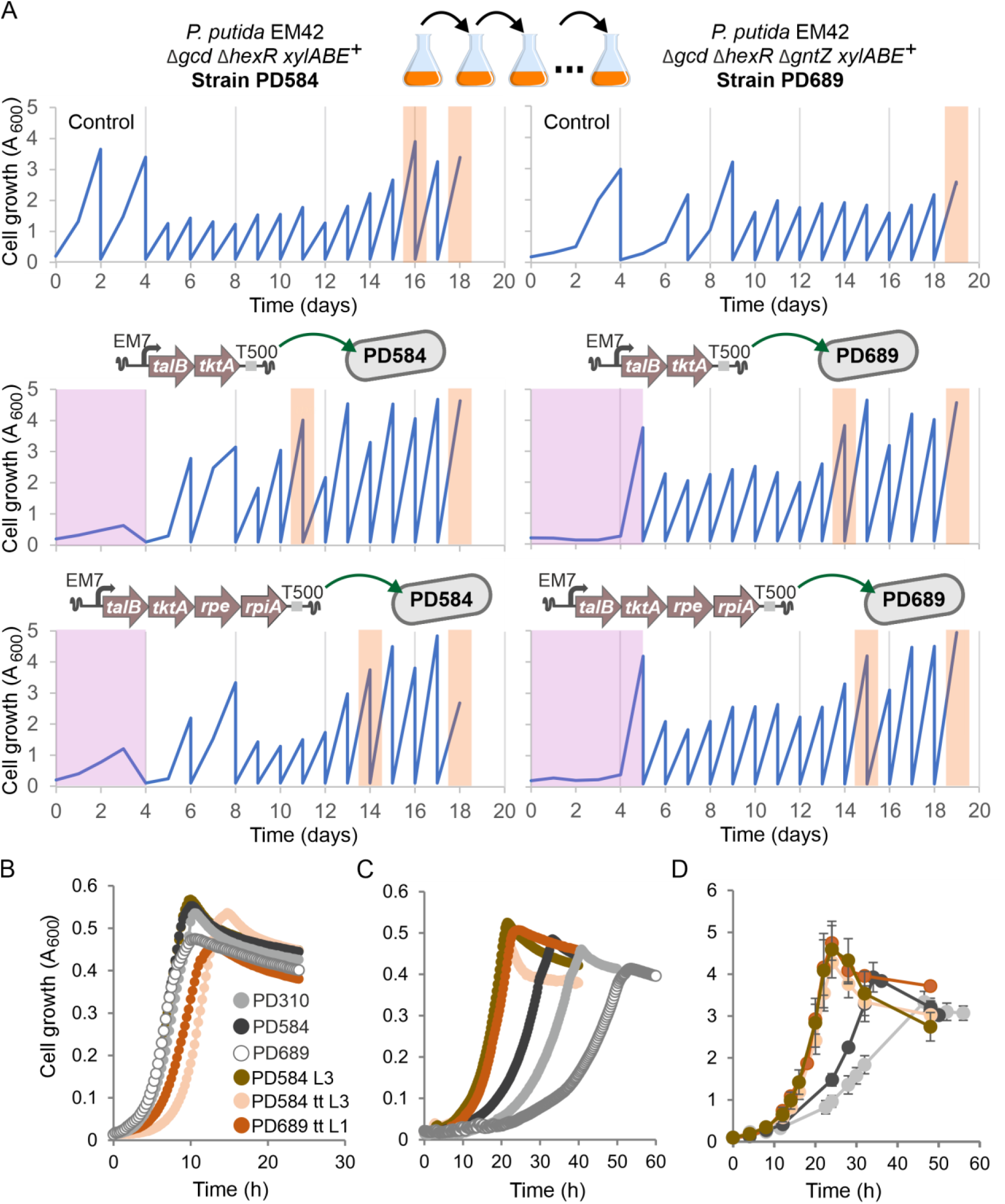
Adaptive laboratory evolution (ALE) on xylose of *Pseudomonas putida* PD584 and PD689 strains with and without integrated synthetic operons bearing pentose phosphate pathway (PPP) genes from *Escherichia coli* (A) and growth assays with the best candidates selected after ALE (B-D). **(A)** Cells were cultured in 20 mL of M9 medium with 5 g L^-1^ D-xylose and kanamycin and passaged in time intervals indicated in the graphs (upper two graphs depict ALE of PD584 and PD689 controls without implanted PPP genes and lower four graphs depict ALE of PD584 and PD689 transformants after insertion of pBAMD1-4 plasmid constructs with PPP genes). The pre-selection period where PD584 and PD589 transformants were cultured in M9 medium with xylose and two antibiotics (Km, Sm) is indicated by pink shading. Days in which samples of the cultures were withdrawn to prepare glycerol stocks and select individual clones are highlighted by orange shading. The first withdrawal occurred when the turbidity of a given culture (OD_600_) first reached the value of ≥3.5 within 24 h of growth (the only exception was the control culture of PD689 which did not pass this threshold during the evolutionary experiment). The elements in the operon schemes are not drawn to scale. Abbreviations: P_EM7_, constitutive promoter; T500, transcriptional terminator. The *talB*, *tktA*, *rpe*, and *rpiA* denote genes that encode *E. coli* transaldolase B, transketolase A, ribulose-phosphate 3-epimerase, and ribose-5-phosphate isomerase, respectively. **(B)**+**(C)** Growth of PD310, PD584, PD689, PD584 L3, PD584 tt L3, and PD689 tt in M9 medium with 2 g L^-1^ D-glucose (B) or 2 g L^-1^ D-xylose (C) in 48-well microtiter plate. Data are shown as mean from six (PD584 L3, PD584 tt L3, PD689 tt L1) or three (controls PD310, PD584, PD689) biological replicates. Error bars are omitted for clarity. **(D)** Growth of PD310, PD584, PD584 L3, PD584 tt L3, and PD689 tt L1 (color coding is the same as in B and C) in M9 medium with 5 g L^-1^ D-xylose in shake flasks. Data are shown as mean ± SD from three biological replicates.

We selected three candidates that demonstrated the fastest growth in the shake-flask format and reached OD600 ˃3.5 within 24 h. All three clones, designated PD584 L3, PD584 tt L3, and PD689 tt L1, were isolates from the end of the evolutionary experiment. Interestingly, strain PD584 L3 was isolated from the control culture and strains PD584 tt L3 and PD689 tt L1 from cultures of transformants with the inserted pBAMD1-4_*talB*-*tktA* construct. All isolates with the integrated *talB*-*tktA*-*rpe*-*rpiA* operon that were pre-selected based on the growth tests in the 48-well plate grew significantly slower in shake flasks than the three best candidates and some of them barely outperformed the PD584 or PD689 template (**Supplementary Fig. S6**). The fact that the high turbidities (OD600 ˃3.5) that were reached in 48-well plates with the *talB*-*tktA*-*rpe*-*rpiA*^+^ transformants (**Fig. 5**) could not be reproduced in shake-flasks (i.e., these clones were not able to maintain the desired phenotype across different cultivation formats) may indicate that a mere short-term adaptation via alterations in gene expression and physiology rather than evolution of genotypes might have occurred in these cell populations (Brooks et al., 2011).

The outcomes of ALE and subsequent screening experiments showed that: (i) the expression of *E. coli rpe* and *rpiA* genes for ribulose-phosphate 3-epimerase and ribose-5-phosphate isomerase is not needed for the improved xylose utilization in *P. putida* and it may rather pose additional burden on cells (**Figs. 5 and S6**), and (ii) ALE of PD584 strain can provide variants with substantially accelerated growth on xylose even without supplementation of any additional exogenous genes. The latter finding is intriguing especially in the context of other recent studies in which *P. putida* was endowed with *E. coli* genes and evolved for better growth on xylose (Elmore et al., 2020; Ling et al., 2022). It demonstrates that a similar outcome can be achieved with fewer engineering interventions. It is also worth noting that a control 3-week ALE of *P. putida* PD310 in two shake-flask cultures did not result in enhanced growth of this strain on xylose (OD_600_ after 48-h passage intervals never exceeded the value of 3.6, **Supplementary Fig. S7**). The *hexR* deletion in PD584 and PD689 may thus represent a key de-bottlenecking step that accelerated further improvement of xylose metabolism in these strains during the ALE experiment. Growth assays in 48-well plates and Erlenmeyer flasks confirmed that all three evolved strains utilize xylose more efficiently than their ancestors (**Table 2**, **Figs. 5B and 5D, Supplementary Fig. S8**). The growth rate of PD584 L3, PD584 tt L3, and PD689 tt L1 increased by 62, 77, and 110 %, respectively, compared with the ancestral strains PD584 and PD689. PD584 L3 and PD689 tt L1 also exhibited higher biomass yield and a shorter growth lag (**Table 2**). The 5-fold reduction of the lag phase of PD698 tt L1 (3.70 h) compared to PD689 (18.25 h) was remarkable. PD689 tt L1 utilized xylose as efficiently as PD584 L3 and PD584 tt L3 strains that, however, originated from the much better-growing ancestor PD584 (**Table 2**).

We also tested the growth of the three evolved strains and their ancestors on glucose (**Fig. 5C**). Glucose is the most abundant hexose sugar in (hemi)cellulose and its simultaneous utilization with xylose is a desirable property of any microbial cell factory applicable in lignocellulose biorefineries (Dvořák and de Lorenzo, 2018; Elmore et al., 2020; Valdivia et al., 2016). The control strains (PD310, PD584, PD689) and PD584 L3 all grew well on glucose (µ ≥ 0.50 h^-1^, growth lag ∼ 1.0 h) with PD584 and PD584 L3 being the fastest (0.58 ± 0.01 h^-1^ and 0.57 ± 0.00 h^-1^, respectively). On the other hand, two evolved strains PD689 tt L1 and PD584 tt L3 showed reduced growth rate (0.42 ± 0.01 h^-1^ and 0.45 ± 0.01 h^-1^, respectively) and a prolonged lag phase (2.21 ± 0.20 h and 4.47 ± 0.15 h, respectively). Therefore, it seems that in case of the slower-growing strains, tailoring of the PPP and subsequent ALE on xylose affected the metabolism of glucose. Further growth experiments with PD584 L3 and PD689 tt L1 confirmed that both new strains also maintained the previously reported ability of the ancestral PD310 to co-utilize glucose and xylose (**Supplementary Fig. S9**) (Dvořák and de Lorenzo, 2018).

### 3.5 Whole-genome sequencing of evolved *P. putida* strains unveils the causes of improved growth on xylose

To reveal the causes of improved growth on xylose, the strains PD584 L3, PD584 tt L3, and PD689 tt L1 were subjected to further characterization. Firstly, the strains were sequenced together with the reference strain PD584 and one of the slower growing mutant PD584 ttrr L2 using Oxford Nanopore technology (**Supplementary methods, Supplementary Table S3**). We aimed to verify the chromosomal integration of the expression cassettes *talB*-*tktA* and *talB*-*tktA*-*rpe*-*rpiA*. The sequencing identified the locus of the cassette integration in PD689 tt L1 (PP_1181, which encodes a two-component system response regulator from OmpR family) and in PD584 ttrr L2 (PP_1145, which encodes RNA polymerase-associated protein RapA) and confirmed the absence of the cassette in controls PD584 and PD584 L3. The unstable phenotype of PD584 ttrr L2 strain can be related to the integration of the *talB*-*tktA*-*rpe*-*rpiA* cassette in the *rapA* locus. RapA is a transcription regulator that stimulates RNA polymerase recycling and the disruption of the corresponding gene may thus cause divergence in global gene expression pattern (Jin et al., 2011). Surprisingly, sequencing revealed that the *talB*-*tktA* cassette was absent in the genome of the PD584 tt L3 strain. The cassette could have been lost during the ALE, or its integration into the chromosome by the Tn5 minitransposon system failed already at the beginning of the evolution experiment. In either case, similarly to PD584 L3, the PD584 tt L3 showed improved growth on xylose, which confirmed that xylose metabolism on the PD584 background can be accelerated without exogenous PPP enzymes. Due to the fact that its growth on glucose was impaired, the strain PD584 tt L3 was eliminated from further experiments.

The genomes of the reference strain PD584 and the two best xylose utilizing strains from ALE, PD584 L3 and PD689 tt L1, were further resolved by Illumina sequencing and complete whole-genome sequences were determined using hybrid assembly (**Supplementary methods, Supplementary Table S3**). We first verified the sequence of the *talB*-*tktA* expression cassette integrated into the chromosome of strain PD689 tt L1. Four silent mutations (three in *talB* and one in *tktA*) and one missense mutation (GCA→ACA resulting in Ala247→Thr247 substitution in TalB) were identified. The Ala247→Thr247 substitution is relatively distant from the binding pocket and can be found in functional TalB variants, for instance in the PDB crystal structure 4S2C (Stellmacher et al., 2016). Hence, we do not expect it to have a negative effect on the enzyme’s activity.

Due to multiple changes (deletions, insertions, single nucleotide exchanges), the chromosomes of PD584 L3 and PD689 tt L1 showed 99.6% and 99.4% pairwise identity, respectively, to PD584 reference (using whole-genome alignment by Mauve Plugin in Geneious Prime). Mutations were detected in non-coding intergenic regions, rRNA and tRNA genes, and protein-coding sequences mostly related to mobile elements (**Supplementary File S4**; note that engineered modifications were excluded from the list). Given the known variability of pseudomonad genomes, such modifications are not surprising (Silby et al., 2011). Amino acid sequences of 14 gene products in PD584 L3 and 10 gene products in PD689 tt L1 were affected (**Supplementary File S4**). Both PD584 L3 and PD689 tt L1 accumulated numerous mutations in the gene encoding the large adhesive protein LapA (PP_0168), the largest protein in *P. putida*. LapA is responsible for the initiation of biofilm formation and its maturation (El-Kirat-Chatel et al., 2014). Bentley and co-workers (2020) reported transposon insertions in *lapA* in their recombinant *P. putida* KT2440 strain evolved for better growth on glucose (Bentley et al., 2020). We argue that the prolonged shake-flask cultivation and multiple transfers of planktonic cells during our ALE experiment posed a selective pressure against biofilm formation and *lapA* became a target of this selection. The occurrence of multiple mutations in this gene may also be related to its specific sequence, which contains numerous repetitions (El-Kirat-Chatel et al., 2014).

An intriguing alteration uncovered in the genomes of PD584 and PD584 L3 was a multiplication of a large region (118,079 bp), discovered by an approximately 6-fold higher sequencing coverage from locus PP_2114 to PP_2219 (**Fig. 6A, Supplementary File S5**). The multiplication was absent in PD689 tt L1. As the two border open reading frames (ORFs) encode transposases, we presume that the multiplication occurred via repeated transposition events. The multiplied region includes 106 CDS (based on Prokka annotation) out of which 22 are genes of hypothetical or uncharacterized proteins. The remaining CDS encode transporters (2 genes), transcriptional regulators including regulators from the AraC and LysR family (11 genes), or enzymes and their subunits (43 genes), out of which 9 have an oxidoreductase function (**Supplementary File S5**). Importantly, the latter set includes genes of transaldolase Tal (PP_2168), glyceraldehyde 3-phosphate dehydrogenase GapB (PP_2149), and soluble pyridine nucleotide transhydrogenase SthA (PP_2151). The increased gene dosage of these two enzymes of the central carbon metabolism (**Fig. 2**) and the transhydrogenase, which transfers electrons between the redox pair NADPH/NAD^+^ helping *P. putida* maintain its redox balance (Ebert et al., 2011; Nikel et al., 2015), could have a positive effect on xylose utilization by PD584 and PD584 L3. However, the fact that the multiplication was also revealed in the ancestral strain PD584 indicated that this genome re-arrangement had already occurred during the manipulation with this strain prior to PPP gene introduction and ALE or even earlier in the strain PD310 (Dvořák and de Lorenzo, 2018). The transposition might have occurred during the re-streaking of the strain on agar plates with xylose, which we did to check the desired phenotype before glycerol stock preparation. To confirm the presence of the multiplication in PD310, we sequenced this strain together with the reference strain EM42 Δ*gcd*, which was used as a template for PD310 preparation. Additionally, we freshly transformed EM42 Δ*gcd* with the pSEVA2213_*xylABE* construct and compared the growth of fresh transformants with the original PD310 strain. Data from Nanopore sequencing confirmed that the multiplication (probably four copies of the PP_2114—PP_2219 region) was, indeed, present in the original PD310 strain published in 2018, but was not present in the EM42 Δ*gcd* template (**Supplementary Table S4**). The fresh EM42 Δ*gcd* pSEVA2213_*xylABE* transformants grew significantly slower on xylose (**Supplementary Fig. S10**). Their specific growth rate in liquid medium was similar to the growth rate of the original PD310, but the fresh clones passed through a very long lag phase (almost 40 h of slow linear growth) before they started growing exponentially (**Table 2**). This experiment demonstrated the positive effect of the multiplication, which was maintained in the whole lineage of PD310, PD584, and PD584 L3 strains.

**Figure 6.**
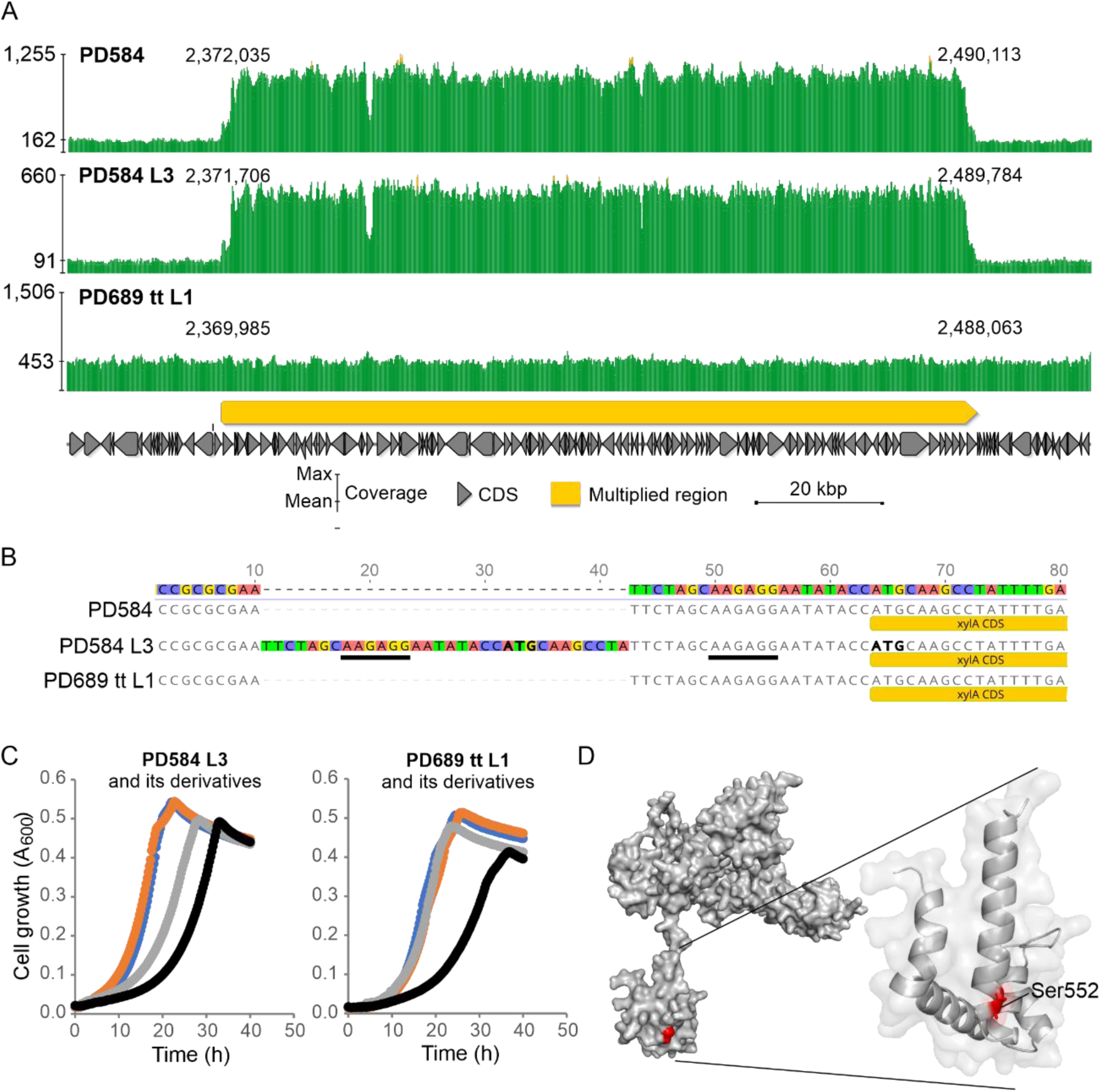
Characterization of *P. putida* EM42 strains PD584 L3 and PD689 tt L1, obtained from adaptive laboratory evolution on xylose, by whole-genome sequencing. **(A)** Multiplied (6x) region identified from the next-generation sequencing data in the chromosome of PD584 and PD584 L3. **(B)** 32 bp duplication identified upstream the *xylA* gene in pSEVA2213_*xylABE* plasmid isolated from PD584 L3. The duplication includes synthetic RBS AAGAGG (underlined), ATG codon (in bold) and eight following nucleotides from 5’-end of *xylA* gene. The coverage graphs in (A) as well as the sequence alignment in (B) were generated by Geneious Prime 2022.2.2. **(C)** Microtiter plate growth assays on xylose (2 g L^-1^) with PD584 L3 (left graph) and PD689 tt L1 (right graph) bearing re-implanted pSEVA2213_*xylABE* plasmid. PD584 L3 was used either intact (orange symbols) or its plasmid pSEVA2213_*xylABE* containing the duplication was removed and the strain was transformed with the same plasmid (blue symbols) or the original (flawless) pSEVA2213_*xylABE* plasmid (grey symbols). PD689 tt L1 strain was used either intact (orange symbols) or its flawless pSEVA2213_*xylABE* plasmid was removed and the strain was transformed with the mutated plasmid from PD584 L3 (blue symbols) or with the original flawless pSEVA2213_*xylABE* plasmid (grey symbols). Strain PD584 was used as a control (black symbols). Data are shown as mean from six biological replicates. Error bars are omitted for clarity. **(D)** Structure of *P. putida* RNA polymerase sigma factor RpoD (PP_0387) predicted by AlphaFold (Uniprot identifier AF-Q88QU7-F1). In PD689 tt L1, serine 552 (TCG codon) was mutated to proline (CCG codon). This residue was predicted to form the end of one of the α-helices in σ4 domain, which interacts with the –35 promoter element (Paget, 2015). RpoD structure was visualized using PyMOL v. 1.6.0.0 (Schrodinger LLC).

Transposable elements and gene duplication events are key factors in the genome evolution and environmental adaptability of *Pseudomonas species* (Silby et al., 2011). Our study, in agreement with several other recent works (Ling et al., 2022; Wirth et al., 2022), highlights the contribution of these types of genome rearrangements in the evolution of biotechnologically relevant genotypes/phenotypes of *P. putida*. In fact, such natural mechanisms could inspire new genetic tools for metabolic engineering that depart from the straight import of engineering concepts to the biological realm and rely more on long-evolved processes of emergence of new phenotypes. These results also suggest that the *gnd* deletion in EM42 Δ*gcd* Δ*gnd* pSEVA2213_*xylABE* and PD689 may not be as harmful to xylose metabolism, as originally thought. The tested Δ*gnd* strains may have had a disadvantage when compared to PD310 and PD584 due to the absence of the multiplication in their chromosomes.

The multiplication was not responsible for the improved phenotype of PD584 L3 after ALE. Other genomic changes that we identified and were able to interpret did not clearly explain the improved phenotype either. Therefore, we focused on the pSEVA2213_*xylABE* plasmid in our further search for causal mutations. The low-copy-number plasmid was included in the hybrid assembly of the genomes of strains PD584, PD584 L3, and PD689 tt L1. The only change was a perfect 32 bp duplication upstream of the *xylA* gene in PD584 L3 (**Fig. 6B**). The duplication encompassed the synthetic Shine–Dalgarno sequence (RBS), the ATG start codon and the following eight nucleotides of the *xylA* gene. Analysis of the resulting mRNA sequence with RBS Calculator (Salis et al., 2009) revealed that the RBS in the duplication became, with its predicted translation initiation rate of 310 a.u., the strongest RBS upstream of the *xylA* gene while the strength of the original *xylA* RBS was reduced 10-fold from 292 to only 29 a.u. We argue that the effect of the duplication lies in the emergence of a translational coupler – a region downstream of the promoter that encodes a short leading peptide (32 amino acids-long in case of the mutated pSEVA2213_*xylABE*) stabilizing translation of the downstream gene (Mutalik et al., 2013).

We demonstrated that the mutated pSEVA2213_*xylABE* plasmid is partially responsible for the improved growth of PD584 L3 on xylose. PD584 L3 was deprived of the plasmid by sub-culturing in rich LB medium without antibiotics and then transformed either with the same mutated plasmid or with the original intact plasmidSEVA2213_*xylABE*. Subsequent growth assay showed that the strain with the re-implanted mutated plasmid grew equally well on xylose compared to PD584 L3 (µ = 0.21 ± 0.00 h^-1^) while the strain transformed with the original plasmid grew slower by 33 % (µ = 0.14 ± 0.00 h^-1^, **Fig. 6C**). Interestingly, when we introduced the mutated plasmid from PD584 L3 into PD689 tt L1 deprived of its own pSEVA2213_*xylABE*, the resulting strain did not grow better than PD689 tt L1 (**Fig. 6C**). Moreover, PD689 tt L1 with implanted original intact pSEVA2213_*xylABE* did not show any growth retardation. These results indicate that PD689 tt L1’s improved growth on xylose can be related to the implanted *talB*-*tktA* operon and mutations in the chromosome that occurred during ALE. To verify the effect of duplication on the production of enzymes encoded by pSEVA2213_*xylABE* plasmid, we measured activities of XylA and XylB in PD584 L3, PD584, and PD689 tt L1 grown in M9 medium with xylose (the latter two strains were used as controls). The determined XylA activity in cell-free extracts prepared from PD584 L3 cells was almost twice as high as the activity measured in extracts from PD584 control (**Supplementary Fig. S11**). The activity of XylB was comparable in both strains. To our surprise, activities of both XylA and XylB were significantly higher in PD689 tt L1 than in the PD584 reference and in case of XylB even ∼20% higher than in PD584 L3. Elevated XylA and XylB activities in PD689 tt L1 cannot be linked to the sequence of pSEVA2213_*xylABE* since no changes occurred in this case. On that account, we hypothesize that the explanation could lie in the identified missense mutation in the *rpoD* gene encoding the RNA polymerase sigma factor σ70 (PP_0387). This mutation leads to quite a drastic change of a polar Ser552 to a nonpolar rigid proline in one of the α−helices of the σ4 domain in the predicted structure of RpoD (**Fig. 6D**) (Paget, 2015). The RpoD sigma factor alters binding of the RNA polymerase to different promoters by its σ4 domain interaction with –35 promoter element. Mutations in the transcription machinery and specifically in *rpo* genes were previously described in several *E. coli* strains exposed to ALE and were identified as a prominent mean of metabolism re-wiring and improved resilience against environmental perturbations, including temperature and acid stress (Choe et al., 2019; Conrad et al., 2010; Harden et al., 2015; Tenaillon et al., 2012). Choe and co-workers (2019) acknowledged the impact of mutated RpoD (Ser253→Pro253) on a specific set of promoters and corresponding transcriptome and metabolic flux re-modeling that led to improved growth of a genome-reduced *E. coli* in mineral salt medium with glucose (Choe et al., 2019). It is probable that mutations in *rpoD* genes can induce a similar effect in *P. putida* (Espeso et al., 2020) and that the mutated sigma factor together with the inserted *talB*-*tktA* cassette enabled broader adaptation of PD689 tt L1 metabolism to the non-native substrate. Importantly, the mutation may have increased the affinity of the *P. putida* RNA polymerase complex to the synthetic σ70 P_EM7_ promoter and had a positive effect on the expression of the exogenous *E. coli* genes *xylA*, *xylB*, and *xylE* (Pang et al., 2022; Tomatis et al., 2019; Tomko and Dunlop, 2017). Moreover, the absence of the 118-kb multiplication in the chromosome of PD689 tt L1, could have inferred an economic advantage due to better accessibility to cellular resources (energy, RNA polymerase molecules, nucleotides, ribosomes, tRNAs, and amino acids) that can be invested in the expression of the exogenous genes while being drained into the expression of the multiplied genes in PD584 and its descendants (Alter et al., 2021; Kim et al., 2020).

### 3.6 Quantitative proteome analyses confirm two distinct evolution scenarios towards faster utilization of xylose in PD584 L3 and PD689 tt L1

Whole-genome sequencing of *P. putida* strains suggested several causes of improved xylose metabolism. In the next step, we aimed to decipher how the proteome reflected the genomic changes. We quantitatively analyzed proteomes of the strains PD310, PD584, PD584 L3, and PD689 tt L1 grown in M9 minimal medium with 2 g L^-1^ xylose (**Supplementary methods**). Altogether, 3981 proteins (92 duplicities were removed) were uniquely detected in the four strains (**Supplementary File S6**). Individual proteomes were compared as follows: PD584 to PD310, PD584 L3 to PD584, and PD689 tt L1 to PD584 L3. The analysis showed relatively subtle differences between PD584 and PD310 (118 downregulated and 87 upregulated proteins, log_2_fold change > 1.0, P<0.05) and between the evolved strain PD584 L3 and its template PD584 (83 downregulated and 58 upregulated proteins), while the proteomes of the two evolved strains PD689 tt L1 and PD584 L3 varied in 667 proteins (376 downregulated and 292 upregulated, **Supplementary Fig. S12** and **Supplementary File S7**). Visualizing the changed protein abundances in the map of *P. putida* central carbon metabolism verified the effects of our targeted engineering interventions and ALE as well (**Fig. 7**). Note that proteins with less prominent changes in abundance (log_2_FC ≥ 0.5 or ≤ –0.5, P<0.05) were also visualized since relatively subtle changes in enzyme quantity can have pronounced effects in the metabolic network.

**Figure 7.**
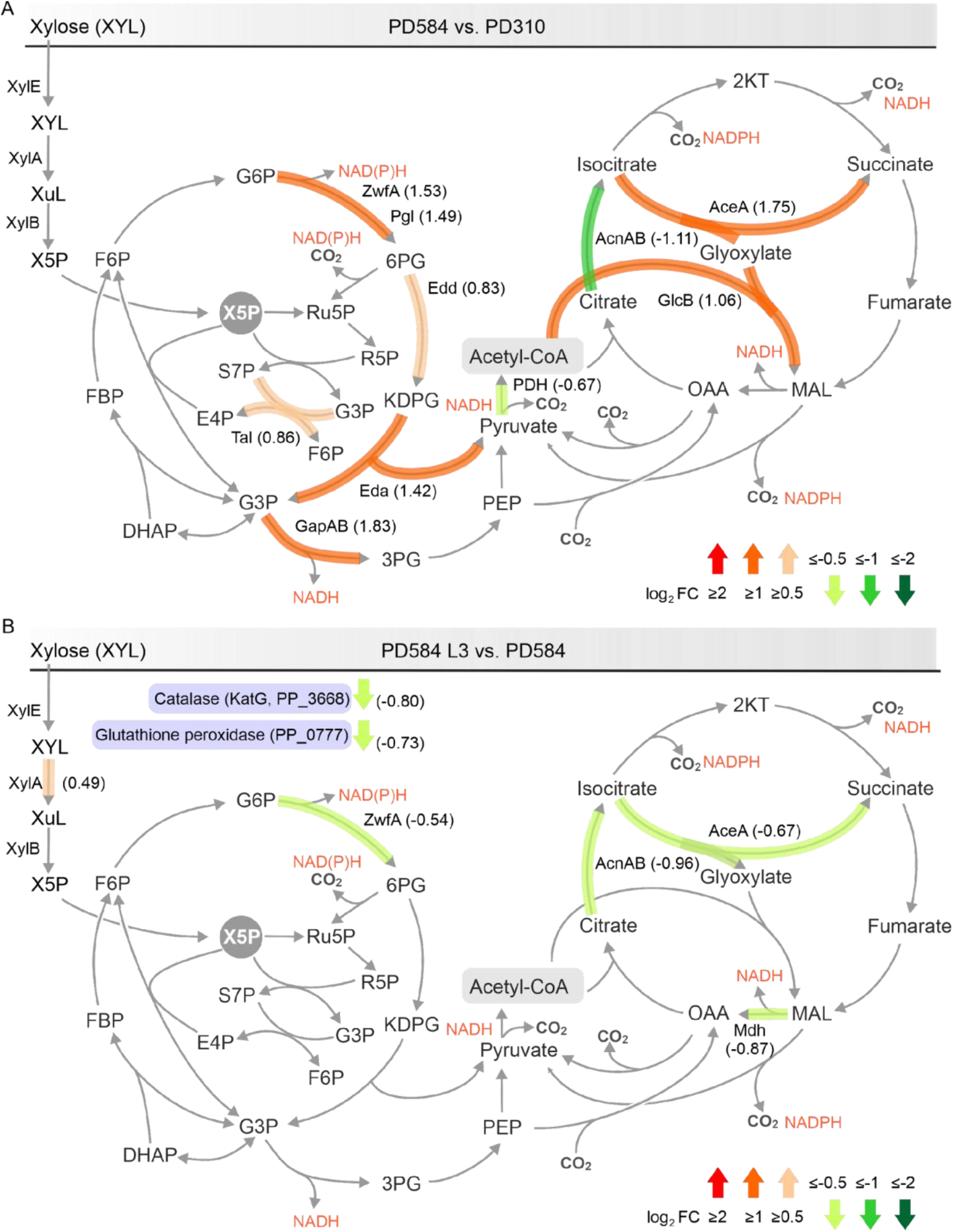
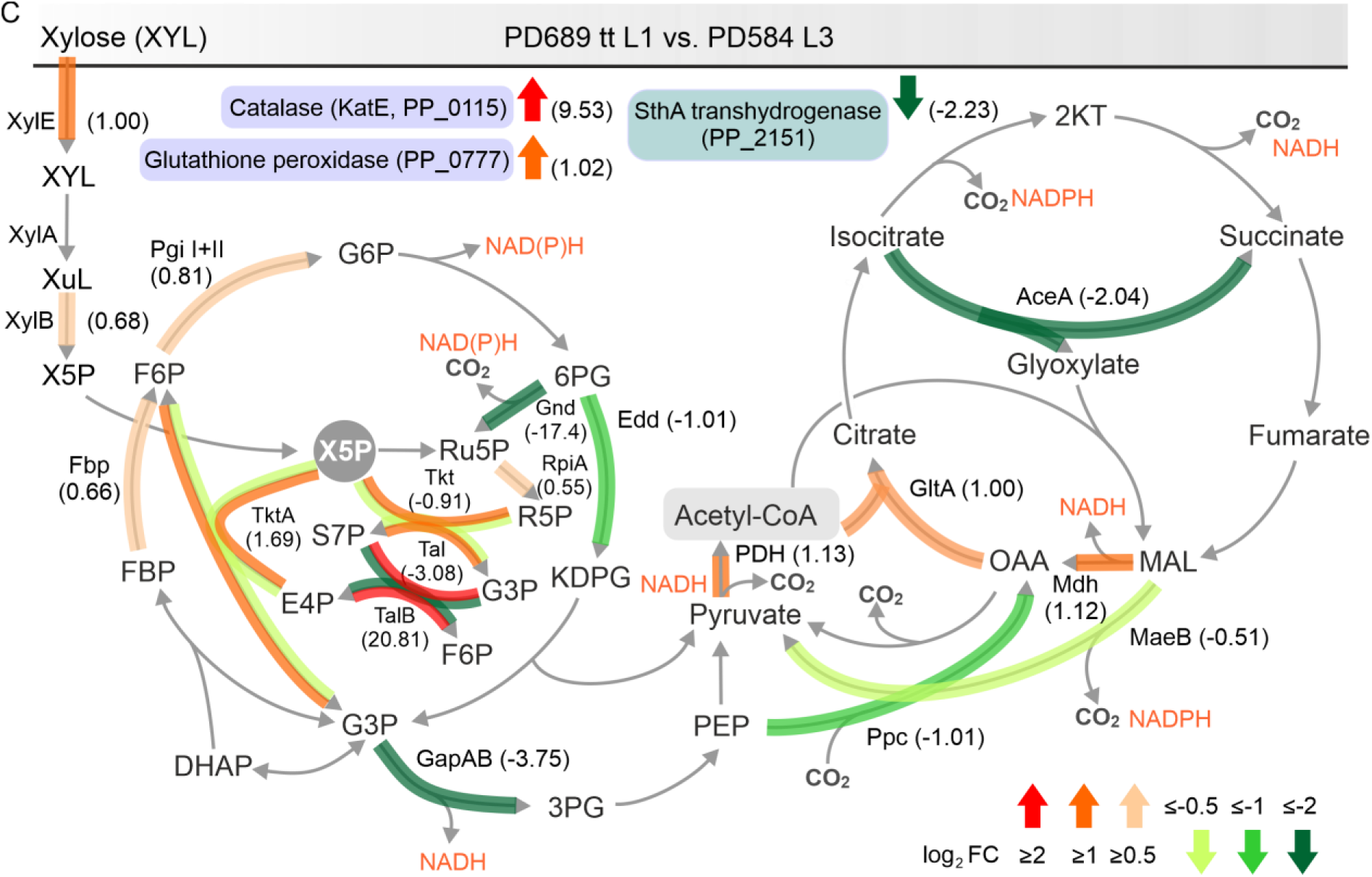
Changes in protein abundances in the upper xylose pathway and central carbon metabolism for strain **(A)** PD584 compared to PD310, **(B)** PD584 L3 compared to PD584, and **(C)** PD689 tt L1 compared to PD584 L3. The figures show log_2_(fold change, FC) values for significantly differentially expressed proteins (p ≤ 0.05). The metabolic map and abbreviations used are the same as in Figure 2A. Please note that Tal and Tkt represent native *P. putida* transaldolase (PP_2168) and transketolase (PP_4965), respectively, while TalB (KEGG ID: JW0007) and TktA (KEGG ID: JW5478) stand for the respective enzymes from *Escherichia coli*. SthA is soluble pyridine nucleotide transhydrogenase.

In the first comparison (PD584 vs. PD310), we could see the expected effect of the *hexR* deletion in PD584 manifested as increased abundances of the enzymes ZwfA, Pgl, Edd, Eda, and Gap (**Fig. 7A**). Higher activity of de-repressed dehydrogenases Zwf and Gap could exacerbate redox disbalance and lead to the apparent upregulation of isocitrate lyase AceA (PP_4116) and malate synthase GlcB (PP_0356) (**Fig. 7A**) of the glyoxylate shunt. Another upregulated enzyme was the native Tal transaldolase (PP_2168) (**Fig. 7A, Supplementary File S7**). Its modestly increased abundance could be attributed either to the *hexR* deletion (Bentley et al., 2020), to a higher number of copies of the PP_2114—PP_2219 segment in the chromosome of PD584 (six copies) when compared to PD310 (four copies, **Supplementary Table S4**), or to both these factors.

Inspection of downregulated and upregulated proteins in the central carbon metabolism of PD584 L3 compared to PD584 revealed only minute changes including a small decrease in the quantity of ZwfA in the EDEMP cycle, AceA in the glyoxylate shunt, or malate dehydrogenase Mdh (PP_0654) in the TCA cycle (**Fig. 7B**). Increase in the abundance of exogenous XylA, which most probably reflects the identified duplication upstream of the *xylA* gene in pSEVA2213_*xylABE* plasmid, was also observed. This data supports our hypothesis that in the lineage of strains PD310, PD584, and PD584 L3 the multiplication of the genomic segment PP_2114—PP_2219 and the de-repression of a part of the EDEMP cycle by the *hexR* deletion were key de-bottlenecking events, which facilitated further adaptation of the bacterium to xylose. In PD584 L3, further improvement was enabled by a slightly increased amount of XylA and additional changes outside the central carbon metabolism that may have led to better fitness.

The increased vigour of PD584 L3 compared to PD584 is suggested by decreased abundances of several universal stress proteins (PP_3294, PP_2326, and PP_3156; log_2_FC = –8.55, –1.09, and –0.73, respectively) and enzymes responsible for ROS elimination and oxidative stress mitigation – catalase KatG and glutathione peroxidase (**Fig. 7B**).

Comparison of PD689 tt L1 and PD584 L3 proteomes revealed many intriguing variances and confirmed that the former strain with the deleted 6-phosphogluconate dehydrogenase gene *gnd* followed a different evolutionary path to the improved xylose utilization phenotype (**Fig. 7C**, **Supplementary Fig. S12**, and **Supplementary File S7**). The proteomic data also supported the hypothesis that flux through the Gnd reaction is linked to the activity of the glyoxylate shunt (Beckers et al., 2016; Meijnen et al., 2012). Isocitrate lyase AceA was significantly downregulated in *gnd* negative PD689 tt L1 (**Fig. 7C, Supplementary Fig. S5B**). Upregulation of pyruvate dehydrogenase complex, citrate synthase GltA (PP_4194), and malate dehydrogenase Mdh probably supported functioning of the TCA cycle including its reductive part between isocitrate and succinate and could compensate for the likely decrease in the level of reducing co-factors due to the *gnd* deletion and Gap downregulation.

We presumed that some of the 376 downregulated proteins in PD689 tt L1 were encoded by genes from the chromosomal region (PP_2114—PP_2219), which was multiplied in PD584 L3 but not in PD689 tt L1. Indeed, we identified 36 proteins including 23 enzymes encoded in this region that were significantly downregulated (P<0.05, log_2_FC < –1.0) in PD689 tt L1 compared to PD584 L3 (**Supplementary File S8**). Not surprisingly, this protein set included Tal transaldolase, GapB glyceraldehyde 3-phosphate dehydrogenase, and SthA soluble pyridine nucleotide transhydrogenase. Absolute quantification of these 36 proteins in PD689 tt L1, PD584, and PD310 showed their higher levels in the latter two strains (**Supplementary File S8**). This result was consistent with the presence of the multiplication in the chromosome of both PD584 and PD310. Proteomic analysis also verified the presence of the exogenous transaldolase TalB and transketolase TktA in PD689 tt L1 (**Fig. 7C**). Native Tal and Tkt were downregulated compared to PD584 L3. Rpe and RpiA enzymes showed no or modest change in quantity. We argue that in contrast to PD310, PD584, and PD584 L3, which benefited from several copies of the native *tal* gene in the multiplied chromosome segment and which had active carbon recycling through the Gnd reactrion, the function of non-oxidative PPP in *gnd*^-^ mutant PD689 tt L1 was enhanced by the enzymes transplanted from *E. coli*. It seems that transaldolase is a key PPP enzyme whose activity is one of the major determinants of xylose utilization rate in *P. putida*. This hypothesis is supported by the measurements of transaldolase activity in six *P. putida* strains discussed in this study (**Supplementary Fig. S13**). The observed trend is that the faster the growth on xylose the higher the detected transaldolase activity, with PD584 L3 and PD689 tt L1 showing the highest values.

Another important element is the expression level of *xylA*, *xylB*, and *xylE* genes of the upper xylose pathway. The abundance of XylA, same as its activity, was very similar in PD689 tt L1 and in PD584 L3 with the duplication upstream of the *xylA* gene (**Figs. 7C and S11**). On the contrary, XylB and XylE were upregulated in PD689 tt L1. Modest upregulation of XylB correlated with its higher activity in PD689 tt L1 (**Fig. S11**). In the study of Elmore and coworkers (2020), higher expression of the *xylE* gene due to a mutation in its promoter was the major determinant of improved xylose utilization by engineered *P. putida* KT2440 (Elmore et al., 2020). But in our case, no mutation was pinpointed in the pSEVA2213_*xylABE* plasmid isolated from PD689 tt L1. Upregulation of XylE, XylA, and XylB in this strain thus signifies a higher expression of the whole synthetic *xylABE* operon. This could be caused by the enhanced *xylABE* transcription due to the Ser552→Pro552 mutation in RNA polymerase sigma factor RpoD discussed earlier. Higher expression of the whole *xylABE* operon and the adoption of exogenous TalB and TktA enzymes do not need to have only positive effects on PD689 tt L1. The substantially increased quantity of ROS-reducing enzymes – catalase KatE (PP_0115) and glutathione peroxidase (PP_0777) in PD689 tt L1 when compared to PD584 L3 (**Fig. 7C**) may reflect metabolic “discomfort” of the former strain. The stress can stem from the overproduction of the exogenous transporter XylE (Wagner et al., 2007) or the implanted *E. coli* pathways. These routes can burden the host by the overexpression of respective genes or by the increased fluxes through the non-oxidative PPP that are common in *E. coli* but unusual in *P. putida* (Elmore et al., 2020; Rojo, 2010; Wirth et al., 2022).

## 4 Conclusions

Upon mapping the xylose assimilation in our original engineered strain PD310 (Dvořák and de Lorenzo, 2018), we identified the partially cyclic character of the upper xylose metabolism. Mathematical modeling and gene knockout experiments showed that this carbon cycling through Gnd reaction is not essential for xylose utilization by *P. putida*. In agreement with Meijnen and co-workers (2012), we also described the negative effect of the HexR regulator on upper xylose metabolism in *P. putida* (Meijnen et al., 2012). Other recent studies on *P. putida* KT2440 with installed isomerase pathway do not discuss the HexR-mediated regulation as a bottleneck for xylose utilization (Elmore et al., 2020; Ling et al., 2022; Wang et al., 2019). But our results demonstrate that the *hexR* knockout should be employed to reduce the lag phase on xylose. What remains to be elucidated is the molecular cause of slower or less efficient de-repression of HexR-regulated operons in xylose-grown cells when compared with cells cultured on glucose.

Another important finding of this work is that *P. putida xylABE*^+^ does not need to be equipped with exogenous PPP genes but it requires some PPP flux enhancement and balancing to accelerate its growth on xylose. The bacterium with the functional carbon cycling via Gnd took advantage of the mobile elements in its genome and multiplied a large segment of its chromosome with genes beneficial to xylose metabolism. One of these multiplied and upregulated genes was a transaldolase whose increased activity in PD310, PD584, and PD584 L3, compared with freshly prepared *P. putida* EM42Δ*gcd* pSEVA2213_*xylABE* (**Supplementary Fig. S13**), correlated with the presence of 118-kb multiplication in the chromosome of these strains. On the other hand, PD689 tt L1 was likely pushed to employ the exogenous *talB* gene because its parental strain PD689 lacked the genomic multiplication and the Gnd enzyme, which helps replenish pools of ribulose 5-phosphate and ribose 5-phosphate. In mutants with *gnd* removed based on the computer-aided design, the absence of carbon cycling had to be compensated. The implanted TalB could provide the necessary “pull” effect to promote a flow of carbon through the preceding non-oxidative PPP reactions (Volke 2022). In the strains constructed and evolved in this study (PD584 L3 and PD689 tt L1), upregulation of transaldolase was pivotal for faster xylose utilization.

Other essential constituents of the engineered xylose metabolism in *P. putida* EM42 are the *E. coli* genes in the synthetic *xylABE* operon. Expression of these genes became a limiting step for xylose utilization by *P. putida* EM42 only when the aforementioned bottlenecks in the core carbohydrate metabolism were removed. Both strains, PD584 L3 and PD689 tt L1, that emerged from the ALE experiment demonstrated a different strategy for balancing gene expression in the *xylABE* operon. Strain PD584 L3 finetuned the expression of *xylA* by embossing the gene with a translational coupler. Together with additional subtle changes in its genome and proteome, this helped improve the growth of PD584 L3 on xylose. In contrast, the adaptation of PD689 tt L1 occurred through a more “brute-force” approach – overexpression of the whole synthetic operon – probably enabled by a missense mutation in the sigma factor RpoD. The resulting *xylABE* overexpression and transcriptome re-modeling, translated in numerous changes on the proteome level, accelerated the growth of PD689 tt L1 on xylose but contributed to the reduced strain’s robustness.

In conclusion, our study describes how *P. putida* benefits from its remarkable genomic and metabolic plasticity when adapting to a non-native substrate. In this particular case, the adaptation occurred through rational engineering cuts combined with evolutionary events including a large genome re-arrangement, a smaller duplication, or single nucleotide substitutions. The optimal nesting of the new route in the pre-existing biochemical and physiological network of the host required substantial as well as fine adjustments of the starting metabolic and regulatory devices that could not be thoroughly addressed through rational calculations but were, to some extent, knowledge-guided (Porcar et al., 2015). As a result, the best growers PD584 L3 and PD689 tt L1 represent discrete local optima on the fitness landscape, which were reached through different evolutionary scenarios. The selection of a suitable microbial host is a critical first step for the success of any bioengineering project employing ALE (Sandberg et al., 2019). In agreement with other recently published works, our results suggest that *P. putida* KT2440 and its derivatives are attractive templates for evolutionary experiments that can be directed by knowledge-guided design (Ling et al., 2022; Wirth et al., 2022). Such an approach adopted here provided *P. putida* strains with a doubled growth rate and substantially reduced growth lag on xylose, which make them promising templates for the biotechnological valorization of xylose or its co-valorization with glucose. The findings presented in this study are instrumental for future attempts to exploit semi-synthetic xylose metabolism in *P. putida* KT2440 and its derivatives for biotechnological purposes as well as for the understanding of bacterial adaptation to new carbohydrate substrates.

### CRediT authorship contribution statement

**Pavel Dvořák** Conceptualization, Investigation, Resources, Data curation, Methodology, Validation, Visualization, Funding acquisition, Supervision, Project administration, Writing – original draft, review & editing. **Barbora Burýšková** Conceptualization, Investigation, Methodology, Validation, Data curation, Visualization, Writing – original draft, Writing – review & editing. **Barbora Popelářová** Conceptualization, Investigation, Methodology, Validation, Data curation, Visualization, Writing – original draft, Writing – review & editing. **Birgitta E. Ebert** Conceptualization, Investigation, Methodology, Validation, Data curation, Writing – review & editing. **Tibor Botka** Investigation, Methodology, Data curation, Visualization, Writing – review & editing. **Dalimil Bujdoš** Methodology, Data curation, Writing – review & editing. **Alberto Sánchez-Pascuala** Methodology, Data curation, Writing – review & editing. **Hannah Schöttler** Methodology, Data curation. **Heiko Hayen** Methodology, Data curation. **Víctor de Lorenzo** Resources, Writing – review & editing. **Lars M. Blank** Resources, Writing – review & editing. **Martin Benešík** Conceptualization, Investigation, Data curation, Methodology, Validation, Visualization, Writing – review & editing.

### Data availability statement

All genome sequences and whole-genome sequencing data are available in GenBank under accession codes listed in Supplementary Table S3. Supplementary data to this manuscript are available in separate files.

## Acknowledgements

We thank Dr. Adam Feist and Dr. Hyungyu Lim for valuable discussions over genomic data of engineered and evolved *P. putida* strains. This work was funded by Czech Science Foundation Project 22-12505S and Grant Agency of Masaryk University GAMU Project MASH Junior 2022 granted to P.D. and Brno Ph.D. Talent granted to B.B. We acknowledge CF CF Prot of CIISB, Instruct-CZ Centre, supported by MEYS CR (LM2018127) and European Regional Development Fund-Project „UP CIISB“ (No. CZ.02.1.01/0.0/0.0/18_046/0015974). This work was supported by the project National Institute of Virology and Bacteriology (Programme EXCELES, ID Project No. LX22NPO5103), funded by the European Union – Next Generation EU.

