## Supplementary material for "Genomic and metabolic plasticity drive alternative scenarios for adapting *Pseudomonas putida* to non-native substrate D-xylose"

by

Pavel Dvořák[1§*], Barbora Burýšková[1§], Barbora Popelářová[1§], Birgitta Ebert[2§], Tibor Botka[1], Dalimil Bujdoš[1,#], Alberto Sánchez-Pascuala[3], Hannah Schöttler[4], Heiko Hayen[4], Víctor de Lorenzo[5], Lars M. Blank[6], and Martin Benešík[1]

[1]*Department of Experimental Biology, Faculty of Science, Masaryk University, Kamenice 753/5, 62500, Brno, Czech Republic.*

[2]*Australian Institute for Bioengineering and Nanotechnology, University of Queensland, Cnr College Rd & Cooper Rd, St Lucia QLD 4072, Australia.*

[3]*Department of Biochemistry and Synthetic Metabolism, Max Planck Institute for Terrestrial Microbiology, Karl-von-Frisch-Straße 10, 35043 Marburg, Germany.*

[4]*Institute of Inorganic and Analytical Chemistry, University of Münster, Corrensstraße 48, 48149 Münster, Germany.*

[5]*Systems and Synthetic Biology Program, Centro Nacional de Biotecnología CNB-CSIC, Cantoblanco, Darwin 3, 28049 Madrid, Spain.*

[6]*Institute of Applied Microbiology, RWTH Aachen University, Worringer Weg 1, 52074 Aachen, Germany.*

[§]Shared first authors

[#]Current address: Food Science Building, College Rd, University College, Cork, Ireland.

*** Corresponding author:**

Dr. Pavel Dvořák

Department of Experimental Biology (Section of Microbiology, Microbial Bioengineering Laboratory)

Faculty of Science, Masaryk University

Kamenice 735/5

Brno 62500, Czech Republic

https://mik.sci.muni.cz/mbl

### Content

***Supplementary methods***

***Supplementary results***

***Supplementary tables***

**Table S1.** Plasmids used in this study.

**Table S2.** Oligonucleotide primers used in this study.

**Table S3.** Accessions to sequencing data of *Pseudomonas putida* strains in this work in NCBI databases.

**Table S4.** Presence of multiplied genomic locus PP_2114 - PP_2219 in the chromosome of sequenced *Pseudomonas putida* strains.

***Supplementary figures***

**Figure S1.** Shake flask cultures of *Pseudomonas putida* PD310 in M9 minimal medium with 5 g $L^{-1}$ glucose or xylose.

**Figure S2.** Comparison of growth rate and specific xylose uptake rate calculated using MFA and FBA metabolic model.

**Figure S3.** Shake flask cultures of *Pseudomonas putida* PD584 or *P. putida* EM42 Δ*hexR* pSEVA2213_*xylABE* in M9 minimal medium with xylose (5 g $L^{-1}$).

**Figure S4.** Growth of $hexR^{+}$ PD310 and $hexR^{-}$ PD584 in M9 minimal medium with 2 g $L^{-1}$ D-glucose or 2 g $L^{-1}$ D-fructose in 48-well microtiter plate.

**Figure S5.** Shake flask cultures of *Pseudomonas putida* PD689 in M9 minimal medium with xylose (2 g $L^{-1}$) and specific activity of 6-phosphogluconate dehydrogenase Gnd determined in cell-free extracts of PD584, PD689, and PD689 tt L1.

**Figure S6.** Screening of selected evolved mutants of *P. putida* PD584 and PD689 strains expressing heterologous pentose phosphate pathway genes on xylose in M9 minimal medium with 2 g $L^{-1}$ xylose in 48-well microtiter plate or shake flasks with 5 g $L^{-1}$ xylose.

**Figure S7.** Adaptive laboratory evolution on xylose of *P. putida* PD310 strain cultured in shake flasks containing 20 mL of M9 minimal salts medium, 5 g $L^{-1}$ D-xylose and kanamycin and passaged every 48 h.

**Figure S8.** Shake flask cultures of *P. putida* PD584 L3, PD584 tt L3, and PD689 tt L1 in M9 minimal medium with xylose (5 g $L^{-1}$).

**Figure S9.** Shake flask cultures of *P. putida* PD584 L3 and PD689 tt L1 in M9 minimal medium with xylose and glucose (2 g $L^{-1}$ each).

**Figure S10.** Growth of *P. putida* PD310 and *P. putida* EM42 Δ*gcd* pSEVA2213_*xylABE* in 48-well microtiter plate with M9 minimal salts medium and 2 g $L^{-1}$ xylose.

**Figure S11**. The specific activity of xylose isomerase XylA (left graph) and xylulokinase XylB (right graph) determined in cell-free extracts of four *P. putida* strains.

**Figure S12**. Volcano plots depicting differentially abundant proteins in strain PD584 compared to PD310, PD584 L3 compared to PD584, and PD689 tt L1 compared to PD584 L3.

**Figure S13.** The specific activity of transaldolase determined in cell-free extracts of six *P. putida* strains.

***Nucleotide sequences of genes used in this study***

***References***

### Supplementary methods

#### General cloning procedures

Plasmid DNA was routinely isolated using GeneJET Plasmid Miniprep Kit (Thermo Fisher Scientific), or E.Z.N.A. Plasmid DNA Mini Kit I (Omega Bio-Tek). Genomic DNA was isolated using RTP Bacteria DNA Mini Kit (INVITEK Molecular). The genes of interest were amplified by polymerase chain reaction (PCR) using Q5 high fidelity DNA polymerase (New England BioLabs) according to the manufacturer's protocol. The reaction mixture (50 μL) further contained polymerase HF buffer or GC buffer in case of a template with a high GC content (New England BioLabs), dNTPs mix (0.2 mM each; Roche), respective primers (0.5 mM each), water, and template DNA. Colony PCR was performed in a 10 μL volume using 2x DreamTaq Green PCR Master Mix (Thermo Scientific) with oligonucleotide primers (0.5 μM each) for strain verification and confirmation of inserts in plasmids. All PCR reactions were carried out in Labcycler Gradient (SensoQuest). All used restriction enzymes were from New England BioLabs. Digested DNA fragments were ligated in a 10 μL reaction using T4 DNA ligase (New England BioLabs) at 16 °C overnight or at RT for 15 min according to the manufacturer´s instructions. PCR products and digested plasmids separated by DNA electrophoresis with 0.8 % (w/v) agarose gels were compared to Quick-Load 1 kb DNA Ladder (New England BioLabs) and visualized using G:Box XT4 Digital Imaging System (Syngene). DNA was purified from PCR mixtures or agarose gels using NucleoSpin Gel and PCR Clean-up (Macherey-Nagel). The purity and concentration of DNA were determined by NanoDrop 2000 (Thermo Fisher Scientific). If needed, DNA was concentrated using DNA 120 SpeedVac Concentrator (Thermo Fisher Scientific). Chemocompetent *E. coli* CC118, DH5α, or CC118λpir cells were transformed with ligation mixtures or plasmid constructs, and individual clones selected on LB agar plates with an antibiotic were used for the preparation of cryogenic glycerol (20% v/v in LB medium) stocks. Plasmid constructs were sequenced by Eurofins Genomics or SEQme Czech Republic. Plasmids were inserted into *P. putida* EM42 by electroporation (voltage of 2.5 kV, capacitance of 25 μF, resistance of 200 Ω) in a 2 mm gap cuvette (Thermo Fisher Scientific) using GenePulser XcellTM (Bio-Rad). The preparation of *P. putida* electrocompetent cells and the electroporation procedure were performed as described elsewhere (Martínez-García and de Lorenzo, 2012). After the electric pulse, 0.9 mL of LB medium was added to the cell suspension (100 µL) in the cuvette and the mixture was transferred to a 2 mL plastic tube. Cells were incubated for 2 h in case of electroporating pSEVA2213, pEMG, and pSW-I plasmids or 5 h in case of pBAMD constructs (200 RPM, 30 °C, NB205 incubator). Alternatively, plasmid constructs were transferred from *E. coli* donors to *P. putida* EM42 by triparental mating, using *E. coli* HB101 helper strain with pRK600 plasmid (**Table 1 in the manuscript**). After electroporation or mating, the cells were plated on selective LB agar plates with an antibiotic, incubated at 30 °C overnight and single colonies of transformants or transconjugants were then re-streaked twice on fresh LB plates with antibiotic and the presence of a plasmid was verified by colony PCR and restriction analysis.

**Enzyme assays**

For enzyme activity measurements in *P. putida* EM42 strains, cell lysates were prepared by lysing cells cultured in 50 mL of LB medium with kanamycin or 50 mL of M9 medium with 2 g $L^{-1}$ xylose or glucose and kanamycin. These cultures were inoculated to a starting $OD_{600}$ of 0.05 from night cultures (grown for 16 h) in 10 mL of LB medium with kanamycin and grown while shaking (200 rpm, IS-971R, Jeio Tech) at 30°C. Cells were collected in mid log phase ($OD_{600}$ = 0.5) and the whole culture was spun down (2,000 g, 4°C, 15 min). Cells were washed 2x in ice-cold 100 mM potassium phosphate buffer of pH 7.1 and the pellets were lysed with 200 uL of B-PER Bacterial Protein Extraction Reagent, 0.2 µL of DNAse I (2.6 U $\mu L^{-1}$), and 0.2 µL lysozyme (50 mg $mL^{-1}$) (all Thermo Scientific) for 15 min at RT with slow agitation. Cell lysates were centrifuged at 21,000 g for 30 min at 4°C and supernatants, termed here as cell-free extracts (CFE), were used for activity determination or stored at -80°C for repeated measurements. Total protein concentration in CFE was measured using the method of Bradford (Bradford, 1976) with a commercial kit (Sigma-Aldrich). Crystalline bovine serum albumin (Sigma-Aldrich) was used as a protein standard.

All accessory enzymes and the majority of chemicals used in the assays described below were purchased from Sigma-Aldrich.

All enzymatic activities were measured in 96-well microtiter plate format. Activities of xylose isomerase (XylA) and xylulokinase (XylB) were measured as described previously (Dvořák and de Lorenzo, 2018). In the XylA assay, activity is coupled to the consumption of NADH by sorbitol dehydrogenase. The assay mixture contained (final concentrations are denoted in all following assays): 1 mM triethanolamine, 0.5 mM NADH, 0.5 U of sorbitol dehydrogenase, 10 mM $MgSO_4$, and 50 mM D-xylose, and 2.5 µL of CFE, all supplemented with 50 mM Tris-HCl buffer (pH 7.5) to a final volume of 200 µL. The reaction mixture was heated to 30°C, and the reaction started with the addition of NADH. In the XylB assay, activity is coupled to pyruvate kinase and lactate dehydrogenase leading to the consumption of NADH. The reaction mixture contained: 0.5 mM NADH, 2 mM ATP, 2 mM $MgCl_2$, 0.2 mM phosphoenolpyruvate, 10 U of pyruvate kinase, 10 U of lactate dehydrogenase, 10 mM D-xylulose, and 2 µL of diluted CFE, adjusted with 50 mM Tris-HCl buffer (pH 7.5) to a final volume of 200 µL. The mixture was heated to 30°C and the reaction started with the addition of NADH.

Activities of 6-phosphogluconate dehydratase (Edd) and 2-keto-3-deoxy-6-phosphogluconate aldolase (Eda) were measured in a combined assay based on the protocol of Stephenson et al. (Stephenson et al., n.d.). The assay measures pyruvate production from 6-phosphogluconate using lactate dehydrogenase and NADH. The reaction mixture contained: 5 mM dithiothreitol, 0.25 mM $MnCl_2$, 1 U of L-lactate dehydrogenase, 0.5 mM NADH, 3.1 mM 6-phosphogluconate, and 6 µL of CFE, adjusted with 100 mM triethanolamine·HCl buffer (pH 7.6) to a final volume of 200 µL. The mixture was heated to 30°C and the reaction started with the addition of 6-phosphogluconate.

Activity of glucokinase (Glk) was measured as described by Sánchez-Pascuala et al. (2019) with some modifications (Sánchez-Pascuala et al., 2019). The reaction mixture contained: 67 µL of 120 mM Tris·HCl buffer (pH = 8.2), 50 mM D-glucose, 4 mM $MgCl_2$, 10 mM ATP, 0.5 mM $NADP^+$, 1 U of glucose-6-P dehydrogenase, 2 µL of CFE, adjusted with water to a final volume of 200 µL. The mixture was heated to 30°C and the reaction started with the addition of $NADP^+$.

The activity of glucose 6-phosphate isomerase (Pgi) was measured following the protocol of Sánchez-Pascuala et al. (2017) with some modifications (Sánchez-Pascuala et al., 2017). The reaction mixture contained: 66 µl of 100 mM glycylglycine buffer (pH = 7.5), 4 mM D-fructose 6-phosphate, 0.5 mM $NADP^+$, 4 mM $MgCl_2$, 1.5 U of glucose-6-phosphate dehydrogenase, and 1 µL of CFE, adjusted with water to a final volume of 200 µL. The mixture was heated to 30°C and the reaction started with the addition of $NADP^+$.

The activity of triose phosphate isomerase (Tpi) was measured following the protocol of Sánchez-Pascuala et al. (2017) with some modifications (Sánchez-Pascuala et al., 2017). The reaction mixture contained: 1 U of glycerol-3-phosphate dehydrogenase, 2.5 mM D,L-glyceraldehyde 3-phosphate, 0.5 mM NADH, and 2 µl of 20x diluted CFE, adjusted with 100 mM triethanolamine buffer (pH = 7.6) to a final volume of 200 µL. The mixture was heated to 30°C and the reaction started with the addition of NADH.

The activity of glucose 6-phosphate dehydrogenase (Zwf) was measured following the protocol of Sánchez-Pascuala et al. (2017) with some modifications (Sánchez-Pascuala et al., 2017). The reaction mixture contained: 1 mM D-glucose 6-phosphate, 5 mM $MgCl_2$, 0.5 mM $NADP^+$, and 2 µL of CFE, adjusted with 100 mM triethanolamine buffer (pH 7.6) to a final volume of 200 µL. The mixture was heated to 30°C and the reaction started with the addition of $NADP^+$.

The activity of transaldolase (Tal) was measured based on the protocol described by Zhu et al. (2017) with minor modifications (Zhu et al., 2017). The reaction mixture contained: 7.5 mM fructose 6-phosphate, 0.75 mM erythrose 4-phosphate, 1 U of glycerol-3-phosphate dehydrogenase, 2 U of triose phosphate isomerase, 0.5 mM NADH, and 2 µL of CFE, adjusted with 100 mM glycylglycine buffer (pH 7.5) to a final volume of 200 µL. The mixture was heated to 30°C and the reaction started with the addition of NADH.

The activity of transketolase (Tkt) was measured based on the protocol of Sobota and Imlay (Sobota and Imlay, 2011). The reaction mixture contained: 0.3 mM thiamine pyrophosphate, 1 U of glycerol-3-phosphate dehydrogenase, 10 U triose phosphate isomerase, 1 mM xylulose 5-phosphate, 1 mM ribose 5-phosphate, 0.5 mM NADH, and 5 µL of CFE, adjusted with 100 mM glycylglycine buffer (pH 7.5) to a final volume of 200 µL. The mixture was heated to 30°C and the reaction started with the addition of NADH.

The activity of 6-phosphogluconate dehydrogenase (Gnd) was measured following the protocol of Sánchez-Pascuala et al. (2017) with some modifications (Sánchez-Pascuala et al., 2017). The reaction mixture contained: 2 mM D-gluconate 6-phosphate, 1 mM $NADP^+$, and 2.5 µL of CFE, adjusted with 100 mM glycylglycine buffer (pH 7.5) to a final volume of 200 µL. The mixture was heated to 30°C and the reaction started with the addition of $NADP^+$.

The oxidation of NADH (decrease in $A_{340}$) or the reduction of $NADP^+$ (increase in $A_{340}$) was measured spectrophotometrically at 340 nm with Infinite M Plex plate reader (Tecan). A molar extinction coefficient of 6.22 $mM^{-1}$ $cm^{-1}$, representing the difference between the extinction coefficients of NAD(P)H and $NAD(P)^+$, was used for activity calculations. 1 unit (U) of activity corresponds to 1 µmol (or 1 nmol in case of Gnd) of a substrate (NADH, NADPH) converted by 1 mg of enzyme per/in 1 min.

#### Analysis of polar metabolites by capillary ion chromatography-mass spectrometry (IC-MS)

***Chemicals and Materials*.** Acetonitrile in LC-MS grade was obtained from VWR International GmbH (Darmstadt, Germany). All other reagents were obtained from Sigma-Aldrich (Steinheim, Germany). Purified water from a Milli-Q®-Academic system (Millipore, Molsheim, France) was used throughout the sample preparation and the measurements. The standards utilized for method development are listed below.

**List of metabolite standards, their abbreviations and purities.**

| Compound | Abbreviation | Purity |
|---|---|---|
| pyruvic acid sodium salt | pyruvate | ≥ 98 % |
| disodium succinate | succinate | ≥ 96 % |
| DL-malic acid | malate | ≥ 99 % |
| sodium fumarate dibasic | fumarate | ≥ 99 % |
| citric acid monohydrate | citrate | ≥ 99 % |
| DL-isocitric acid trisodium salt | isocitrate | ≥ 93 % |
| $\alpha$-D-glucose 1-phosphate disodium hydrate | G1P | ≥ 97 % |
| D-glucose 6-phosphate sodium salt | G6P | ≥ 99 % |
| fructose-6-phosphate disodium salt hydrate | F6P | ≥ 98 % |
| D-ribose 5-phosphate disodium salt hydrate | R5P | ≥ 98 % (TLC) |
| 6-phosphogluconic acid trisodosium salt | 6PG | ≥ 95 % |
| D-(-)-3-phosphoglyceric acid disodium salt | G3P | ≥ 93 % |
| D-fructose 1,6-bisphosphate trisodium salt hydrate | FBP | ≥ 97 % (TLC) |

Beside the pyruvic acid standard, which was purchased from Merck KgaA (Darmstadt, Germany), all metabolite standards were obtained from Sigma Aldrich Chemie GmbH (Steinheim, Germany).

The samples were reconstituted in purified water immediately before analysis.

***Capillary IC-MS analysis.*** Capillary IC-MS analysis was performed using a Dionex™ ICS-4000 Capillary HPIC™ system connected to a Q Exactive™ Plus (Thermo Fisher Scientific, Dreieich, Germany) utilizing a HESI-II electrospray ionization (ESI) source. Sample injections were performed by a Dionex™ AS-AP autosampler. For delivery of a regeneration water flow and a make-up solution flow, the system was equipped with two external AXP™ Auxiliary pumps from Thermo Fisher Scientific (Dreieich, Germany).

Samples were separated on a Dionex IonPac AS11-HC-4µM column (250 x 0.4 mm, 4 µm; Thermo Fisher Scientific) that was maintained at 35°C. The flow rate was set to 17 µL/min and the injection volume was 0.4 µL. The KOH-gradient program utilized for the separation was: 1 mmol/L KOH held for 2 min, increased to 15 mmol/L at 8 min, 20 mmol/L at 12 min, 30 mmol/L at 22 min, 70 mmol/L at 37 min held for 3 min and finally increased to 100 mmol/L at 41 min held for 2 min followed by a 2 min decrease back to initial conditions, which was held for 5 min. The total analysis time was 50 min. For an improved ionization, an acetonitrile/water solution (1:1) containing 0.1 Vol.-% ammonium hydroxide was delivered as make-up flow at a flow rate of 30 µL/min and combined with the eluent via a low dead volume mixing tee, and passed through a grounding union before entering the ESI source.

The Q Exactive™ mass spectrometer was operated in negative ionization mode and the spray voltage was set to 2.8 kV. The capillary temperature was set to 250 °C, the sheath gas flow rate was 25 (arbitrary units), the auxiliary gas flow rate was 8 (arbitrary units), the sweep gas flow rate was 0 (arbitrary units), and the S-lens level was set to 50.

The samples were measured in targeted selected ion monitoring (tSIM) mode with the following parameters for the analysis of the cell extracts: resolution, 140,000 (at *m*/*z* 200); auto gain control target, 1 x $10^5$; maximum ion injection time, 100 ms; and an isolation window with the mass range of 7 *m*/*z* to include the $^{13}$C-labelled isotopes. The exact masses for the unlabelled metabolites and the measuring time for each tSIM-window were experimentally evaluated with authentic standards. To include isotopes with multiple incorporated $^{13}$C-atoms and to centre the tSIMs' mass range on the $^{13}$C-labelled isotopes, the measured accurate masses from the authentic standards were expanded by 3 *m*/*z* for the inclusion list. The targeted masses and retention time windows are listed below.

**List of evaluated metabolite standards including the retention time and measured accurate mass and the hereon based targeted masses and the defined time windows utilized as inclusion list for targeted selected ion monitoring (tSIM) measurements of *P. putida* extracts.**

| **metabolite** | $t_R$ /min | **exact mass** *m*/*z* | **accurate mass** *m*/*z* | **targeted mass** *m*/*z* | **tSIM-window** /min |
|---|---|---|---|---|---|
| pyruvate | 5.72 | 87.0077 | 87.0084 | 90.0084 | 4.50-12.00 |
| succinate | 11.74 | 117.0182 | 117.0193 | 120.0193 | 11.00-16.00 |
| malate | 11.73 | 133.0131 | 133.0134 | 136.0134 | 11.00-17.00 |
| fumarate | 13.79 | 115.0026 | 115.0037 | 118.0037 | 12.00-18.00 |
| citrate | 20.83 | 191.0186 | 191.0199 | 194.0189 | 20.00-26.00 |
| isocitrate | 21.74 | | | | |
| glucose 1-phosphate | 10.65 | 259.0213 | 259.0224 | 262.0224 | 10.00-18.00 |
| glucose 6-phosphate* | 13.23 | | | | |
| fructose 6-phosphate* | 13.23 | | | | |
| ribose 5-phosphate | 14.02 | 229.0108 | 229.0120 | 232.0120 | 11.00-18.00 |

| metabolite | $t_R$ /min | exact mass *m*/*z* | accurate mass *m*/*z* | targeted mass *m*/*z* | tSIM-window /min |
|---|---|---|---|---|---|
| 6-phosphogluconate | 17.75 | 275.0163 | 275.0175 | 278.0175 | 15.00-20.00 |
| | | 137.0039* | 137.0051* | 140.0051 | |
| 3-phosphoglyceric acid | 19.47 | 184.9846 | 184.9857 | 187.9857 | 18.00-24.00 |
| fructose 1,6-bisphosphate | 29.61 | 338.9877 | 338.9888 | 341.9888 | 28.50-35.00 |

*Glucose 6-phosphate and fructose 6-phosphate could not be separated.

For IC-MS instrument control, Xcalibur 4.1 software and the SII plugin (Thermo Scientific) was utilized. Metabolites were assigned by accurate mass (< 10 ppm relative mass deviation) and retention times by comparison to authentic standards. Identification and quantification of metabolites and the $^{13}C$-labelled isotopes was performed on basis of extracted ion chromatograms (relative mass deviation of ±10 ppm) of each isotope by peak area determination with Xcalibur QualBrowser software.

**Whole-genome sequencing**

*P. putida* EM42-derived strains were cultured in LB medium at 30°C till mid-exponential phase. Cells were then collected and enzymatically treated as previously described (Fišarová et al., 2021) with the following modification: achromopeptidase (1,000 U $mL^{-1}$; Sigma-Aldrich) and lysozyme (5 mg $mL^{-1}$; Sigma-Aldrich) were used instead of lysostaphin and mutanolysin. Genomic DNA was extracted using the Genomic DNA Clean & Concentrator-25 kit (Zymo Research) according to the manufacturer's instructions. For sequencing with the Oxford Nanopore platform, the library was prepared using the SQK-RAD004 Rapid Sequencing kit (Oxford Nanopore Technologies) according to the manufacturer's instructions. The library was sequenced with a FLO-FLG001 flow cell (R9.4.1) in a MinION device controlled by MinION software (release 22.05.5, Oxford Nanopore Technologies). Basecalling, demultiplexing, and barcode trimming were performed using standalone ONT Guppy software v6.1.7 using the config file dna_r9.4.1_450bps_sup.cfg with the default minimum q-score threshold of 10. For Illumina-based sequencing, a 500-bp sequencing library was prepared with xGen™ DNA Lib Prep EZ (Integrated DNA Technologies) according to the manufacturer's instructions. In brief, 500 ng total DNA was enzymatically fragmented for 8 minutes. CS_UMI Adapter (Integrated DNA Technologies) and dual indexing primers were used to adapt the library. The library was amplified for 8 PCR cycles. After QC and pooling, the library was sequenced using Illumina Nextseq instrument with 300 cycles mid output chemistry. Illumina reads were trimmed and filtered using Trimmomatic v0.38.1 with the sliding window model using a required average quality of 20 (Bolger et al., 2014). Complete bacterial genome sequences were obtained using a hybrid assembly with Unicycler v0.4.8 (Wick 2017) with minimal k-mer size of 0,2 and highest k-mer size of 0,95 with 10 k-mer steps used in SPAdes assembly. The resulting assembly was polished with Pilon v1.24 (Walker et al., 2014). Prokka v1.14.6 (https://doi.org/10.1093/bioinformatics/btu153) was used to annotate gene products for in-house proteomic analysis. The sequence of whole plasmid pSEVA2213_*xylABE* isolated from individual *P. putida* EM42 mutants was also verified separately by Plasmidsaurus service (USA) using Oxford Nanopore technology (https://www.plasmidsaurus.com). Genomic data were handled by Geneious Prime 2022.2.2 (Biomatters, Inc.): Mapping of sequencing reads was performed using Minimap2 (for Nanopore reads) and Bowtie (for Illumina reads) Plugins, and whole-genome alignment was performed using Mauve Plugin.

All sequencing data and assembled whole-genome sequences were deposited under NCBI BioProject PRJNA914626. The whole-genome sequences have been deposited in the GenBank database under accession numbers summarized in **Supplementary Table S3**. The genomes were annotated using the NCBI Prokaryotic Genome Annotation Pipeline (Tatusova et al., 2016). Only raw sequencing data have been deposited in SRA database (NCBI) for the following strains: PD584 ttrr L2, SRR22840560; PD584 tt L3, SRR22840561; EM42 Δ*gcd*, SRR22840563; PD310, SRR22840562.

#### Proteomic analyses

*P. putida* EM42 strains were pre-cultured overnight in 2.5 mL of LB medium with kanamycin. Cells were spun down (2,000 rpm, RT, 7 min), washed with M9 medium, and used for the inoculation of main cultures in 250 mL shake flasks with 50 mL of M9 medium, 2 g $L^{-1}$ xylose or glucose and kanamycin to a starting $OD_{600}$ of 0.05 (200 rpm, IS-971R, Jeio Tech). Cells (25 mL) were collected in the mid-exponential phase ($OD_{600}$=0.5) in pre-chilled falcon tubes (2,500 g, 4°C, 5 min), washed twice with ice-cold PBS buffer (per 1 L: 8 g NaCl, 0.2 g KCl, 1.44 g $Na_2HPO_4$, 0.24 g $KH_2PO_4$, pH 7.4) and pelleted in 1.5 mL Eppendorf tubes (15,000 g, 4°C, 2 min). Cell pellets were immediately frozen at -80°C and kept frozen until further use.

***Sample preparation and LC-MS analyses.*** Cell pellets were lysed using SDT buffer (4 % sodium dodecyl sulfate, 0.1 M dithiotreitol, 0.1 M Tris/HCl, pH 7.6) at 95°C for 30 min and the resulting protein solution was cleared by centrifugation at 20,000 g for 15 min. The protein lysates were processed by filter-aided sample preparation (FASP) (Wiśniewski et al., 2011) with some modifications as specified. The samples were mixed with 8 M UA buffer (8 M urea in 100 mM Tris-HCl, pH 8.5), loaded onto the Microcon device with MWCO 30 kDa (Merck Millipore), and centrifuged (7,000 g, 20°C, 30 min). The retained proteins were washed (all centrifugation steps after sample loading were performed at 14,000 g) with 200 µL UA buffer. The washed protein concentrates kept in the Microcon device were mixed with 100 µL of UA buffer containing 50 mM iodoacetamide and incubated in the dark for 20 min. After the next centrifugation step, the samples were washed three times with 100 µL of UA buffer and three times with 100 µL of 50 mM $NaHCO_3$. Trypsin (sequencing grade, Promega) was added onto the filter and the mixture was incubated for 18 h at 37°C (enzyme:protein ratio 1:100). The tryptic peptides were eluted by centrifugation followed by two additional elutions with 50 µL of 50 mM $NaHCO_3$. Peptides were then cleaned by liquid-liquid extraction (3 iterations) using water-saturated ethyl acetate (Yeung et al., 2008). Cleaned FASP eluate was evaporated completely in SpeedVac concentrator (Thermo Fisher Scientific). The resulting peptides were extracted into LC-MS vials by 2.5% formic acid (FA) in 50% acetonitrile (ACN) and 100% ACN with the addition of polyethylene glycol (20,000; final concentration 0.001%) (Stejskal et al., 2013) and concentrated in a SpeedVac concentrator (Thermo Fisher Scientific). LC-MS/MS analyses of all peptide mixtures were done using RSLCnano system connected to Orbitrap

Exploris 480 spectrometer (Thermo Fisher Scientific) with EASY Spray ion source (Thermo Fisher Scientific) installed. Prior to LC separation, tryptic digests were online concentrated and desalted using a trapping column (300 µm × 5 mm, µPrecolumn, 5 µm particles, Acclaim PepMap100 C18, Thermo Fisher Scientific). After washing the trapping column with 0.1% FA, the peptides were eluted (flow 300 nL $min^{-1}$) from the trapping column onto Acclaim PepMap RSLC C18 column (2 µm particles, 75 µm × 250 mm; Thermo Fisher Scientific) by 104 min long gradient. Mobile phase A (0.1% FA in water) and mobile phase B (0.1% FA in 80% acetonitrile) were used in both cases. The gradient elution started at 3% of mobile phase B and increased from 3% to 37% during the first 94 min, then increased linearly to 80% of mobile phase B in the next 7 min and remained at this state for the next 3 min. Equilibration of the trapping column and the column was done prior to sample injection to the sample loop. The analytical column outlet was directly connected to the Easy Spray ion source.

Data was acquired in a data-independent acquisition mode (DIA). The survey scan covered the *m*/*z* range of 350-1400 at a resolution of 60,000 (at *m*/*z* 200) and a maximum injection time of 55 ms. HCD MS/MS (27% relative fragmentation energy) was acquired in the range of *m*/*z* 200-2000 at 30,000 resolution (maximum injection time 55 ms). An overlapping windows scheme in *m*/*z* range from 400 to 800 was used as the isolation window placements.

***Analysis of proteomics data.*** DIA data were processed in DIA-NN version 1.8 (Demichev et al., 2020) in library-free mode against the modified cRAP database (based on http://www.thegpm.org/crap/; 112 sequences in total) and UniProtKB protein database for *Pseudomonas putida* (number of protein sequences: 310,975) and UniProtKB protein database for *Escherichia coli* (number of protein sequences: 4,402). No optional, carbamidomethylation as fixed modification and trypsin/P enzyme with 1 allowed missed cleavages and peptide length 7-30 were set during the library preparation. False discovery rate (FDR) control was set to 1%. MS1 and MS2 accuracies as well as scan window parameters were set based on the initial test searches (median value from all samples ascertained parameter values). MBR was switched on. Protein intensities reported in the DIA-NN main report file (**Supplementary File 6**) were further processed using the software container environment (https://github.com/OmicsWorkflows), version 4.1.3a. The processing workflow is available upon request. Briefly, it covered: a) removal of low-quality precursors and contaminant protein groups, b) protein group intensities log2 transformation, c) LoessF normalization, d) filtering out of protein groups not quantified in more than half of the replicates of at least one sample type (e) imputation of the missing values from the random distribution around the global minimal value, f) normalized and imputed protein intensities were used for differential expression using LIMMA statistical test and g) proteins with adjusted p-value <0.05 (P values adjustment on multiple hypothesis testing was done using Benjamini & Hochberg method) and log2 fold change >1 were used for Volcano plots.

Duplicities were removed in Excel. Gene identifiers were coupled to UniProt identifiers using the ID mapping tool available at uniprot.org. Data were visualized in Escher (escher.github.io) using a genome-scale model iJN1463 of *P. putida* KT2440.

### Supplementary results

#### *Deletion of hexR gene in P. putida EM42*

The purpose of the *hexR* deletion on the background of wild-type *P. putida* EM42 strain was to verify the connection between HexR and periplasmic oxidation of xylose to xylonate, which was observed in *P. putida* S12 (Meijnen et al., 2012). The *ΔhexR* mutant of S12 strain did not form the xylonate by-product while growing on xylose and the authors suggested that HexR may also control transcription of PPQ pyrroloquinoline quinone (a Gcd co-factor) biosynthesis genes in *P. putida*. The same effect was, nonetheless, not confirmed in *P. putida* EM42 Δ*hexR xylABE*$^+$. The mutant still oxidized ~40 % of xylose to xylonate and, consequently, its biomass yield was reduced (**Supplementary Fig. S3**).

### Supplementary tables

**Table S1.** Plasmids used in this study.

| Plasmid | Characterization | Source or reference |
|---|---|---|
| pRK600 | Helper plasmid for triparental mating: *oriV*(*ColE1*) RK2 *tra*$^+$ *mob*$^+$, Cm$^R$ | (Kessler et al., 1994) |
| pSEVA2213 | Expression vector: *oriV*(RK2) pEM7, Km$^R$ | (Silva-Rocha et al., 2013) |
| pBAMD1-4_P$_{EM7}$ | Mini-Tn5 delivery plasmid: *ori*(R6K) P$_{EM7}$, Amp$^R$ Sm$^R$/Sp$^R$ | (Martínez-García et al., 2014) |
| pEMG | Plasmid for genome editing in Gram-negative bacteria: *ori* (R6K), Km$^R$ | (Martínez-García and de Lorenzo, 2012) |
| pSW-I | Expression vector encoding I-SceI homing endonuclease from *Saccharomyces cerevisiae*: ori(RK2) *xylS-Pm*, Amp$^R$ | (Martínez-García and de Lorenzo, 2012) |
| pJNN_*phzA*1-*G*1, *phzM*, *phzS* | Expression vector bearing a pathway for pyocyanin production with respective regulatory sequences, Amp$^R$, Gm$^R$ | (Bator et al., 2020) |
| pEX_*rpe*-*rpiA* | Cloning vector with synthesized genes *rpe* of D-ribulose-5-phosphate 3-epimerase and *rpiA* of ribose 5-phosphate isomerase A, codon optimized for *P. putida* KT2440 and equipped with synthetic RBS, Amp$^R$ | This study (Eurofins Genomics) |
| pSEVA2213_*xylABE* | pSEVA2213 with synthetic *xylABE* operon encoding XylA xylose isomerase, XylB xylulokinase, XylE xylose-proton symporter from *Escherichia coli* (*Eco*RI/*Hind*III) | (Dvořák and de Lorenzo, 2018) |
| pEMG_*gcd* | pEMG with the cloned sequences (507+500 bp) spanning upstream and downstream of the *gcd* gene (PP1444) in *P. putida KT2440* genome (*Eco*RI/*Bam*HI) | (Dvořák and de Lorenzo, 2018) |
| pEMG_*hexR* | pEMG with the cloned sequences (500+500 bp) spanning upstream and downstream of the *hexR* gene (PP1021) in *P. putida KT2440* genome (*Eco*RI/*Bam*HI) | This study |
| pEMG_*gnd* | pEMG with the cloned sequences (503+502 bp) spanning upstream and downstream of the *gnd* gene (PP4043) in *P. putida KT2440* genome (*Sac*I/*Bam*HI) | Dr. Sánchez-Pascuala |
| pEMG_*pgi*-I | pEMG with the cloned sequences (592/650 bp) spanning upstream and downstream of the *pgi*-I gene (PP1808) in *P. putida KT2440* genome (*Eco*RI/*Bam*HI) | Dr. Sánchez-Pascuala |

| Plasmid | Characterization | Source or reference |
| --- | --- | --- |
| pEMG_*pgi*-II | pEMG with the cloned sequences (560+547 bp) spanning upstream and downstream of the *pgi*-II gene (PP4701) in *P. putida KT2440* genome (*Eco*RI/*Bam*HI) | Dr. Sánchez-Pascuala |
| pEMG_*edd* | pEMG with the cloned sequences (500+500 bp) spanning upstream and downstream of the *edd* gene (PP1010) in *P. putida KT2440* genome (*Eco*RI/*Bam*HI) | (Sánchez-Pascuala et al., 2019) |
| pBAMD1-4_EM7_*talB-tktA* | pBAMD1-4 _$P_{EM7}$ with a synthetic operon encoding transaldolase TalB and transketolase TktA from *E. coli* BL21(DE3) | This study |
| pBAMD1-4_EM7_*talB-tktA-rpe-rpiA* | pBAMD1-4 _$P_{EM7}$ with a synthetic operon encoding transaldolase TalB and transketolase TktA from *E. coli* BL21(DE3) and codon-optimized ,synthesized genes encoding D-ribulose-5-phosphate 3-epimerase Rpe and D-ribose-5-phosphate isomerase RpiA from *E. coli* BL21(DE3) | This study |

Abbreviations: $P_{EM7}$, constitutive promoter EM7; RBS, ribosom binding site; Amp, ampicillin; Cm, chloramphenicol; Gm, gentamicin; Km, kanamycin; Sm, streptomycin; Sp, spektinomycin.

**Table S2.** Oligonucleotide primers used in this study.

| Primer | Sequence (5'→3')[a] | Source |
|---|---|---|
| TS1F-gcd (EcoRI) | GGAATTCGCGGCAGTGCCGAGGTGTCGAAGTGGCGGTGG | (Dvořák and de Lorenzo, 2018) |
| TS1R-gcd | **GGCCTGAAGATCCAGAGCAGTTTCTAACCC**GCGACACCGCTCCCGCAGGCTCAACCCTGAGG | (Dvořák and de Lorenzo, 2018) |
| TS2F-gcd | GGGTTAGAAACTGCTCTGGATCTTCAGGCC | (Dvořák and de Lorenzo, 2018) |
| TS2R-gcd (BamHI) | CGGGATCCGTCAGCCGGCCGCCCTCAGCGGCGCCGCCT | (Dvořák and de Lorenzo, 2018) |
| gcd-check fw | CTTCAGCTCTTCGCTGTACA | (Dvořák and de Lorenzo, 2018) |
| gcd-check rv | GCGTGCGCTACAACCTTAC | (Dvořák and de Lorenzo, 2018) |
| TS1F-hexR (EcoRI) | ATTGAATTCGGCGGCTGGACCTGTGC | This study |
| TS1R-hexR | **CCCGACCCAAGGACACACCC**TCAACTGAGCCGGTGCG | This study |
| TS2F-hexR | GGGTGTGTCCTTGGGTCGGG | This study |
| TS2R-hexR (BamHI) | AATGGATCCGATAGCCCTCGGGCTGAAGG | This study |
| hexR PCR2 fw (EcoRI) | ATTGAATTCGGCGGCTGGACC | This study |
| hexR PCR2 rv (BamHI) | AATGGATCCGATAGCCCTCGG | This study |
| hexR-check fw | cGAACTCAAGCTGCAACTg | This study |
| hexR-check rv | gCATGTAGATGTCGGTATCttcc | This study |
| TS1F-gnd (SacI) | CGAGCTCCTGGCCAGGTCGGTGCCGAA | Dr. Sánchez-Pascuala |
| TS1R-gnd | **CGAAAAGGGAGCATTGGCTC**ATGACTAAACAGACCCTTGC | Dr. Sánchez-Pascuala |
| TS2F-gnd | GAGCCAATGCTCCCTTTTCG | Dr. Sánchez-Pascuala |
| TS2R-gnd (BamHI) | CGGGATCCGATGAAGGACAGCCGCTTGC | Dr. Sánchez-Pascuala |
| TS1F-pgi-I (EcoRI) | ATTTGAATTCACATCGACGACTTCCGCCAC | Dr. Sánchez-Pascuala |
| TS1R-pgi-I | AAGATCCTTGATACGGGTAAAGCCAG | Dr. Sánchez-Pascuala |
| TS2F-pgi-I | **CTGGCTTTACCCGTATCAAGGATCTT**CCCTGCTTGATACTGGCCCG | Dr. Sánchez-Pascuala |
| TS2R-pgi-I (BamHI) | ATTTGGATCCCTCAGAGGGGTCTCGGCAGG | Dr. Sánchez-Pascuala |

| Primer | Sequence (5’→3’)[a] | Source |
|---|---|---|
| TS1F-pgi-II (EcoRI) | ATTTGAATTCACCTGAATATGCCGATCCAGATCATC | Dr. Sánchez-Pascuala |
| TS1R-pgi-II | GTGCGTGGTGAGGGCTTGG | Dr. Sánchez-Pascuala |
| TS2F-pgi-II | **CCAAGCCCTCACCACGCAC**TCGCGGGTAAACCCGCCTAC | Dr. Sánchez-Pascuala |
| TS2R- pgi-II (BamHI) | ATTTGGATCCCGCCTTTCTTCACACCGCG | Dr. Sánchez-Pascuala |
| TS1F-edd (EcoRI) | GGAATTCGCACTGACCGCGATACGGTC | (Sánchez-Pascuala et al., 2019) |
| TS1R-edd | **CACCACCAGCAGGTGCTTCAT**GTACTGGACTCCAGGCTAATTG | (Sánchez-Pascuala et al., 2019) |
| TS2F-edd | ATGAAGCACCTGCTGGTTGGTG | (Sánchez-Pascuala et al., 2019) |
| TS2R-edd (BamHI) | CGGGATCCCCTACCGGCAGGTCAACATG | (Sánchez-Pascuala et al., 2019) |
| talB fw (SacI) | AATGAGCTCGCTGTTTAAAGAGAAATACTATCATGACG | This study |
| talB-PCR1 rv | TTACAGCAGATCGCCGATCA | This study |
| talB-PCR2 rv | **GCACGCCCTTAACGACTTG**TTACAGCAGATCGCCGATCA | This study |
| tktA fw | CAAGTCGTTAAGGGCGTGC | This study |
| tktA rv (SacI) | AATGAGCTCTTACAGCAGTTCTTTTGCTTTCG | This study |

Restriction sites are underlined, complementary sequences used in overlap extension (SOEing) PCR (Horton, 1990) are in bold.

**Table S3.** Accessions to sequencing data of *Pseudomonas strains* in this work in NCBI databases.

| BioProject PRJNA914626 | | | |
|---|---|---|---|
| Strain | BioSample Accession | GenBank Accession | SRA Accession[1] |
| PD584 | SAMN32340832 | CP115665 - chromosome<br>CP115666 - plasmid | SRR23032142<br>SRR23032143<br>SRR23032144 |
| PD584 L3 | SAMN32340833 | CP115663 - chromosome<br>CP115664 - plasmid | SRR23032140<br>SRR23032141 |
| PD689 tt L1 | SAMN32340834 | CP115661 - chromosome<br>CP115662 - plasmid | SRR23032137<br>SRR23032138<br>SRR23032139 |
| EM42del_gcd | SAMN32340957 | - | SRR22840563 |
| PD310 | SAMN32340958 | - | SRR22840562 |
| PD584 tt L3 | SAMN32340959 | - | SRR22840561 |
| PD584 ttrr L2 | SAMN32340960 | - | SRR22840560 |

[1] SRAs concern the whole genome, i.e. chromosomal and plasmid sequences.

**Table S4.** Presence of multiplied genomic locus PP_2114 - PP_2219 in the chromosome of sequenced *Pseudomonas putida* strains.

| | Coverage | | | | | | | | | | |
|---|---|---|---|---|---|---|---|---|---|---|---|
| | **Chromosome without multiplied region (A)** | | | | **Multiplied region (B)** | | | | **B/A ratio** | | |
| **Strain** | Illumina | SD | Nanopore | SD | Illumina | SD | Nanopore | SD | Illumina | Nanopore | **avg.** |
| **PD584** | 107.4 | 23.3 | 34.2 | 9.1 | 768.6 | 131.8 | 161.8 | 12.5 | 7.2 | 4.7 | **5.9** |
| **PD584 L3** | 56.2 | 14.2 | 27.4 | 8.8 | 386.7 | 69.4 | 148.6 | 14.5 | 6.9 | 5.4 | **6.2** |
| **PD689 tt L1** | 425.9 | 66.1 | 30.9 | 6.6 | 388.8 | 47.8 | 29.3 | 6.0 | 0.9 | 0.9 | **0.9** |
| **PD310** | N/A | N/A | 4.6 | 2.5 | N/A | N/A | 17.1 | 5.9 | N/A | 3.7 | **3.7** |
| **EM42 Δgcd** | N/A | N/A | 6.4 | 2.5 | N/A | N/A | 5.7 | 2.6 | N/A | 0.9 | **0.9** |

N/A - was not performed. Coverage was determined by Minimap2 plugin for Nanopore platform and Bowtie plugin for Illumina platform (Geneious Prime 2022.2.2).

### Supplementary figures

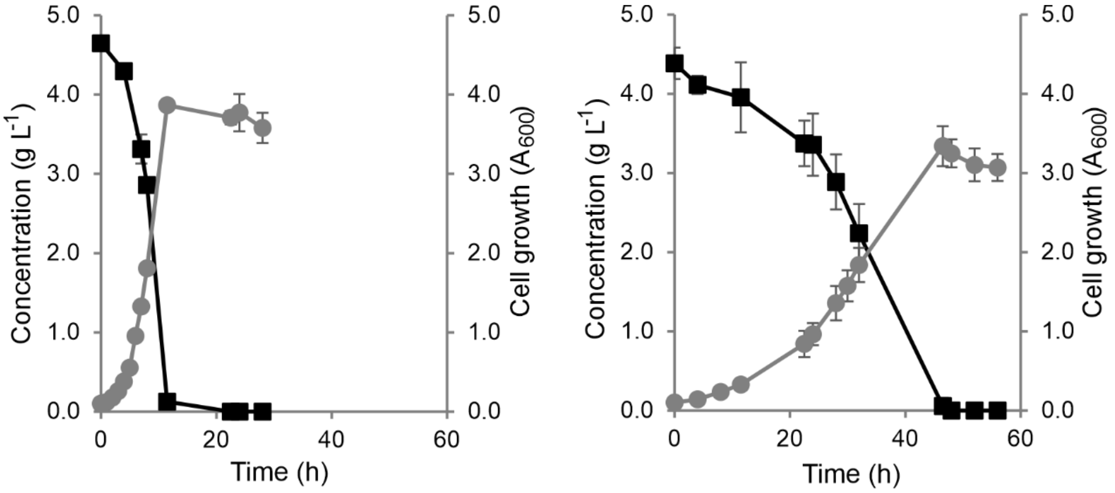


**Figure S1.** Shake flask cultures of *Pseudomonas putida* PD310 in M9 minimal medium with 5 g $L^{-1}$ glucose (left) or xylose (right). Cell growth, grey circles; glucose or xylose, black squares. Data points represent means ± standard deviations calculated from three biological replicates.

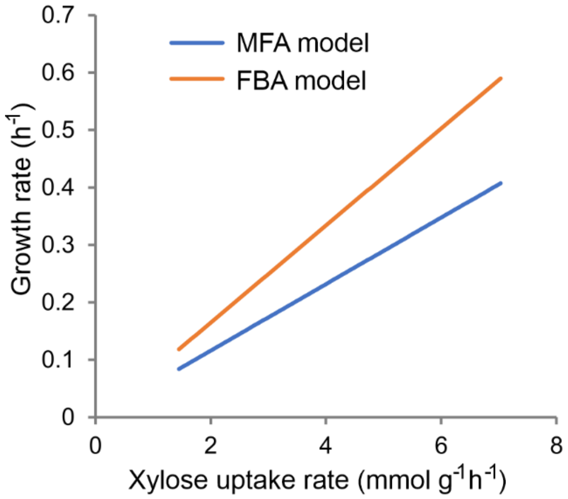


**Figure S2.** Comparison of growth rate and specific xylose uptake rate calculated using two metabolic models: MFA model (constrained with upper and lower bounds from experimental metabolic flux analysis of PD310 cells grown on labeled xylose) and FBA model (unconstrained model). Starting specific xylose uptake rate used in the calculation was $q_S$ = 1.45 mmol gCDW$^{-1}$ h$^{-1}$. The FBA model provides ≥40% higher growth rate.

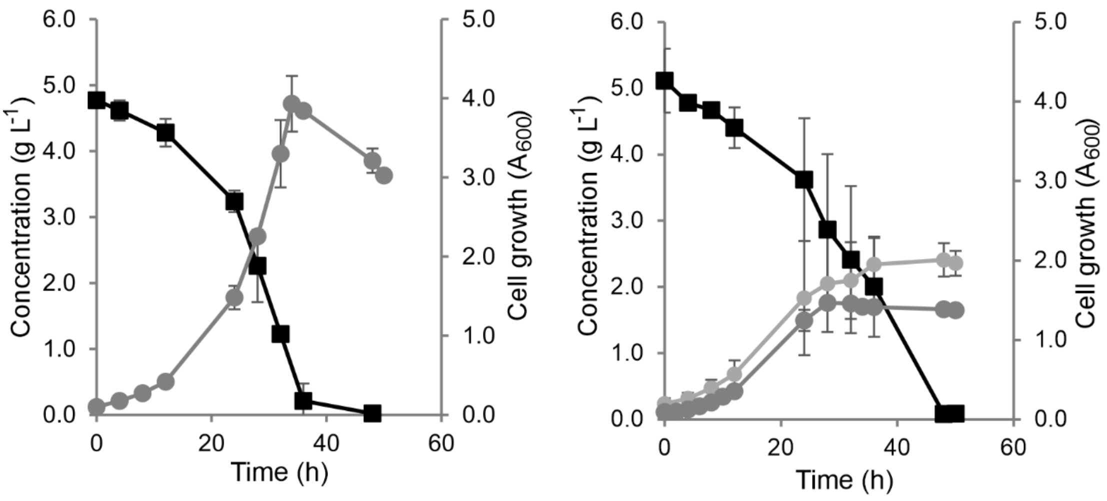


**Figure S3.** Shake flask cultures of *Pseudomonas putida* PD584 (left) or *P. putida* EM42 Δ*hexR* pSEVA2213_*xylABE* (right) in M9 minimal medium with xylose (5 g $L^{-1}$). Cell growth, dark grey circles; xylose, black squares; xylonic acid, pale grey circles. Data points represent means ± standard deviations calculated from three (PD584) or two (EM42 Δ*hexR* pSEVA2213_*xylABE*) biological replicates.

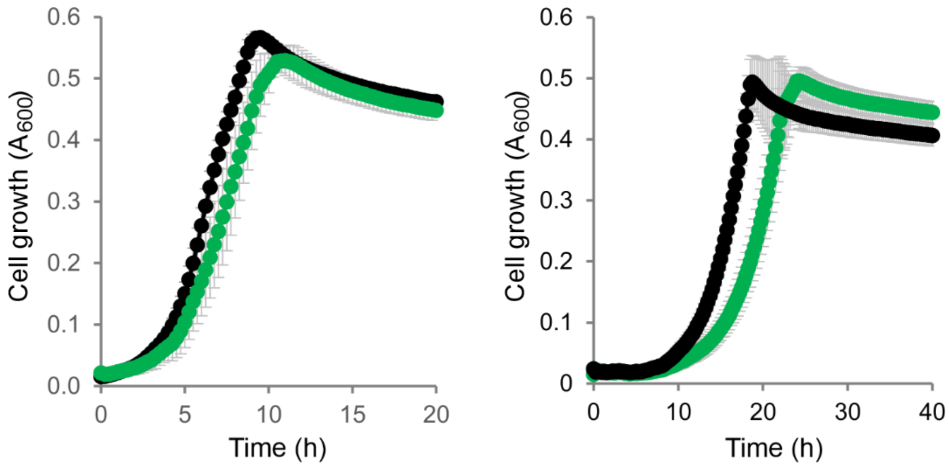


**Figure S4.** Growth of *hexR*$^+$ PD310 (green symbols) and *hexR*$^-$ PD584 (black symbols) in M9 minimal medium with 2 g $L^{-1}$ D-glucose (left graph) or 2 g $L^{-1}$ D-fructose (right graph) in 48-well microtiter plate. Data are shown as means ± standard deviations from three biological replicates. Determined growth parameters on (i) glucose: PD310 max. growth rate μ=0.47±0.00 $h^{-1}$, lag phase 1.5±0.2 h, PD584 μ=0.55±0.01 $h^{-1}$, lag phase 0.8±0.0 h; (ii) on fructose: PD310 μ=0.24±0.00 $h^{-1}$, lag phase 8.6±0.3 h, PD584 μ=0.30±0.00 $h^{-1}$, lag phase 6.7±0.2 h.

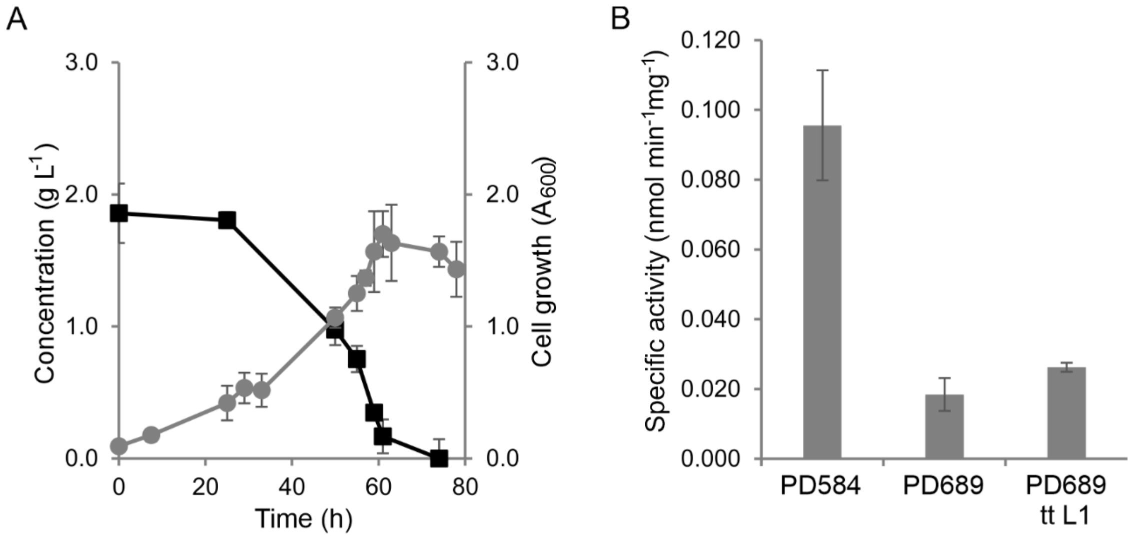


**Figure S5.** Shake flask cultures of *Pseudomonas putida* PD689 (**A**) in M9 minimal medium with xylose (2 g $L^{-1}$) and (**B**) specific activity of 6-phosphogluconate dehydrogenase Gnd determined in cell-free extracts of PD584, PD689, and PD689 tt L1. (**A**) Cell growth, dark grey circles; xylose, black squares. Data points represent means ± standard deviations calculated from three biological replicates. (**B**) Columns represent means ± standard deviations calculated from four biological replicates.

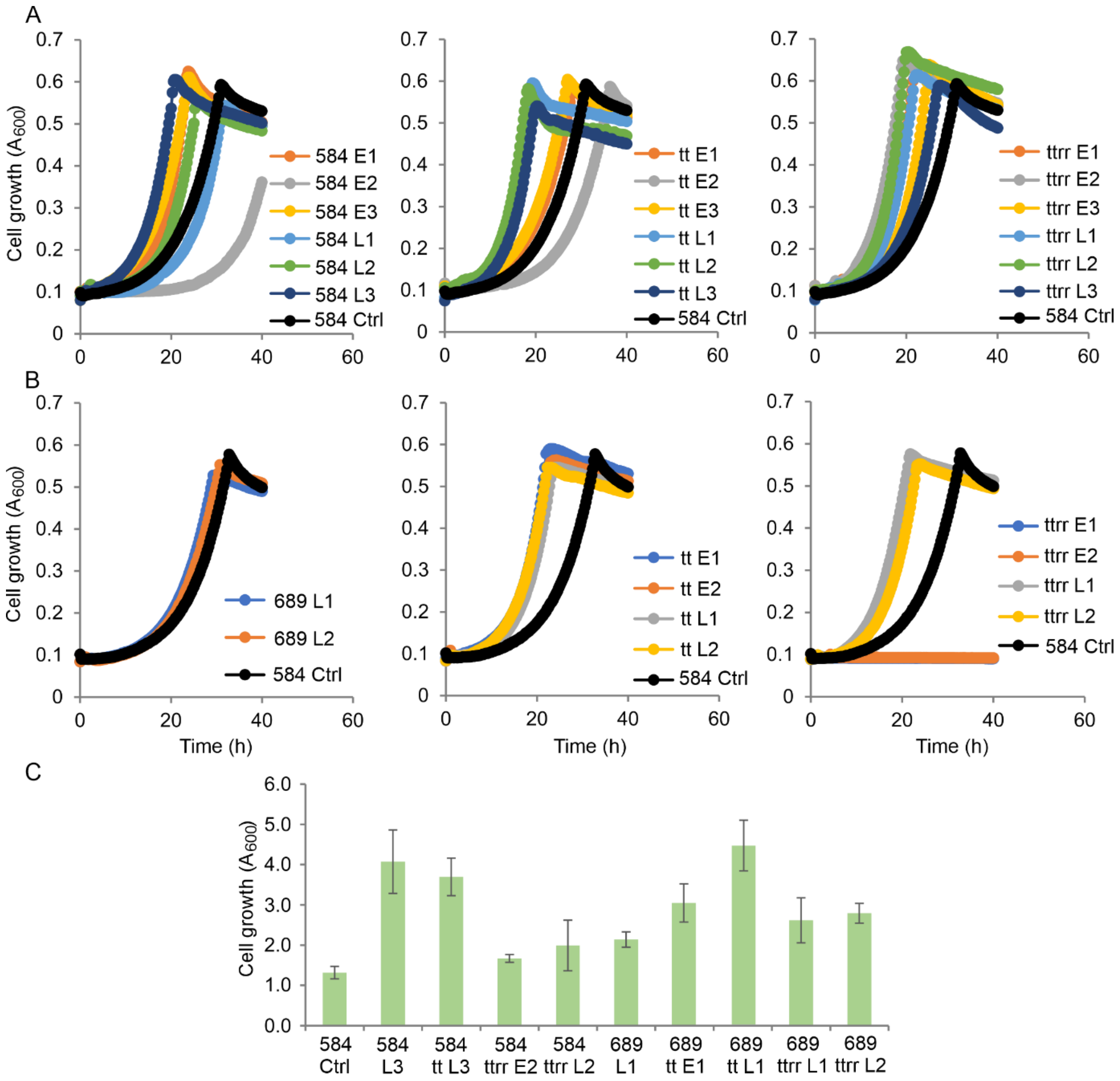


**Figure S6.** Screening of selected evolved mutants of *P. putida* PD584 and PD689 strains expressing heterologous pentose phosphate pathway genes on xylose in M9 minimal medium with 2 g $L^{-1}$ xylose in 48-well microtiter plate (**A** and **B**) or shake flasks with 5 g $L^{-1}$ xylose (**C**). E and L in the figure legends in **A** (PD584-derived candidates) and **B** (PD689-derived candidates) represent clones picked in the earlier phase of the ALE experiment when $OD_{600}$ of a given culture 24 h after inoculation reached the value of at least 3.50 for the first time (E), or clones picked at the end of the ALE experiment (L). Data in (**C**) are shown as means ± standard deviations from two independent experiments each of three biological replicates.

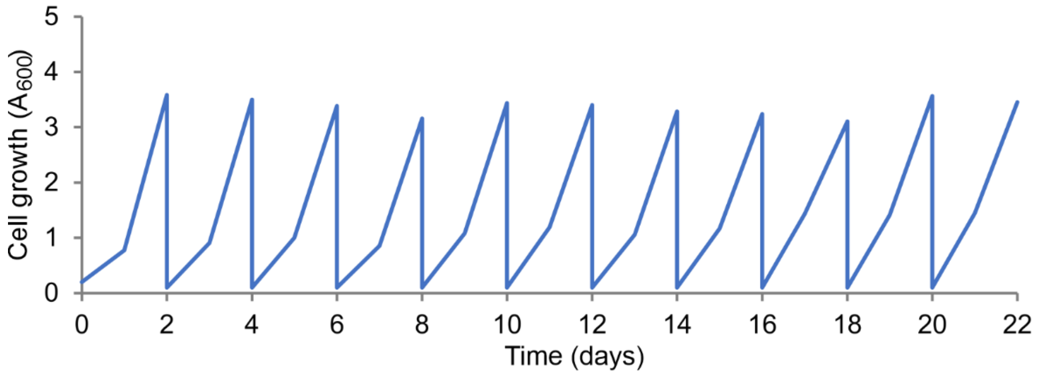


**Figure S7.** Adaptive laboratory evolution on xylose of *P. putida* PD310 strain cultured in shake flasks containing 20 mL of M9 minimal salts medium, 5 g L$^{-1}$ D-xylose and kanamycin and passaged every 48 h. Data points represent the means of two biological replicates.

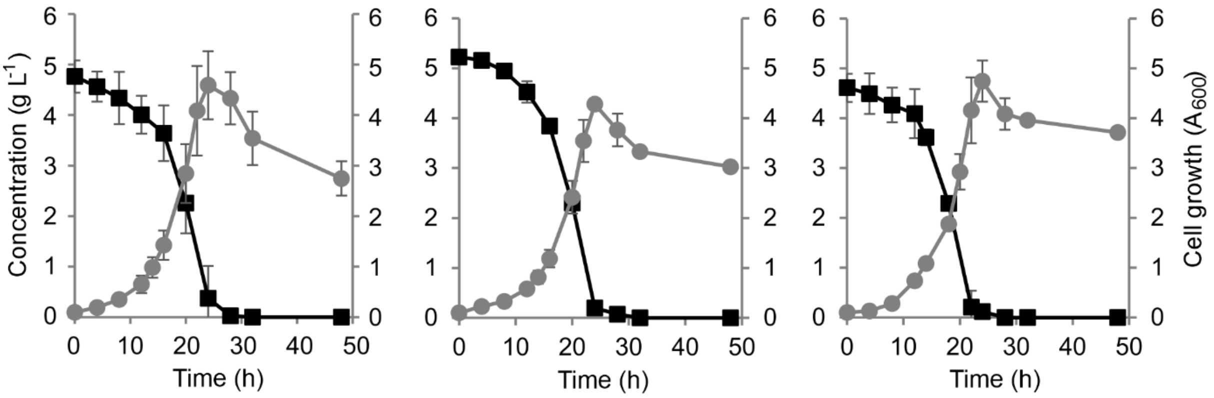


**Figure S8**. Shake flask cultures of *P. putida* PD584 L3 (left graph), PD584 tt L3 (central graph), and PD689 tt L1 (right graph) in M9 minimal medium with xylose (5 g L$^{-1}$). Cell growth, dark grey circles; xylose, black squares. Data points represent means ± standard deviations calculated from three biological replicates.

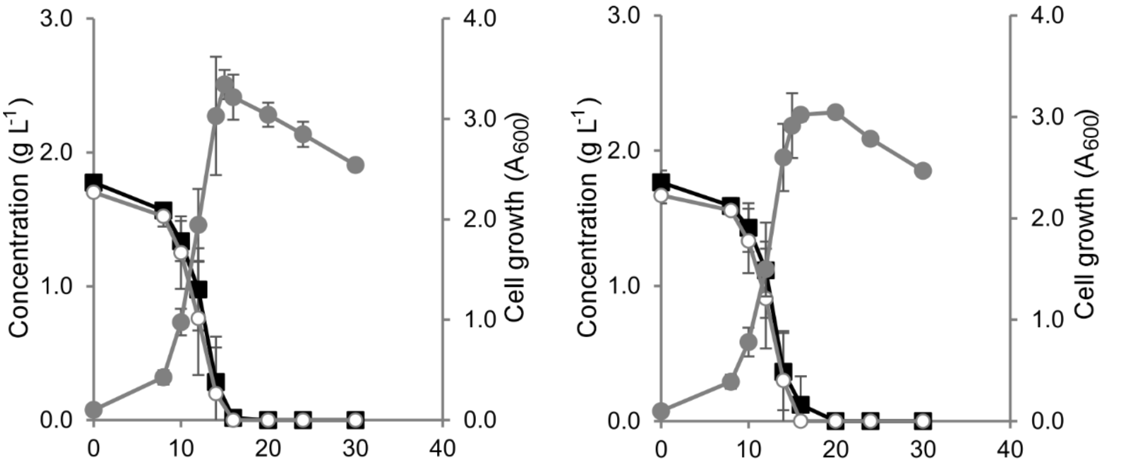


**Figure S9.** Shake flask cultures of *P. putida* PD584 L3 (left graph) and PD689 tt L1 (right graph) in M9 minimal medium with xylose and glucose (2 g $L^{-1}$ each). Cell growth, dark grey circles; xylose, black squares; glucose, white circles. Data points represent means ± standard deviations calculated from three biological replicates.

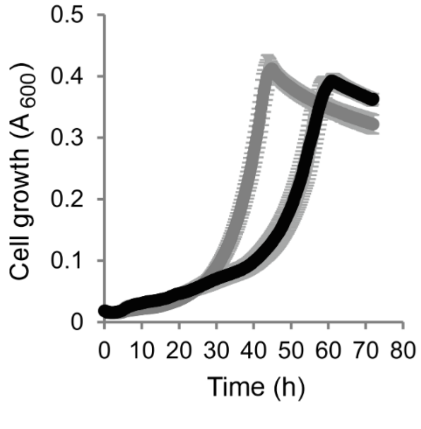


**Figure S10**. Growth of *P. putida* PD310 (grey symbols) and *P. putida* EM42 Δ*gcd* pSEVA2213_*xylABE* (black symbols) in 48-well microtiter plate with M9 minimal salts medium and 2 g $L^{-1}$ xylose. PD310 contains the multiplication of the ~118 kbp segment in its chromosome. Data for PD310 are shown as means ± standard deviations from three biological replicates each of two technical replicates. Data for *P. putida* EM42 Δ*gcd* pSEVA2213_*xylABE* are calculated as means ± standard deviations from cultures of two representative clones, each conducted in three biological replicates and two technical replicates.

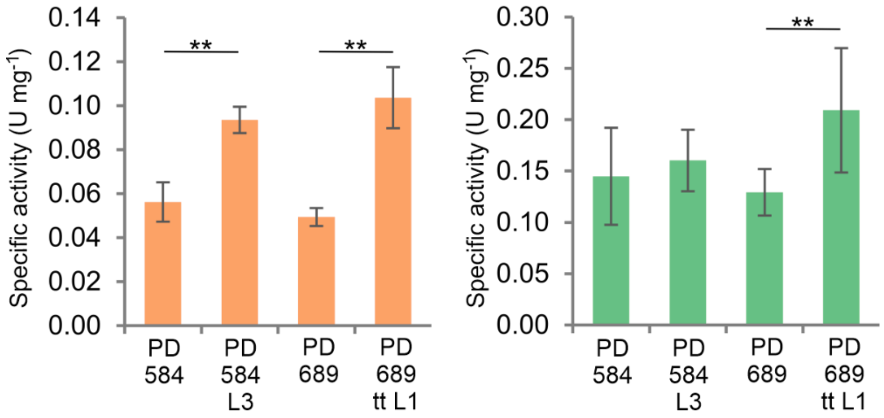


**Figure S11.** The specific activity of xylose isomerase XylA (left graph) and xylulokinase XylB (right graph) determined in cell-free extracts of *P. putida* strains. Columns represent means ± standard deviations calculated from two independent experiments, each of four biological replicates. Asterisks denote the significance of the difference between the two means at $p < 0.01$.

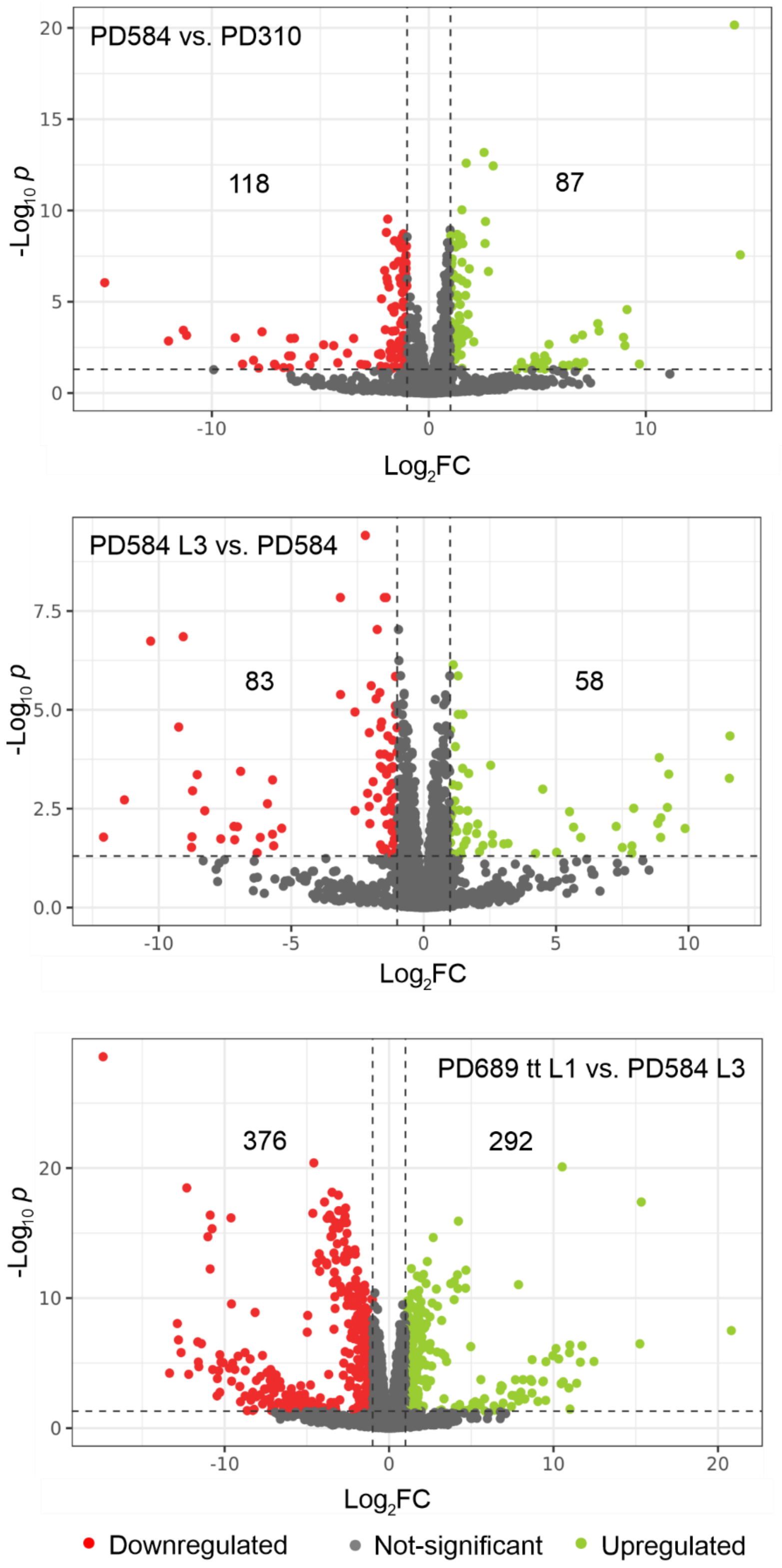


**Figure S12.** Volcano plots depicting differentially abundant proteins in strain PD584 compared to PD310, PD584 L3 compared to PD584, and PD689 tt L1 compared to PD584 L3. The strains were grown in M9 minimal salt medium with 2 g $L^{-1}$ and harvested in the mid-exponential phase. The biomass was further processed as described in the Materials and methods section in the main body of the article. Significantly downregulated proteins ($p < 0.05$, $\log_2$ fold change <-1.0) are shown as red dots, significantly upregulated proteins ($p < 0.05$, $\log_2$ fold change >1.0) are shown as green dots. Each point in the plot represents an individual protein. Upregulated and downregulated proteins from individual comparisons are listed in **Supplementary File S7**.

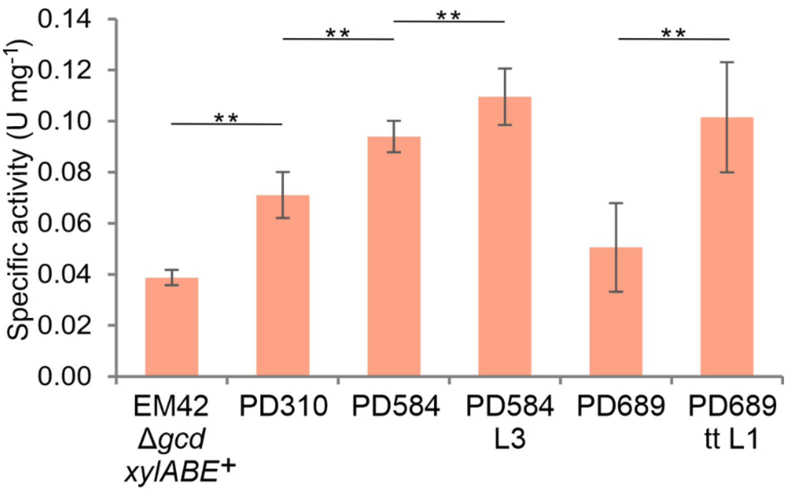


**Figure S13.** The specific activity of transaldolase determined in cell-free extracts of six *P. putida* strains. Columns represent means ± standard deviations calculated from two independent experiments, each of four biological replicates. Asterisks denote the significance of the difference between the two means at $p < 0.01$.

**Supplementary sequences**

Expression cassette (4348 bp) with *talB* and *tktA* genes from *E. coli* BL21(DE3)

Includes (in the following order): mosaic element I (ME-I), EM7 promoter, *talB* gene with its native RBS, *tktA* gene with its native RBS, T500 terminator, *aadA* gene for aminoglycoside (3'') (9) adenylyltransferase conferring resistance to spectinomycin and streptomycin, mosaic element O (ME-O).

```
ctgtctcttatacacatctttgtgtctcaggccgcctaggttgttgacaattaatcatcggc
atagtatatcggcatagtataatacgacaaggtgaggaactaaacccgtaggccgccgcgcg
cgaattcgagctcgctgtttaaagagaaatactatcatgacggacaaattgacctcccttcg
tcagtacaccaccgtagtggccgacactggggacatcgcggcaatgaagctgtatcaaccgc
aggatgccacaaccaacccttctctcattcttaacgcagcgcagattccggaataccgtaag
ttgattgatgatgctgtcgcctgggcgaaacagcagagcaacgatcgcgcgcagcagatcgt
ggacgcgaccgacaaactggcagtaaatattggtctggaaatcctgaaactggttccgggcc
gtatctcaactgaagttgatgcgcgtctttcctatgacaccgaagcgtcaattgcgaaagca
aaacgcctgatcaaactctacaacgatgctggtattagcaacgatcgtattctgatcaaact
ggcttctacctggcagggtatccgtgctgcagaacagctggaaaaagaaggcatcaactgta
acctgaccctgctgttctccttcgctcaggctcgtgcttgtgcggaagcgggcgtgttcctg
atctcgccgtttgttggccgtattcttgactggtacaaagcgaataccgataagaaagagta
cgctccggcagaagatccgggcgtggtttctgtatctgaaatctaccagtactacaaagagc
acggttatgaaaccgtggttatgggcgcaagcttccgtaacatcggcgaaattctggaactg
gcaggctgcgaccgtctgaccatcgcaccggcactgctgaaagagctggcggagagcgaagg
ggctatcgaacgtaaactgtcttacaccggcgaagtgaaagcgcgtccggcgcgtatcactg
agtccgagttcctgtggcagcacaaccaggatccaatggcagtagataaactggcggaaggt
atccgtaagtttgctattgaccaggaaaaactggaaaaaatgatcggcgatctgctgtaaca
agtcgttaagggcgtgcccttcatcatccgatctggagtcaaaatgtcctcacgtaaagagc
ttgccaatgctattcgtgcgctgagcatggacgcagtacagaaagccaaatccggtcacccg
ggtgcccctatgggtatggctgacattgccgaagtcctgtggcgtgatttcctgaaacacaa
cccgcagaatccgtcctgggctgaccgtgaccgcttcgtgctgtccaacggccacggctcca
tgctgatctacagcctgctgcacctcaccggttacgatctgccgatggaagaactgaaaaac
ttccgtcagctgcactctaaaactccgggtcacccggaagtgggttacaccgctggtgtgga
aaccaccaccggtccgctgggtcagggtattgccaacgcagtcggtatggcgattgcagaaa
aaacgctggcggcgcagtttaaccgtccgggccacgacattgtcgaccactacacctacgcc
ttcatgggcgacggctgcatgatggaaggcatctcccacgaagtttgctctctggcgggtac
gctgaagctgggtaaactgattgcattctacgatgacaacggtatttctatcgatggtcacg
ttgaaggctggttcaccgacgacaccgcaatgcgtttcgaagcttacggctggcacgttatt
cgcgacatcgacggtcatgacgcggcatctatcaaacgcgcagtagaagaagcgcgcgcagt
gactgacaaaccttccctgctgatgtgcaaaaccatcatcggtttcggttccccgaacaaag
ccggtacccacgactcccacggtgcgccgctgggcgacgctgaaattgccctgacccgcgaa
caactgggctggaaatatgcgccgttcgaaatcccgtctgaaatctatgctcagtgggatgc
gaaagaagcaggccaggcgaaagaatccgcatggaacgagaaattcgctgcttacgcgaaag
cttatccgcaggaagccgctgaatttacccgccgtatgaaaggcgaaatgccgtctgacttc
gacgctaaagcgaaagagttcatcgctaaactgcaggctaatccggcgaaaatcgccagccg
taaagcgtctcagaatgctatcgaagcgttcggtccgctgttgccggaattcctcggcggtt
ctgctgacctggcgccgtctaacctgaccctgtggtctggttctaaagcaatcaacgaagat
gctgcgggtaactacatccactacggtgttcgcgagttcggtatgaccgcgattgctaacgg
tatctccctgcacggtggcttcctgccgtacacctccaccttcctgatgttcgtggaatacg
```

cacgtaacgccgtacgtatggctgcgctgatgaaacagcgtcaggtgatggtttacacccac
gactccatcggtctgggcgaagacggcccgactcaccagccggttgagcaggtcgcttctct
gcgcgtaaccccgaacatgtctacatggcgtccgtgtgaccaggttgaatccgcggtcgcgt
ggaaatacggtgttgagcgtcaggacggcccgaccgcactgatcctctcccgtcagaacctg
gcgcagcaggaacgaactgaagagcaactggcaaacatcgcgcgcggtggttatgtgctgaa
agactgcgccggtcagccggaactgattttcatcgctaccggttcagaagttgaactggctg
ttgctgcctacgaaaaactgactgccgaaggcgtgaaagcgcgcgtggtgtccatgccgtct
accgacgcatttgacaagcaggatgctgcttaccgtgaatccgtactgccgaaagcggttac
tgcacgcgttgctgtagaagcgggtattgctgactactggtacaagtatgttggcctgaacg
gtgctatcgtcggtatgaccaccttcggtgaatctgctccggcagagctgctgtttgaagag
ttcggcttcactgttgataacgttgttgcgaaagcaaaagaactgctgtaagagctcggtac
ccggggatcctctagagtcgacctgcaggcatgcaagcttgcggccgccaaagcccgccgaa
aggcgggcttttctgtatttaaatgaaccttgaccgaacgcagcggtggtaacggcgcagtg
gcggttttcatggcttgttatgactgtttttttggggtacagtctatgcctcgggcatccaa
gcagcaagcgcgttacgccgtgggtcgatgtttgatgttatggagcagcaacgatgttacgc
agcagggcagtcgccctaaaacaaagttaaacatcatgagggaagcggtgatcgccgaagta
tcgactcaactatcagaggtagttggcgtcatcgagcgccatctcgaaccgacgttgctggc
cgtacatttgtacggctccgcagtggatggcggcctgaagccacacagtgatattgatttgc
tggttacggtgaccgtaaggcttgatgaaacaacgcggcgagctttgatcaacgaccttttg
gaaacttcggcttcccctggagagagcgagattctccgcgctgtagaagtcaccattgttgt
gcacgacgacatcattccgtggcgttatccagctaagcgcgaactgcaatttggagaatggc
agcgcaatgacattcttgcaggtatcttcgagccagccacgatcgacattgatctggctatc
ttgctgacaaaagcaagagaacatagcgttgccttggtaggtccagcggcggaggaactctt
tgatccggttcctgaacaggatctatttgaggcgctaaatgaaaccttaacgctatggaact
cgccgcccgactgggctggcgatgagcgaaatgtagtgcttacgttgtcccgcatttggtac
agcgcagtaaccggcaaaatcgcgccgaaggatgtcgctgccgactgggcaatggagcgcct
gccggcccagtatcagcccgtcatacttgaagctagacaggcttatcttggacaagaagaag
atcgcttggcctcgcgcgcagatcagttggaagaatttgtccactacgtgaaaggcgagatc
accaaggtagtcggcaaataagacaattgtctaattaattgcggaccctagaggtcccctttt
tttattttaaaatttttcacaaaacggtttacaagcataaaatctctgaagatgtgtata
agagacag

Expression cassette (5727 bp) with *talB*, *tktA*, *rpe*, and *rpiA* genes from *E. coli* BL21(DE3)

Includes (in the following order): mosaic element I (ME-I), EM7 promoter, *talB* gene with its native RBS, *tktA* gene with its native RBS, *rpe* gene with synthetic RBS, *rpiA* gene with synthetic RBS, T500 terminator, *aadA* gene for aminoglycoside (3'') (9) adenylyltransferase conferring resistance to spectinomycin and streptomycin, mosaic element O (ME-O).

```
ctgtctcttatacacatctttgtgtctcaggccgcctaggttgttgacaattaatcatcggc
atagtatatcggcatagtataatacgacaaggtgaggaactaaacccgtaggccgccgcgcg
cgaattcgagctcgctgtttaaagagaaatactatcatgacggacaaattgacctcccttcg
tcagtacaccaccgtagtggccgacactggggacatcgcggcaatgaagctgtatcaaccgc
aggatgccacaaccaacccttctctcattcttaacgcagcgcagattccggaataccgtaag
ttgattgatgatgctgtcgcctgggcgaaacagcagagcaacgatcgcgcgcagcagatcgt
ggacgcgaccgacaaactggcagtaaatattggtctggaaatcctgaaactggttccgggcc
gtatctcaactgaagttgatgcgcgtctttcctatgacaccgaagcgtcaattgcgaaagca
aaacgcctgatcaaactctacaacgatgctggtattagcaacgatcgtattctgatcaaact
ggcttctacctggcagggtatccgtgctgcagaacagctggaaaaagaaggcatcaactgta
acctgaccctgctgttctccttcgctcaggctcgtgcttgtgcggaagcgggcgtgttcctg
atctcgccgtttgttggccgtattcttgactggtacaaagcgaataccgataagaaagagta
cgctccggcagaagatccgggcgtggtttctgtatctgaaatctaccagtactacaaagagc
acggttatgaaaccgtggttatgggcgcaagcttccgtaacatcggcgaaattctggaactg
gcaggctgcgaccgtctgaccatcgcaccggcactgctgaaagagctggcggagagcgaagg
ggctatcgaacgtaaactgtcttacaccggcgaagtgaaagcgcgtccggcgcgtatcactg
agtccgagttcctgtggcagcacaaccaggatccaatggcagtagataaactggcggaaggt
atccgtaagtttgctattgaccaggaaaaactggaaaaaatgatcggcgatctgctgtaaca
agtcgttaagggcgtgcccttcatcatccgatctggagtcaaaatgtcctcacgtaaagagc
ttgccaatgctattcgtgcgctgagcatggacgcagtacagaaagccaaatccggtcacccg
ggtgcccctatgggtatggctgacattgccgaagtcctgtggcgtgatttcctgaaacacaa
cccgcagaatccgtcctgggctgaccgtgaccgcttcgtgctgtccaacggccacggctcca
tgctgatctacagcctgctgcacctcaccggttacgatctgccgatggaagaactgaaaaac
ttccgtcagctgcactctaaaactccgggtcacccggaagtgggttacaccgctggtgtgga
aaccaccaccggtccgctgggtcagggtattgccaacgcagtcggtatggcgattgcagaaa
aaacgctggcggcgcagtttaaccgtccgggccacgacattgtcgaccactacacctacgcc
ttcatgggcgacggctgcatgatggaaggcatctcccacgaagtttgctctctggcgggtac
gctgaagctgggtaaactgattgcattctacgatgacaacggtatttctatcgatggtcacg
ttgaaggctggttcaccgacgacaccgcaatgcgtttcgaagcttacggctggcacgttatt
cgcgacatcgacggtcatgacgcggcatctatcaaacgcgcagtagaagaagcgcgcgcagt
gactgacaaaccttccctgctgatgtgcaaaaccatcatcggtttcggttccccgaacaaag
ccggtacccacgactcccacggtgcgccgctgggcgacgctgaaattgccctgacccgcgaa
caactgggctggaaatatgcgccgttcgaaatcccgtctgaaatctatgctcagtgggatgc
gaaagaagcaggccaggcgaaagaatccgcatggaacgagaaattcgctgcttacgcgaaag
cttatccgcaggaagccgctgaatttacccgccgtatgaaaggcgaaatgccgtctgacttc
gacgctaaagcgaaagagttcatcgctaaactgcaggctaatccggcgaaaatcgccagccg
taaagcgtctcagaatgctatcgaagcgttcggtccgctgttgccggaattcctcggcggtt
ctgctgacctggcgccgtctaacctgaccctgtggtctggttctaaagcaatcaacgaagat
gctgcgggtaactacatccactacggtgttcgcgagttcggtatgaccgcgattgctaacgg
tatctccctgcacggtggcttcctgccgtacacctccaccttcctgatgttcgtggaatacg
cacgtaacgccgtacgtatggctgcgctgatgaaacagcgtcaggtgatggtttacacccac
```

gactccatcggtctgggcgaagacggcccgactcaccagccggttgagcaggtcgcttctct
gcgcgtaaccccgaacatgtctacatggcgtccgtgtgaccaggttgaatccgcggtcgcgt
ggaaatacggtgttgagcgtcaggacggcccgaccgcactgatcctctcccgtcagaacctg
gcgcagcaggaacgaactgaagagcaactggcaaacatcgcgcgcggtggttatgtgctgaa
agactgcgccggtcagccggaactgattttcatcgctaccggttcagaagttgaactggctg
ttgctgcctacgaaaaactgactgccgaaggcgtgaaagcgcgcgtggtgtccatgccgtct
accgacgcatttgacaagcaggatgctgcttaccgtgaatccgtactgccgaaagcggttac
tgcacgcgttgctgtagaagcgggtattgctgactactggtacaagtatgttggcctgaacg
gtgctatcgtcggtatgaccaccttcggtgaatctgctccggcagagctgctgtttgaagag
ttcggcttcactgttgataacgttgttgcgaaagcaaaagaactgctgtaagagctcggtac
ccggggatcctctagagtcgacctgcaggcatgccaattaaagggaggtccaaatgaagcag
tacctgatcgccccaagcatcctgagcgccgacttcgcccgtctgggcgaagataccgccaa
agcgctggccgccggtgccgacgtggtgcacttcgacgtgatggacaaccactacgtgccga
acctgaccatcggtcccatggtgctgaagtcgctgcgcaactacggcatcaccgcgccaatc
gacgtgcacctgatggtgaagccggtggaccgcatcgtgccggatttcgccgccgcgggtgc
cagcatcatcaccttccatccggaagccagcgagcacgtggatcgcaccctgcaactgatca
aagagaacggctgcaaagccggtctggtgttcaacccagcgaccccactgagctacctggac
tacgtcatggacaagctggacgtgatcctgctgatgagcgtgaacccaggcttcggtggcca
gagcttcatcccgcagaccctggataagctgcgcgaggtgcgtcgccgtatcgacgagagcg
gcttcgacatccgcctggaagtggacggtggcgtgaaggtgaacaacatcggcgagatcgcc
gcggccggtgcggacatgttcgtggccggtagcgccatcttcgaccagccggactacaagaa
agtgatcgacgaaatgcgctcggagctggccaaggtgagccacgagtgaggtacctgagctg
gaaatattaatgacccaggacgagctgaagaaggccgtcggctgggccgcgctgcaatacgt
gcaaccaggcaccatcgtcggtgtcggcaccggtagcaccgccgcgcacttcatcgacgccc
tgggcaccatgaagggccagatcgagggtgccgtgagcagcagcgacgccagcaccgagaag
ctgaaaagcctgggcatccacgtgttcgacctgaacgaggtggactcgctgggcatctatgt
ggatggtgcggacgagatcaacggccacatgcagatgatcaaaggcggtggtgccgcgctga
cccgtgagaagatcatcgccagcgtggccgagaagttcatctgcatcgccgacgcctcgaaa
caggtggatatcctgggcaagttcccgctgccggtggaagtgatcccgatggcgcgtagcgc
cgtggcgcgtcagctggtgaaactgggtggccgtccggaataccgccagggtgtcgtgaccg
acaacggcaacgtcatcctggacgtgcacggcatggaaatcctggacccgatcgccatggaa
aacgccatcaacgcgatcccaggcgtggtgaccgtgggcctgttcgccaaccgtggtgccga
tgtggccctgatcggcaccccagacggcgtcaagaccatcgtgaagtgaggatccgcggccg
ccaaagcccgccgaaaggcgggcttttctgtatttaaatgaaccttgaccgaacgcagcggt
ggtaacggcgcagtggcggttttcatggcttgttatgactgtttttttggggtacagtctat
gcctcgggcatccaagcagcaagcgcgttacgccgtgggtcgatgtttgatgttatggagca
gcaacgatgttacgcagcagggcagtcgccctaaaacaaagttaaacatcatgagggaagcg
gtgatcgccgaagtatcgactcaactatcagaggtagttggcgtcatcgagcgccatctcga
accgacgttgctggccgtacatttgtacggctccgcagtggatggcggcctgaagccacaca
gtgatattgatttgctggttacggtgaccgtaaggcttgatgaaacaacgcggcgagctttg
atcaacgaccttttggaaacttcggcttcccctggagagagcgagattctccgcgctgtaga
agtcaccattgttgtgcacgacgacatcattccgtggcgttatccagctaagcgcgaactgc
aatttggagaatggcagcgcaatgacattcttgcaggtatcttcgagccagccacgatcgac
attgatctggctatcttgctgacaaaagcaagagaacatagcgttgccttggtaggtccagc
ggcggaggaactctttgatccggttcctgaacaggatctatttgaggcgctaaatgaaacct
taacgctatggaactcgccgcccgactgggctggcgatgagcgaaatgtagtgcttacgttg
tcccgcatttggtacagcgcagtaaccggcaaaatcgcgccgaaggatgtcgctgccgactg

ggcaatggagcgcctgccggcccagtatcagcccgtcatacttgaagctagacaggcttatc
ttggacaagaagaagatcgcttggcctcgcgcgcagatcagttggaagaatttgtccactac
gtgaaaggcgagatcaccaaggtagtcggcaaataagacaattgtctaattaattgcggacc
ctagaggtcccctttttattttaaaatttttcacaaaacggtttacaagcataaaatct
ctgaagatgtgtataagagacag
